# Dense Sampling of Taxa and Genomes Untangles the Phylogenetic Backbone of a Non-model Plant Lineage Rife with Deep Hybridization and Allopolyploidy

**DOI:** 10.1101/2023.10.21.563444

**Authors:** Chao Xu, Zetao Jin, Hui Wang, Siyu Xie, Xiaohua Lin, Richard G.J. Hodel, Yu Zhang, Daikun Ma, Bing Liu, Guangning Liu, Shuihu Jin, Liang Zhao, Jun Wu, Chen Ren, Deyuan Hong, Binbin Liu

## Abstract

Phylogenetic networks, rather than purely bifurcating trees, more accurately depict the intricate evolutionary dynamics of most lineages, especially those characterized by extensive hybridization and allopolyploidization events. However, the challenges of achieving complete taxon sampling, and limited financial resources for studying non-model plant lineages, have hindered comprehensive and robust estimation of phylogenetic backbones with guidance from networks. The bellflower tribe, Campanuleae, characterized by a reticulate evolutionary history, serves as an ideal model to investigate how to diagnose nested ancient reticulation events. Here, by integrating multiple genomic data sources and a range of phylogenetic inference methods, we produced a robust phylogenetic backbone for the tribe Campanuleae. Our investigation of reticulate evolution indicates that hybridization and allopolyploidization were instrumental in shaping the diversity of the bellflower tribe, particularly during the initial diversification of the subtribe Phytematinae. Additionally, we ascertained that conflicting topologies resulting from distinct genomic datasets and inference methodologies significantly impact downstream estimates of divergence dating, ancestral area construction, and diversification rates. This study offers a universally relevant framework for deciphering how to use network-based phylogenetic structures using various genomic sources and inference methods. [Campanulaceae, Campanuleae, Cytonuclear discordance, paralog, phylogenomics, reticulate evolution]

---

Building on the legacy of Darwin’s Origin of Species (Darwin 1859), the Tree of Life (ToL) has been used as a model and a research tool to explore the evolution and relationships between living and extinct organisms with an assumption of bifurcating phylogeny (Mindell 2013). Given the prevalence of reticulation via incomplete lineage sorting (ILS), hybridization, polyploidization, and introgression, modeling the evolutionary connectivity of all life using a bifurcating phylogeny is problematic biologically and unrealistic (Rothfels 2021; Stull et al. 2023). A growing body of genomic and/or phylogenomic studies have provided mounting evidence supporting a network-like structure of life, such as the Bacteria and Archaea lineages (Dagan and Martin 2009; Gontier 2015), the vascular plant lineages (Leebens-Mack et al. 2019; Stull et al. 2021), birds (Jarvis et al. 2014), and mammals (Upham et al. 2019). Over the past decades, improved bioinformatic methods have been developed for teasing apart the various mechanisms underlying complex reticulate evolutionary histories, and multiple software programs have also been developed for resolving the same process, e.g., PhyloNet (Wen et al. 2018) and SNaQ (Solís-Lemus et al. 2017). However, the lack of a standardized procedure for untangling weblike relationships has impeded our understanding of evolutionary patterns and phylogenetic relationships.

In contrast to model plants such as *Arabidopsis thaliana* in Brassicaceae, rice (*Oryza sativa*) and maize (*Zea mays*) in Poaceae, and the tobacco plant (*Nicotiana tabacum*) in Solanaceae, non-model plants represent the vast majority of plant diversity on Earth, and most of them have significant ecological, agricultural, or medicinal importance. Studying non-model plants is critical for gaining a broader understanding of plant biology, evolution, and adaptation, especially given the tremendous diversity of plants in nature. However, the understudied background knowledge, cost and resource constraints, and unavailable biological samplings greatly challenged the phylogenomic studies of non-model plants. Critically, when species diversity in non-model lineages is not thoroughly sampled using multiple sources of genomic data, it can be impossible to tease apart ancient evolutionarily significant events such as hybridization. The decreased High-Throughput Sequencing (HTS) cost, especially in China (Liu et al. 2021), promoted the genome-level sequencing of non-model plants. It is important to investigate in depth a non-model plant lineage characterized by pervasive hybridization and allopolyploidy, and employ a multi-source genomic approach to explore its phylogenetic backbone and deep reticulation.

The Campanuleae tribe, the largest lineage in the Campanulaceae family with over 620 species, has undergone extensive hybridization and polyploidy events, as noted in previous studies (e.g., Lammers 2007a, 2007b; Crowl et al. 2017). The tribe Campanuleae, along with other two tribes, Cyanantheae and Wahlenbergieae, forms the Campanuloideae subfamily (or Campanulaceae sensu stricto), which features radial floral symmetry and has a center of diversity in the Holarctic region (Hong and Wang 2015). Since the description of *Campanula* L., numerous taxonomists have dedicated to proposing a “natural” infra-tribal classification based on morphological and karyological evidence, e.g., Candolle et al. (1830), Boissier (1875), Fedorov (1957), and Dambodlt (1976, 1978). While these investigations provided essential clues for understanding evolutionary relationships, convergent evolution, cryptic species, and frequent reticulate evolution events can hinder comprehensive and accurate taxonomic classification (Crowl et al. 2016, 2017). The genetic age enabled the clarification of some recalcitrant evolutionary relationships. For example, Eddie et al. (2003) estimated the phylogeny of Campanulaceae and diagnosed the polyphyly of *Campanula*, with *Edraianthus* and *Phyteuma* nested within *Campanula*, using nuclear ribosomal internal transcribed spacer (ITS) sequences (Fig. 1a). Subsequently, a series of phylogenetic studies inferred the maternally phylogenetic backbone of Campanulaceae using plastid regions (Fig. 1b-g; Mansion et al. 2012; Crowl et al. 2016; Jones et al. 2017; Yoo et al. 2018; Xu and Hong 2021); these studies confirmed the polyphyly of *Campanula*.

**Figure 1.**
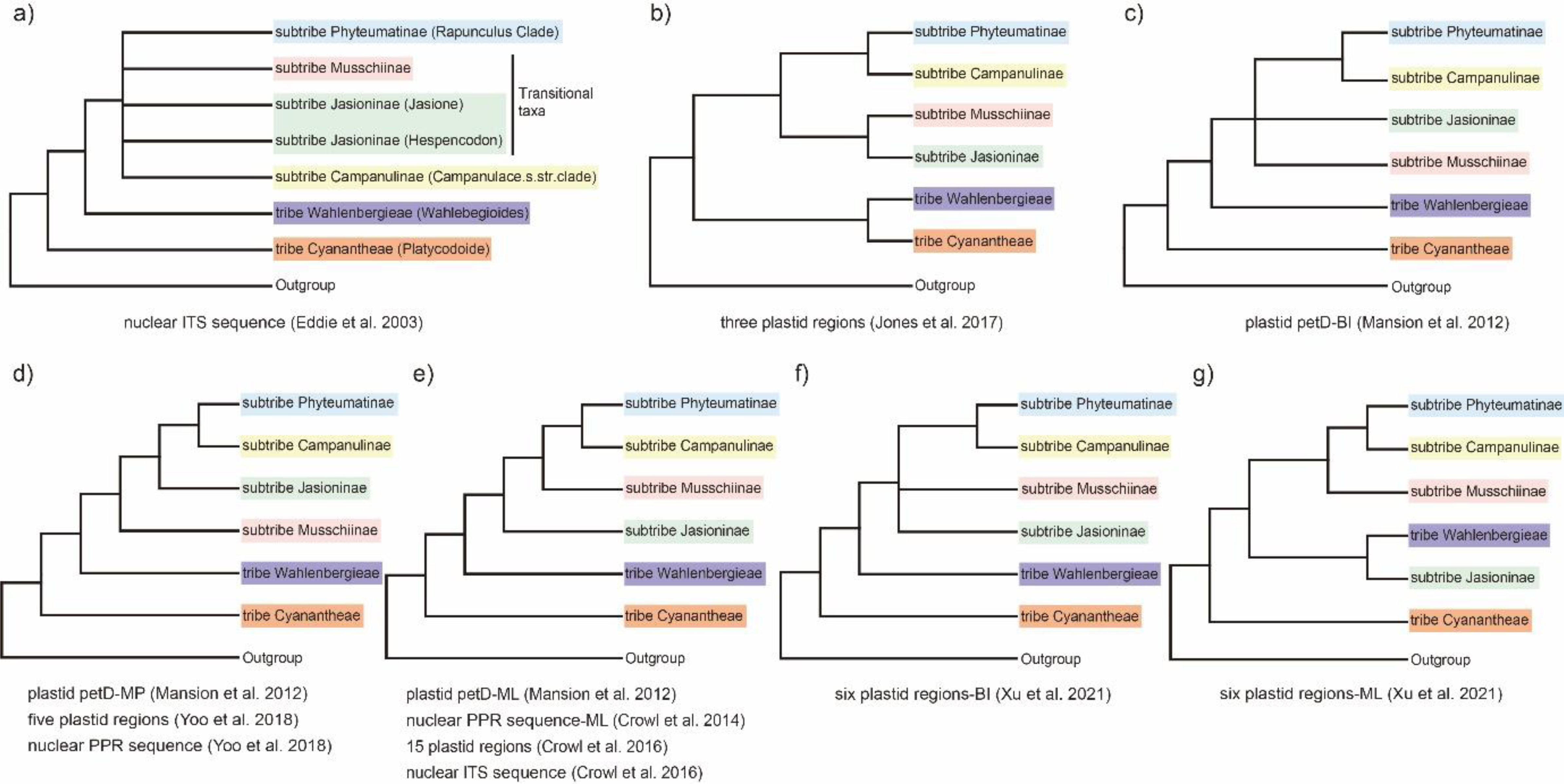
Phylogenetic hypotheses estimated in previous studies among the tribes Campanuleae, Cyanantheae, and Wahlenbergieae, particularly emphasizing the relationships among the four subtribes within the tribe Campanuleae, i.e., Campanulinae, Jasioninae, Musschiinae, and Phyteumatinae (referring to **Fig. 2a**). **a)** a combined nuclear ITS1 and ITS2 sequences (MP tree; Eddie et al. 2003). **b)** three plastid regions (*pet*D, *rpl16*, and *trn*K*/mat*K; MP, MO, and BI trees; Jones et al. 2017). **c)** plastid *pet*D sequence (BI tree; Mansion et al. 2012). **d)** plastid *pet*D sequence (MP tree; Mansion et al. 2012); five plastid regions (*atp*B, *mat*K, *pet*D, *rbc*L, and *trn*L-F) and nuclear PPR70 sequence (ML and BI trees; Yoo et al. 2018). **e)** plastid *pet*D sequence (ML tree; Mansion et al. 2012); nuclear PPR70 sequence (ML tree; Crowl et al. 2014); 15 plastid regions (ML tree; Crowl et al. 2016); nuclear ITS sequence (ML tree; Crowl et al. 2016). **f)** six plastid regions (*atp*B-*rbc*L, *mat*K, *pet*D-intron, *rbc*L, *rpl*16, and *trn*L-F; BI tree; Xu and Hong 2021). **g)** six plastid regions (ML tree; Xu and Hong 2021).

Recently, Xu and Hong (2021) generated a data matrix with extensive taxon sampling and six plastid regions, identifying four major clades (Fig. 1f, g; the Campanuleae I, II, III, and IV clades) and 24 subclades within the bellflower tribe, including 18 clades confirmed by Mansion et al. (2012) (further subdividing Cam 04 to two clades) and six separate genera: *Favratia*, *Feeria*, *Homocodon*, *Jasione*, *Peracarpa*, and *Trachelium* (Xu and Hong 2021). Because plastid markers have limited variability and maternal inheritance (Gitzendanner et al. 2018), these data painted an incomplete picture of the evolutionary relationships among the bellflower tribe. With the development of next-generation sequencing (NGS) technology and accompanying phylogenomic inference programs, we can now utilize hundreds or thousands of biparentally inherited nuclear genes, whole plastomes, and mitochondrial genes for estimating phylogenies. Newly developed approaches such as Deep Genome Skimming (DGS; Liu et al. 2021, 2022) and target enrichment sequencing (Hyb-Seq; Weitemier et al. 2014; Baker et al. 2022) are getting us closer to the goal of accurate phylogenies, which can be used to detect underlying mechanisms for gene tree and cytonuclear discordance (Guo et al. 2021). It is clear that multiple sources of genomic data can overcome some of the limitations and idiosyncracieaes involved with using single markers types.

Complementary lines of evidence have shown that reticulate evolution played a significant role in the diversification of the bellflower tribe, especially via hybridization, polyploidization, and ILS (Lammers 2007b; Crowl et al. 2017). Integrating plastomes and 130 nuclear loci, Crowl et al. (2017) uncovered cryptic tetraploid and octoploid *Campanula*, a lineage with four species from the Mediterranean, and revealed that morphological traits failed to distinguish polypoid lineages because only one parental morphology is retained. Previous cytological evidence also showed that nearly 13% of the Campanuloideae are presumed polyploid derivatives (Lammers 2007a), indicating a substantial role of polyploidization in the evolutionary history of bellflowers and their relatives.

Increasing genomic data resources in public databases, including highly reusable data types (Guo et al. 2021), such as DGS, Whole-Genome Sequencing (WGS), and transcriptomic sequencing (RNA-Seq) enable using multiple lines of genomic evidence to investigate reticulation histories. In this study, we integrate multiple genomic data sources into a phylogenomic study of Campanulaceae. In total, 659 single-copy nuclear (SCN) genes, derived from Hyb-Seq, RNA-Seq, DGS, as well as plastid protein-coding sequences (CDSs) will be used for phylogenomic analysis. Using this genomic dataset with different gene histories, we assessed cytonuclear discordance to identify potential ILS and hybridization events throughout the deep nodes along the phylogenetic backbone. Molecular dating and biogeographic analysis were used to infer when and where both recent and ancient hybridization and polyploidization occurred, promoting the diversification of Campanuleae. Explicitly, we aim to explore the utility of multi-source genomic data for 1) building a well-supported phylogenetic network backbone for a non-model plant lineage, the tribe Campanuleae, and 2) elucidating hybridization and polyploidization events deep in phylogeny based on phylogenomic and biogeographic analyses.

## Materials and Methods

### Taxon Sampling, DNA Extraction, and Sequencing

In this study, we adopted the taxonomic system as described by Lammers (2007a, 2007b), which offers a comprehensive synopsis of generic and species delimitation, and has gained extensive acceptance within the Campanulaceae community. Later, Mansion et al. (2012) subdivided Campanula sensu lato into 17 distinct clades, relying on chloroplast *petD* group II intron sequences. Our sampling was meticulously designed to encompass all 17 major clades of Campanuleae, to optimize our ability to obtain a well-supported nuclear and plastid backbone for the bellflower tribe. Specifically, our samples comprised 134 accessions, which included 110 ingroup species (116 individuals) spanning all 17 clades in Campanuleae and 18 outgroup species (Mansion et al. 2012; see Supplementary Table S1 for details).

Our approach integrated data from diverse sequencing strategies, to optimize the data available to guide phylogenomic inference, as demonstrated by Liu et al. (2021, 2022). For the phylogenomic analyses of Campanuleae, we harnessed data from DGS, WGS, Hyb-Seq, and RNA-Seq (detailed in Supplementary Table S1). For DGS, WGS, and Hyb-Seq sequencing, we extracted total genomic DNAs from silica-gel dried leaves and, where applicable, herbarium/museum specimens. The extraction was performed using the modified CTAB (mCTAB) method (Li et al. 2013) and carried out at the Institute of Botany, Chinese Academy of Science (IBCAS). For DGS and WGS, we utilized the NEB Next Ultra DNA Library Prep Kit for Illumina (NEB, USA), following the manufacturer’s guidelines. Sequencing was performed on the DNBSEQ- T7 and BGISEQ-500 Sequencing System (Novogene, Beijing), yielding paired-end reads of 2 × 150 bp for 35 accessions and 2 × 100 bp for five accessions, respectively (see Supplementary Table S1 for more information). For Hyb-Seq, library preparation was completed using the Fast Library Prep Kit (IGeneTech, Beijing). Subsequent solution-based hybridization and target enrichment were carried out with the TargetSeq One^®^ Kit at the iGeneTech facility in Zhejiang, China. We used the Illumina NovaSeq 6000 platform to generate paired-end reads (2 × 150 bp) for 82 accessions (refer to Supplementary Table S1 for details). For RNA-Seq, we extracted total genomic RNAs from silica-gel dried leaves via the mCTAB method. The libraries were prepared using the NEBNext^®^ Ultra RNA Library Prep Kit for Illumina (USA). This process resulted in paired-end reads of 2 × 150 bp, which were sequenced on the Illumina NovaSeq 6000 Sequencing System for a total of eight accessions (further details in Supplementary Table S1). All raw reads (130 accessions) sequenced for this study have been deposited in the NCBI Sequence Read Archive (SRA) under the BioProject PRJNA895940.

### Raw Reads Processing, and Nuclear SCN and Plastid Sequence Assembly

In a recent series of studies, we have developed approaches to integrate multi-source genomic data for phylogenomic analyses (Liu et al. 2021, 2022; Jin et al. 2023). Here, the processing and assembly of raw reads follow the workflow established in previous studies. After sequencing, low-quality reads and base calls were trimmed, and adaptor sequences were removed using Trimmomatic v. 0.39 (Bolger et al. 2014). The quality of the results was subsequently checked using FastQC v. 0.11.9 (Andrews 2018). Leveraging the transcriptome of *Adenophora polyantha* Nakai (SRA accession: SRX8528008), we screened putative single-copy genes with MarkerMiner v. 1.0 (Chamala et al. 2015), which yielded 659 SCN genes. The clean reads were then used to assemble the SCN genes via the HybPiper v. 2.0 pipeline (Johnson et al. 2016). Specifically, the “hybpiper assemble” command was executed to assemble contigs and extract sequences with the 659 SCN gene sequences as the references. We summarized gene recovery statistics using the “hybpiper stats” command. The gene recovery statistics were used to generate a visual representation of recovery efficiency through the “hybpiper recovery_heatmap” command, enabling us to assess the assembly quality of each gene. Finally, the “hybpiper paralog_retriever” command was run to obtain the sequences, potentially containing paralogs, for all recovered genes. This process generated an unaligned multi-FASTA file for each gene.

Given the history of gene rearrangements and diverse repeat sequences in the Campanulaceae plastome (Li et al. 2020), we focused on assembling only the protein-coding sequences (hereafter referred to as plastid CDSs) for plastid phylogenetic inference. We extracted 79 CDS sequences from three chloroplast genomes using Geneious Prime (Kearse et al. 2012); they are *Adenophora remotiflora* (GenBank accession: KP889213), *Campanula takesimana* (GenBank accession: KP006497), and *Trachelium caeruleum* (GenBank accession: EU090187). These 79 CDS sequences served as the reference for assembly. For the assembly of the plastid CDSs, we employed the HybPiper v. 2.0 pipeline (Johnson et al. 2016), mainly following the approach described in our earlier nuclear SCN genes assembly, except the final multi-FASTA file for each CDS sequence were retrieved using “hybpiper retrieve_sequence” command.

### Orthology Inference and Data Matrices Generation for Nuclear SCN Genes

Considering the prevalence of allopolyploid species documented in the Index to Plant Chromosome Numbers (IPCN), we applied several methods to identify paralogs and differentiate potential orthologs from homologs. We adopted the orthology inference methodology developed by the Ya Yang Group (Yang and Smith 2014; Morales-Briones et al. 2022). This strategy includes the Monophyletic Outgroups (MO), Rooted Ingroups (RT), and one-to-one orthologs (1to1) approaches. These methods are effective for minimizing the influence of paralogs in phylogenetic inference. The 1to1 method retains only homologs with no duplicated taxa, which avoid the introduction of potential paralogs. In the MO approach, paralogs were identified and pruned using homologs with monophyletic, non-repeating outgroups, and this process involved rerooting and pruning paralogs from root to tip. When the outgroup was absent, only those without duplicated taxa were used. Meanwhile, the RT method removed paralogs by extracting ingroup clades and cutting paralogs from root to tip.

Similarly, when the outgroup was missing, only those sequences free from duplicated taxa were considered. For both methods, we set a minimum threshold of 25 ingroup taxa. The full details of paralog assessment and orthology inference are available online (https://bitbucket.org/dfmoralesb/target_enrichment_orthology/src/master/). We generated three datasets from these analyses to use as the basis for phylogenetic inference, i.e., the 1to1, MO, and RT datasets.

### Cleaning of Nuclear and Plastid Sequences

We implemented a series of processing steps to clean low-quality sequences, a method previously employed with success in our studies, such as Liu et al. (2021, 2022) and Jin et al. (2023). To achieve refined alignments despite sequences with inconsistent quality, we utilized MAFFT v. 7.505 (Nakamura et al. 2018) to align each SCN sequence, employing the Smith-Waterman algorithm and the “--maxiterate 1000” parameter. The resulting multiple sequence alignments were trimed by trimAl v. 1.2 (Capella-Gutiérrez et al. 2009) to remove spurious sequences or poorly aligned regions. Specifically, columns with gaps in over 20% of the sequences or with a similarity score below 0.001 were removed using the parameters “-gt 0.8 -st 0.001”. Further cleaning of the sequences was performed using Spruceup (Borowiec 2019), which identified, visualized, and eliminated outlier sequences, with a window size of 50 and overlap of 25. Alignments produced before and after using Spruceup were concatenated and split, respectively, using AMAS v. 1.0 (Borowiec 2016). Recognizing that exceptionally short sequences could hinder accurate phylogenetic inference for each SCN gene, sequences shorter than 250 bp in each alignment were excluded using a Python script (exclude_short_sequences.py) from Liu et al. (2022). The cleaned sequences then served as inputs to infer gene trees through RAxML v. 8.2.12 (Stamatakis 2014), using the option “-f a” and 200 BS replicates for clade support evaluation. To ensure the accuracy of species tree inference, TreeShrink v. 1.3.9 (Mai and Mirarab 2018) was employed to identify and remove excessively long branches in each gene tree. After the above processing steps were complete, the sequences were termed ‘clean nuclear genes’, with three separate data matrices: 1to1, MO, and RT approach.

### Accurate Phylogenetic Inference with Multiple Methods

To obtain accurate phylogenies and identify the topological discordance between trees, we employed both concatenated and coalescent-based methods. The gene trees, refined by removing long branches via TreeShrink, served as the input trees to estimate the species tree using ASTRAL-III (Zhang et al. 2018), which is statistically consistency with the multi-species coalescent model. Notably, any input gene tree branches with low support (≤ 10) were collapsed using phyx (Brown et al. 2017), as collapsing gene tree nodes with BS support below a threshold value can enhance accuracy (Zhang et al. 2018). Clean nuclear genes were used for both ML and BI tree inference. The most suitable partitioning schemes and molecular evolution models were identified through PartitionFinder2 (Stamatakis 2006; Lanfear et al. 2016), with default settings. The resulting schemes and models were then used in subsequent ML tree estimates via IQ-TREE2 v. 2.2.0.3 (Minh et al. 2020) — with 1000 SH-aLRT and ultrafast bootstrap replicates — and RAxML v. 8.2.12 (Stamatakis 2014) using the GTRGAMMA model for each partition and 200 rapid bootstrap (BS) replicates for clade support. BI analysis was conducted using MrBayes 3.2.7a (Ronquist et al. 2012), running Markov Chain Monte Carlo (MCMC) analyses for 50 million generations. Stationarity was achieved when the average standard deviation of split frequencies remained under 0.01. Trees were analyzed every 1,000 generations, with the initial 25% of samples discarded as burn-in. Subsequent trees were used to generate a 50% majority-rule consensus tree.

We compiled a dataset comprising 79 plastid CDS sequences, hereafter referred to as the ‘plastid CDS dataset’, for phylogenetic analysis. While processing the plastid sequences and performing the phylogenetic inference, we largely follow the nuclear SCN methodology, excluding the steps of eliminating short sequences and long branches.

### Gene Tree and Species Tree Discordance Analyses

We used *phyparts* to calulate unique, conflicting, and concordant bipartitions within individual orthologs across the phylogeny (Smith et al. 2015). We conducted both quick concordance (-a 0) and full concordance (-a 1) analyses to address the issue arising from orthologs with missing taxa. The conflict analysis produces a pie chart for each node, segmented into five sections. These sections depict varying proportions of orthologs, such as those supporting the clade (in blue), those supporting the main alternative for that clade (in green), those supporting the remaining alternatives (in red), the uninformative (in dark grey), and the missing ones (in light grey). Additionally, we computed the ‘internode certainty all’ (ICA) scores on the input concatenated/coalescent-based tree based on the set of ortholog trees (Salichos et al. 2014). The results from *phyparts* were illustrated using phypartspiecharts_missing_uninformative.py, a Python script developed by Morales-Briones, which is available at: https://bitbucket.org/dfmoralesb/target_enrichment_orthology/src/master/phypartspiech arts_missing_uninformative.py

As a counterpart to *phyparts* in analyzing phylogenomic discordance, QS adeptly identifies discordance in large-sparse and genome-wide datasets. This method addresses challenges related to alignment sparsity and can differentiate between strong conflict and weak support (Pease et al. 2018). For each internal branch, QS produces three distinct scores: Quartet Concordance (QC), Quartet Differential (QD), and Quartet Informativeness (QI). Each approach for quantifying discord offers unique yet complementary insights. The results from QS are visually represented using the plot_QC_ggtree.R, an R package developed by Shui-Yin Liu, accessible at https://github.com/ShuiyinLIU/QS_visualization.

### Coalescence Simulation for Testing the Effect of ILS

Gene tree discordance can arise from single evolutionary events, including ILS, hybridization, and allopolyploidization, or some combination of these factors. To distinguish among these, we implemented a series of analyses. We utilized coalescence simulation to test if ILS could explain gene tree conflicts; this method has proven effective in recent research (Moralis-Briones et al. 2021; He et al. 2022; Liu et al. 2022). First, we employed the previously described method (in the phylogenetic inference section) to estimate the species tree using ASTRAL-III (Zhang et al. 2018). Using this ASTRAL ultrametric species tree, we simulated 10,000 gene trees under the multi-species coalescent (MSC) model with the “sim.coaltree.sp” function in the R package Phybase v. 1.5 (Liu and Yu 2010). We subsequently compared the distribution of tree-to-tree distances between simulated and empirical gene trees using the DendroPy v. 4.5.2 Python package (Sukumaran and Holder 2010). The result was visualized in a column chart where the extent of overlap between simulated and empirical gene tree bars represents the goodness-of-fit of the coalescent model, indicating whether ILS is a plausible explanation for gene tree discordance.

### Inference of Global Split Networks

The split network is a valuable tool for visualizing inconsistencies within a dataset. In such a network, ancestral species are not designated by specific nodes. Instead, parallel edges denote the splits that derive from the dataset, and their length indicates the importance of these splits. For the tribe Campanuleae, we constructed a split network using SplitsTree v. 4.19.0 (Huson and Bryant 2006), drawing on aligned SCN sequences from the MO dataset. This construction involved applying split decomposition to the uncorrected_P distances. Given the large divergence in this dataset, we presented the split network produced by the NeighborNet method for achieving higher resolution, and we then employed the EqualAngle network construction algorithm to infer the split network.

### Phylogenetic Network Estimation

In this study, we used network approaches investigate the early diversification of major clades within the tribe Campanuleae, encompassing three tribes and four subtribes. To reduce the computational burden of the network analysis, we selected a subset of 11 species. This dataset is specifically tailored to test the potential ancient reticulate evolution events that occurred among the three tribes and four subtribes. Given the comprehensive taxon sampling within the subtribe Phyteumatinae, we grouped all 78 samples (73 species) into six monophyletic clusters, i.e., six subclades, across 12 nuclear and four plastid trees. Additionally, we generated another dataset, comprising seven species, to assess the impacts of hybridization and allopolyploidization during the initial diversification of the subtribe Phyteumatinae.

We used Species Networks applying Quartets (SNaQ) approach, as detailed by Solís-Lemus and Ané (2016), for the Maximum pseudolikelihood estimation of species networks. This method is integrated within the PhyloNetworks package (Solís-Lemus et al. 2017) in Julia. Notably, it accounts for ILS via the coalescent model while addressing horizontal gene inheritance through reticulation nodes in the network. The methodology leverages pseudolikelihood, avoiding the intensive computation associated with full likelihood and facilitating estimations at the quartet level. This enhances computational efficiency due to its easy parallelization (Solís-Lemus and Ané 2016). For phylogenetic network analysis, we followed the comprehensive protocol provided by Solís-Lemus, available at https://github.com/crsl4/PhyloNetworks.jl/wiki. We used *h* values ranging between 0 and 6, undertaking 50 runs for the network inference.

### Allopolyploidy Analysis

Gene-tree Reconciliation Algorithm with Multi-labeled trees (MUL-trees) for Polyploid Analysis (GRAMPA) adapted an algorithm for topology-based gene-tree reconciliation to work with MUL-trees (Thomas et al. 2017), and this program, GRAMPA, can identify the parental lineages that hybridized to form auto/allopolyploids. Given the similarity between processes such as hybridization and allopolyploidization, we also tested potential auto/allopolyploidy events among the major clades in whole tree (including tribe Wahlenbergieae, tribe Cyanantheae, and four clades in tribe Campanuleae) and among the six subtribes in clade I of the tribe Campanuleae. Because GRAMPA can only infer one WGD at a time, we employed the method proposed by Morales-Briones et al. (2022) to successively explore WGD events along the backbone of a phylogeny. We applied this approach to the backbone of the tribe Campanuleae. Briefly, we specified the clades metioned above as -h1 and -h2 and perform reconciliation search using all MO ortholog trees against the MO species tree estimated with ASTRAL-III (Zhang et al. 2018). The MO ortholog trees, and species tree after removing the clade identified as polyploid in former GRAMPA analysis, were used to run the next round of analysis on the remaining clades with the same settings until no polyploid clade were detected.

### Dating Analysis and Ancestral Area Reconstruction

The earliest known macrofossils of the family Campanulaceae are seeds from *Campanula paleopyramidalis*, discovered in Miocene deposits ca. 17-16 million years ago (Mya), of Nowy Sacz in the Carpathians, Poland (Oszast and Stuchlik 1977; Łańcucka-Środoniowa 1979; Nemcok et al. 1998). These fossilized seeds have been a pivotal calibration point in several prior studies, including Cellinese et al. (2009), Mansion et al. (2012), Olesen et al. (2012), Crowl et al. (2014, 2016), and Jones et al. (2017). This species is closely related to the extant *Campanula pyramidalis*. We utilized this fossil as the MRCA of a combined clade comprising *Campanula carpatica*, *C. pulla*, *C. rainerii*, and *Favratia zoysii*. Furthermore, we incorporated two secondary calibration points to constrain the deeper nodes in Campanulaceae: the crown clade of the family Campanulaceae is dated between 72.24 to 52.66 Mya, and the crown clade of the subfamily Campanuloideae is dated between 63.52 to 44.58 Mya (Li et al. 2019).

To investigate the impact of conflicting topologies on dating analyses, we used the MCMCTree package in PAML v. 4.9j to estimate divergence times of Campanulaceae based on multiple distinct topologies (concatenated and coalescent-based trees) and datasets (plastome and nuclear). We started by determining the nucleotide substitution rate and Hessian Matrix through the MCMCTree program, with the independent rates clock model and the GTR substitution model. For each dataset, we performed two separate runs of the MCMCTree analysis, every time initiated with different seeds. The initial 1,000,000 generations of each Markov Chain Monte Carlo (MCMC) were discarded as burn-in. Subsequently, we collected samples every ten generations, amounting to a total of 500,000 samples. These samples were assessed using Tracer v. 1.7.1 (Rambaut et al. 2018) to confirm convergence and ensure that the effective sample size (ESS) of all parameters exceeded 200. Finally, we visualized the results from both runs using FigTree v. 1.4.4 and cross-validated them to ascertain convergence.

We utilized the software BioGeoBEARS v. 1.1.2 (Matzke 2018) implemented in RASP v. 4.2 to reconstruct the ancestral area of the bellflower tribe. Based on the current distribution of Campanulaceae, we classified its geographic range into six distinct regions: (A) Europe, (B) Northern Asia, (C) Africa, (D) North America, (E) Southern East Asia combined with Australasia, and (F) South West Asia. Three distinct estimated trees were used as input to test the effect of topological discordance on ancestral area reconstruction, in which a maximum of four areas were specified. Six models provided by BioGeoBEARS v.1.1.2 were utilized to estimate the biogeographic history, and the AICc values from these models were then compared to identify the optimal model for reconstructing the ancestral area of Campanuleae.

### Chromosome Number Reconstructions

We collected haploid chromosome numbers (n) from both the Chromosome Count Database (http://ccdb.tau.ac.il) and the Campanulaceae monograph by Lammers (2007a). Using the PAML ultrametric tree inferred from the nuclear concatenation-based tree as input, we employed ChromEvol v. 2.2 to infer the ancestral haploid chromosome number through a likelihood-based approach. Notably, species lacking chromosomal data are denoted with the symbol ‘X’. When a single species exhibited multiple chromosome counts, we selected the count observed most frequently for subsequent analysis. Our analysis was conducted under the parameter “_mainType = All_Models”, encompassing ten evolutionary models, and we utilized “_simulationsNum 1000000” to enhance the precision of our analysis. The best-fit model was estimated using the Akaike Information Criterion (AIC) score.

### Diversification Analyses

We used Bayesian Analysis of Macroevolutionary Mixtures (BAMM; Rabosky et al. 2014a) to estimate the diversification rate of Campanulaceae. The dated tree, constructed from the aforementioned concatenated orthologs tree, served as the input tree. We adopted a non-random incomplete taxon sampling strategy to avoid error introduced by incomplete sampling species and computed the sampling fraction for each clade within Campanulaceae. MCMC simulation with four chains was performed for 10,000,000 generations under the “species-extinction” model and sampling every 1,000 generations. The first 1,000,000 generations were discarded as burn-in, and the remaining samples were then analyzed using the R package BAMMTOOLS (Rabosky et al. 2014b) to assess whether the effective sample size (ESS) exceeded 200 and to generate plots.

## Results

### SCN and Plastid Genes Assembly and Nuclear Orthology Inference

This study integrated multiple sources of genomic data, including 82 Hyb-Seq, eight RNA-Seq, and 48 DGS data, for phylogenomic analyses. The nuclear assembly from HybPiper resulted in a variable number of SCN genes, ranging from 519 to 654 (Supplementary Fig. S1). Non-chimeric sequences recovered from HybPiper have been utilized to perform orthology inference using three different methods, and this process resulted in three distinct datasets: 1to1 containing 445 genes, MO with 645 genes, and RT with 660 genes. The final alignment of these three datasets, each with 134 taxa, contained 598,993, 864,819, and 887,333 characters, respectively. We employed HybPiper for the assembly of 79 plastome protein-coding sequences (CDSs), and the recovery efficiency was visualized as a heatmap (Supplementary Fig. S2).

### Nuclear Phylogenetic Inference and Gene Tree Discordance Analyses

Integrating concatenation and coalescent-based methods, we generated four trees for each dataset based on various phylogenetic inference methods: two trees from ML (RAxML and IQ-TREE2), one from Bayesian Inference (MrBayes), and one species tree estimated based on coalescent theory (ASTRAL-III). These 12 nuclear trees confirmed the monophyly of the subfamily Campanuloideae and the three-tribe classification within Campanuloideae: tribe Campanuleae, tribe Cyanantheae (orange), and tribe Wahlenbergieae (purple) (Supplementary Figs. S3-S14), almost all the informative gene trees were concordant with these nodes (node 1: 521 out of 522, ICA = 0.98; node 2: 510 out of 512; ICA = 0.98), and full QS support (1/-/1) (Fig. 2a; Supplementary Figs. S15-S22).

**Figure 2.**
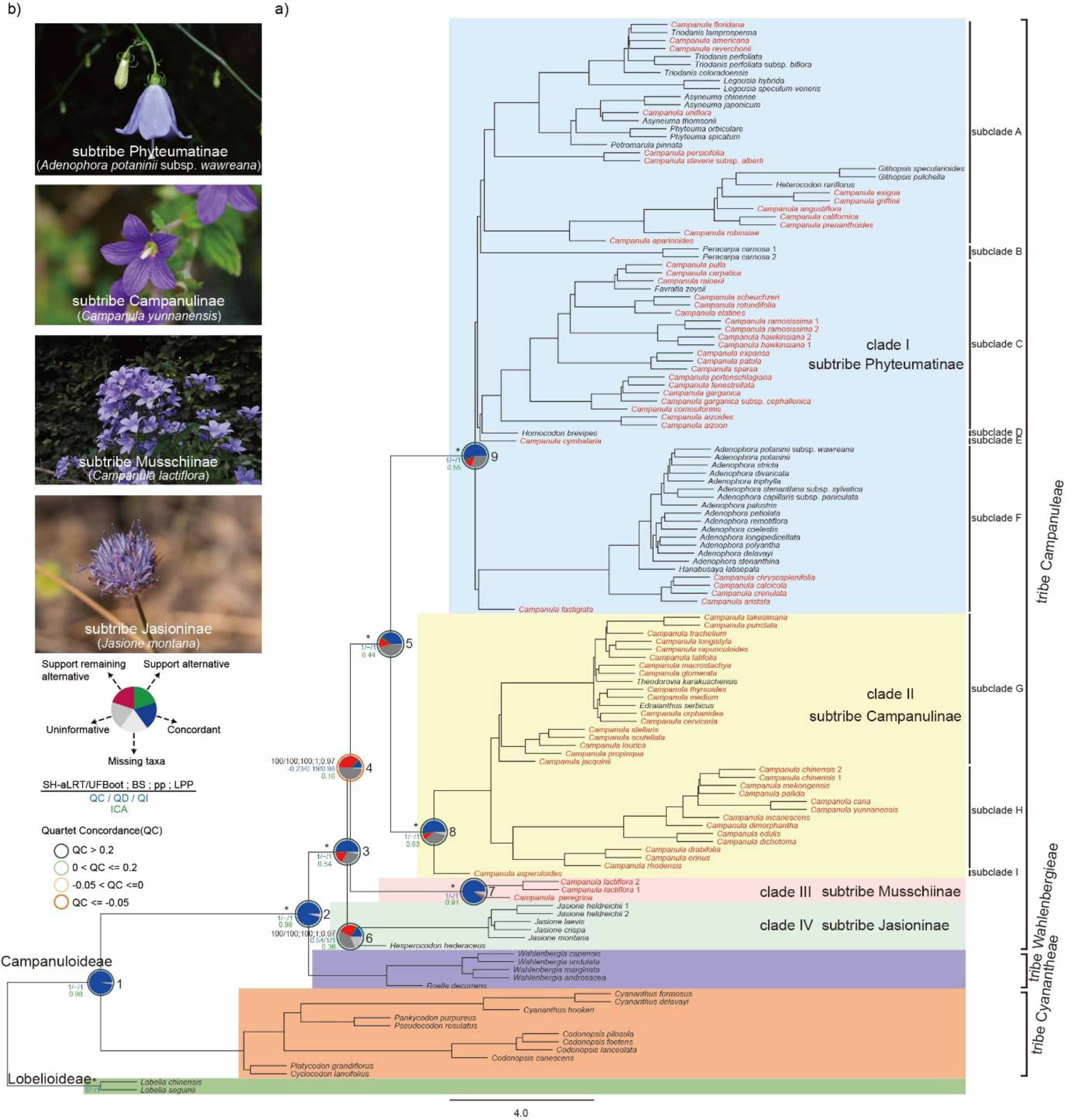
A represented tree-like phylogenetic backbone of the subfamily Campanuloideae inferred from SCN genes, emphasizing the major clades in the tribe Campanuleae. **a)** Species tree of the tribe Campanuleae in the framework of the subfamily Campanuloideae inferred from ASTRAL-III of the nuclear Monophyletic Outgroup (MO) orthologs. Summarized phylogenetic supports of the focal nine nodes from four trees based on the nuclear MO dataset were presented above the branch. From left to right (labeled in black above branch), the SH-aLRT support and Ultrafast Bootstrap (UFBoot) estimated from IQ-TREE2 (details referring to Supplementary Fig. S8); the bootstrap support (BS) values from RAxML analysis (details referring to Supplementary Fig. S7); Bayesian posterior probability values (pp) from MrBayes (details referring to Supplementary Fig. S9); the local posterior probability (LPP) from ASTRAL-III (details referring to Supplementary Fig. S10) (e.g., 100/100; 100; 1; 0.97); asterisks (*) indicated full support (100/100; 100; 1; 1). Values for Quartet Concordance/ Quartet Differential/ Quartet Informativeness estimated from Quartet Sampling analysis were provided below branches (e.g., 1/-/1, labeled in blue) (details referring to Supplementary Fig. S20). The Pie charts on these nodes illustrated the proportion of gene trees that were concordant with the corresponding clade in the species tree (blue), the proportion that supported the main alternative (green), the proportion that supported the first remaining alternative (red), the proportion considered uninformative (deep gray), and the proportion that are missing in the trees (gray); the value of partial sampling ICA were also presented below (labeled in green) (details referring to Supplementary Fig. S19). Campanuloideae was classified into three tribes, tribe Cyanatheae (orange), tribe Wahlenbergieae (purple), and tribe Campanuleae; tribe Campanuleae was further divided into four clades /subtribes, clade Ⅰ (subtribe Phyteumatinae with blue background), clade Ⅱ (subtribe Campanulinae with yellow background), clade Ⅲ (subtribe Musschiinae with pink background), and clade Ⅳ (subtribe Jasioninae with light green background); clade Ⅰ and Ⅱ were subdivided into nine subclades (A-I). All species belonging to the genus *Campanula* were highlighted in red, indicating the polyphyly of *Campanula*. **b)** Represented species of four subtribes in tribe Campanuleae, indicating the morphological diversity of flowers. From top to bottom: subtribe Phyteumatinae (*Adenophora potaninii* subsp. *wawreana*), subtribe Campanulinae (*Campanula yunnanensis*), subtribe Musschiinae (*Campanula lactiflora*), and subtribe Jasioninae (*Jasione montana*). Photos credit to You-Pai Zeng, Hai-Lei Zheng, Ke Cheng, and Jia-Nong Li (from top to bottom).

Additionally, all nuclear trees consistently supported the monophyly of four major clades within the bellflower tribe, such as clade Ⅰ (blue), clade Ⅱ (yellow), clade Ⅲ (pink), and clade Ⅳ (light green) (Fig. 2a; Supplementary Figs. S3-S14). All four clades were recovered with maximum support and full QS support (1/-/1), except clade Ⅳ which includes five individuals of *Jasione* and one species of *Hesperocodon hederaceus*—received relatively lower support in the three ASTRAL-III coalescent trees (LPP = 0.97) and strong QS support (0.54/1/1) (Fig. 2a nodes 6-9; Supplementary Figs. S6, S10, S14, and S20). Monophyly of the clade I, clade II, and clade III were supported by most of the informative trees: clade I with 296 concordant trees (out of 370 informative trees; ICA = 0.55; Fig. 2a node 9), clade II with 275 concordant trees (out of 325 informative trees; ICA = 0.63; Fig. 2a node 8), and clade III with 479 concordant trees (out of 489 informative trees; Fig. 2a node 7); but clade IV were supported by only 74 concordant trees out of 223 informative trees (ICA = 0.36; Fig. 2a node 6).

The sister group consisting of clade I and clade II was recovered with maximum support, 203 concordant trees out of 294 informative trees (ICA = 0.44), and full QS score (1/-/1) (Fig. 2a node 5). Clade III was recovered as sister of clade I + clade II with relatively high support (SH-aLRT/UFBoot = 100/100; BS = 100; pp = 1; LPP = 0.97), only 55 concordant trees (out of 208 informative trees; ICA = 0.16), and counter QS support with a strong majority of quartets supporting an alternative discordant topology (-0.23/0.19/0.98) (Fig. 2a node 4). Finally, clade IV was recovered as the sister to the rest of tribe Campanuleae with maximum support, 265 concordant trees out of 361 informative trees (ICA = 0.54), and full QS score (1/-/1) (Fig. 2a node 3).

According to the monophyletic groups recovered in the 12 nuclear trees, we subdivided clade I into six subclades (A-F) and clade II into three subclades (G-I) (Fig. 2a; Supplementary Figs. S3-S14). The phylogenetic relationships in the 12 nuclear trees were concordant for the three subclades (G-I) in clade II; however, these 12 nuclear trees revealed significant conflicting topologies in clade I.

In both the concatenation and coalescent-based tree of plastid CDSs (Fig. 3a; Supplementary Figs. S23-S28), the sister relationship between subclade F and E was recovered by 5 out of 18 informative trees (ICA = 0.07) and strong QS support with discordant skew (0.48/0.14/0.66) (Fig. 3a node 5). Subclade C was recovered as sister to a combined clade of subclade F and E with weak QS support with discordant skew (0.25/0.28/0.75), and only four concordant trees (out of 18 informative trees; ICA = 0.06) (Fig. 3a node 4). Subclade D was recovered as sister to a large clade consisting of subclade C, subclade E, and subclade F, with 22 concordant trees (out of 27 informative trees; ICA = 0.44) and full QS support (1/-/1) (Fig. 3a node 3). The sister group composed of subclade A and subclade B was recovered with full QS support (1/-/1) and 21 concordant trees (out of 28 informative trees; ICA = 0.43) (Fig. 3a node 2), and was placed as sister to the rest of clade I with full QS support (1/-/1) and 29 concordant trees (out of 36 informative trees; ICA = 0.45) (Fig. 3a node 1).

**Figure 3.**
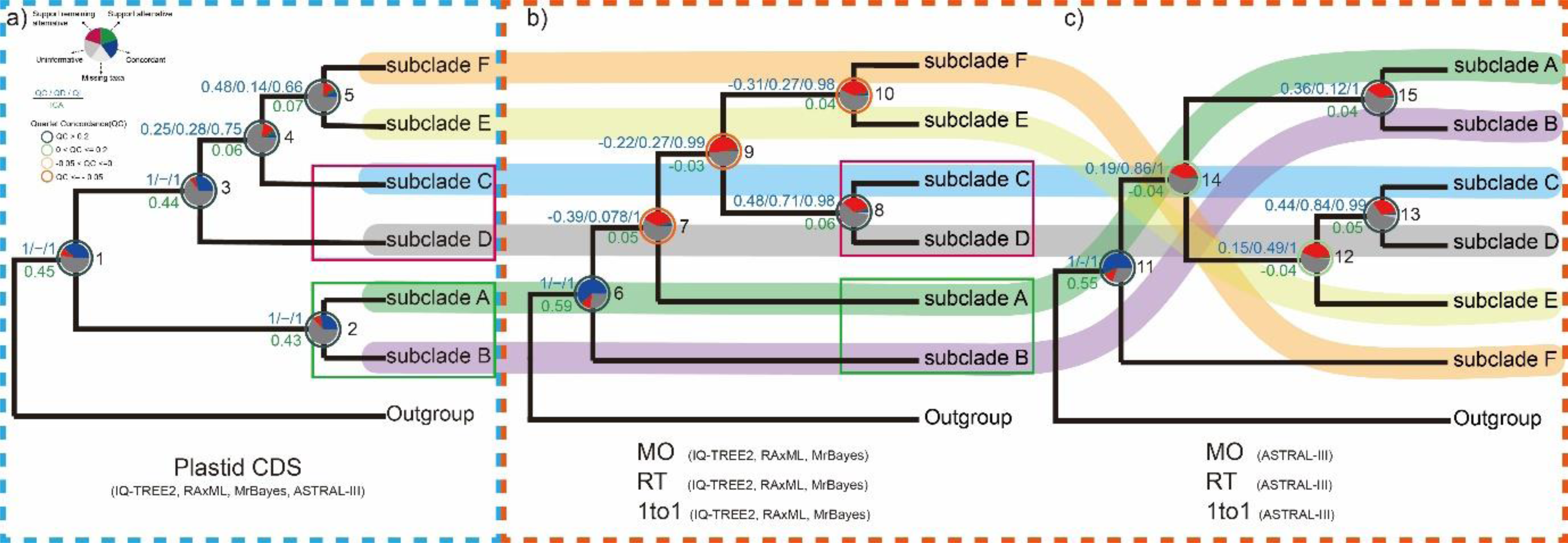
Comparative visualization of three conflicting topologies from different datasets and inference methods for clade I (equivalent to subtribe Phyteumatinae). **a)** The same topology inferred from IQ-TREE2, RAxML, MrBayes, and ASTRAL-III based on the plastid CDSs dataset. **b)** The same topology estimated by IQ-TREE2, RAxML, and MrBayes across all three nuclear datasets (MO, RT, and 1to1). **c)** The same topology estimated by ASTRAL-III across all three nuclear datasets. Quartet Sampling analysis values for QC/QD/QI are displayed above branches (e.g., 1/-/1, in blue). Pie charts on the nodes, determined by *phyparts*, illustrated the proportion of gene trees that were concordant with the corresponding clade in the species tree (blue), the proportion that supported the main alternative (green), the proportion that supported the first remaining alternative (red), the proportion considered uninformative (deep gray), and the proportion that are missing in the trees (gray). Below the branches, values of the partial sampling ICA are presented (in green). Different color bandings beneath subclades A to F visually highlight the topological discrepancies between them.

All the 12 nuclear trees recovered the monophyly of clade I with full QS support (1/-/1), and almost all the informative gene trees being concordant with these nodes (219 out of 262 for node 6, ICA = 0.59; 296 out of 370 for node 11, ICA = 0.55). However, almost all the concerned nodes within the clade I showed conflict with a little percentage of supporting trees, but QS scores of most nodes revealed supported alternative topologies (Fig. 3 nodes 6-15). The sister group consisting of subclade C and subclade D was recovered with strong QS support (i.e., a strong majority of quartets supported the focal nodes, and the low skew in discordant frequencies indicated no alternative history was favored) in both concatenation-based (node 8, 0.48/0.71/0.98) and coalescent-based trees (node 13, 0.44/0.84/0.99), moreover, a few informative trees showed concordance on this nodes (ten out of 139 for concatenation-based trees, ICA = 0.06, node 8; nine out of 187 for coalescent-based trees, ICA = 0.05, node 9). The topologies of the rest of clade I showed high level varied among concatenation and coalescent-based trees. In the concatenation-based trees, the sister group composed of subclade E and subclade F was recovered with only nine concordant trees (out of 157, ICA = 0.04, node 10), and counter QS support (- 0.31/0.27/0.98) with a skew in discordance suggesting an alternative discordant topology; furthermore, this group was placed as the sister to subclade C + subclade D, with five concordant trees (out of 185, ICA = -0.03, node 9), and counter QS support (- 0.22/0.27/0.99). Then subclade A was recovered as sister to the clade consisting of subclades C to F, with 11 concordant trees (out of 153, ICA = 0.05, node 7) and counter QS support. Finally, subclade B was the earliest-divergent lineage within clade I. In the ASTRAL trees based on nuclear, subclade E was placed as the sister to the group consisting of subclade C and subclade D, with only one concordant tree (out of 284, ICA = -0.04, node 12), and the QS score also showed weak support with discordant skew (0.15/0.49/1) indicating a possible alternative topology. The group consisting of subclade A and subclade B with was recovered with nine concordant trees (out of 218, ICA = 0.04, node 15), and weak QS support with discordant skew (0.36/0.12/1), moreover, this group was recovered as the sister of the rest subclades within clade I except subclade F, with six concordant trees (out of 230, ICA = -0.04, node 14) and weak QS support (0.19/0.86/1). Finally, subclade F was placed as the earliest-divergent lineage within clade I.

### Cytonuclear Discordance

The results from different datasets (nuclear and plastid CDSs) and various phylogenetic inference methods (concatenation and coalescent-based) consistently indicated similar topologies on the main clade nodes (Fig. 3; Supplementary Figs. S3- S14, S23-S26). However, some conflicts were observed within clade I. In the concatenation and coalescent-based trees inferred from plastid CDSs, clade I was divided into two major monophyly, subclade A and subclade B consisted the first-branding lineage, the other subclades composed another monophyly (Fig. 3a). Similar to the trees of plastid CDSs, the monophyly composed by subclades C to F was recovered in the concatenation-based trees of nuclear too, with two groups consisting respectively of subclade C + subclade D and subclade E + subclade F, but the subclade A and subclade B were placed as the first and the second branding lineages(Fig. 3b).

The topology of the ASTRAL trees was vastly different with trees of plastid CDSs, except the monophyly consisting of subclade A and subclade B, subclade C, subclade D, and subclade E consisted a group, the subclade F was placed as the earliest-divergent lineage which was sister to the other with in clade I (Fig. 3c).

### Reticulate Evolution of the Four Major Clades in the tribe Campanuleae

Reticulate evolution events were analyzed in more detail due to the strong cytonuclear discordance among several branches. We built a split network utilizing sequences from the MO dataset and designated *Roella decurrens* as the outgroup. Similar to the nuclear phylogenetic tree, the split network indicated the separation of four clades in the tribe Campanuleae; the clades were each well differentiated from the rest of the tribe, with clades III and IV being relatively close together (Fig. 4a).

**Figure 4.**
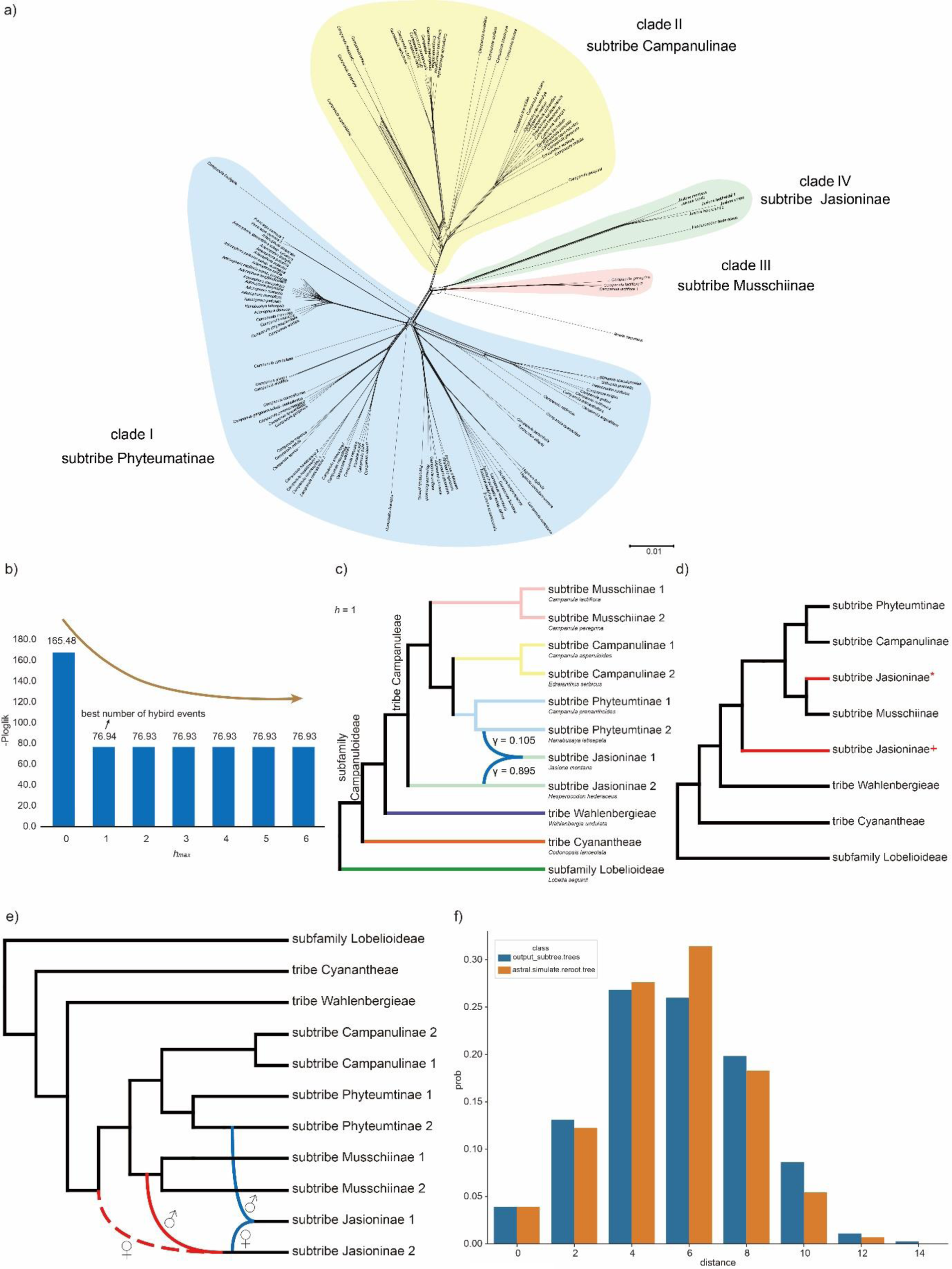
Exploring the phylogenetic network of the major clades, including three tribes in the subfamily Campanuloideae and four subtribes in the tribe Campanuleae. **a)** Split network estimated from the nuclear MO dataset, showcasing four distinct subtribes. Clade I, represented in blue, corresponds to the subtribe Phyteumatinae; Clade II, shown in yellow, aligns with the subtribe Campanulinae; Clade III, depicted in pink, associates with the subtribe Musschiinae; and Clade IV, in light green, represents the subtribe Jasioninae. **b)** Bar chart representing the pseudo-loglikelihood scores (-ploglik) across a range of maximum reticulations (from zero to six). The chart highlights the optimal number of hybrid events, with *hmax* = 1 being identified as the best number. **c)** Phylogenetic network inferred from SNaQ analysis with *hmax* = 1 as the optimal network. On the right, species names are displayed in a reduced font size beneath their corresponding tribe/subtribe names, indicating the species chosen as representatives from various subfamilies, tribes, or subtribes in the analysis. **d)** The simplified most parsimonious multi-labeled trees (MUL-trees) derived from the species tree based on the nuclear MO dataset that includes all taxa. Branches marked in red signify the allopolyploid origin of the subtribe Jasioninae, further highlighted by a red asterisk or a red plus sign from different parents. **e)** Summary of the potential phylogenetic network among the major clades within the subfamily Campanuloideae. Red curves denote the potential allopolyploidization events inferred from the GRAMPA analysis, with the dotted lines indicating extinct ancestors. The symbols ♂ and ♀ on the curving branches signify paternal and maternal parents. Blue curves highlight potential hybridization events based on the SNaQ analysis. **f)** Distribution of tree-to-tree distances between empirical gene trees and the species tree inferred from ASTRAL-III analysis, compared to those from the coalescent simulation.

Additionally, it also revealed significant levels of reticulation in the clade, implying intricate relationships among the species within tribe Campanuleae (Fig. 4a).

The coalescent simulation analysis for tribe Campanuleae suggested that ILS could not fully explain the observed conflict between gene trees and species trees (Fig. 4f). We selected specific species as representatives to investigate possible hybrid events within tribe Campanuleae. The plot of pseudo-loglikelihood scores suggested that the optimal number of hybrid events, inferred from the SNaQ network analysis, was one (Fig. 4b; Supplementary Fig. S29). This indicated the hybrid origin of subtribe Jasioninae 1, arising between subtribe Jasioninae 2 (γ = 0.895) and subtribe Phyteumatinae 2 (γ = 0.105) (Fig. 4c).

GRAMPA was employed to investigate potential allopolyploidy events associated with the origin of the subtribe Jasioninae. The inferred MUL-tree supported that the subtribe Jasioninae was of allopolyploid origin between subtribe Musschiinae and an unsampled or extinct lineage sister to tribe Campanuleae (Fig. 4d; Supplementary Fig. S30).

### Reticulate Evolution of the Six Subclades in the Subtribe Phyteumatinae

Given the intensive discordance among the topologies based on different datasets and alternative phylogenetic inference methods, several analyses were employed to test the causes. We used coalescent simulation analysis to assess the impact of ILS, and the results indicated that ILS was not the primary factor contributing to the conflict within clade I (Fig. 5b). Several species were chosen from different subclades as representatives to identify possible hybrid events within clade I. Using the SNaQ network analysis, we identified the optimal number of hybrid events (*hmax* = 1) with the best score (Fig. 5c; Supplementary Fig. S31), revealing the hybrid origin of subclade F, with parental lineages subclade E (γ = 0.728) and subclade B (γ = 0.272) (Fig. 5d).

**Figure 5.**
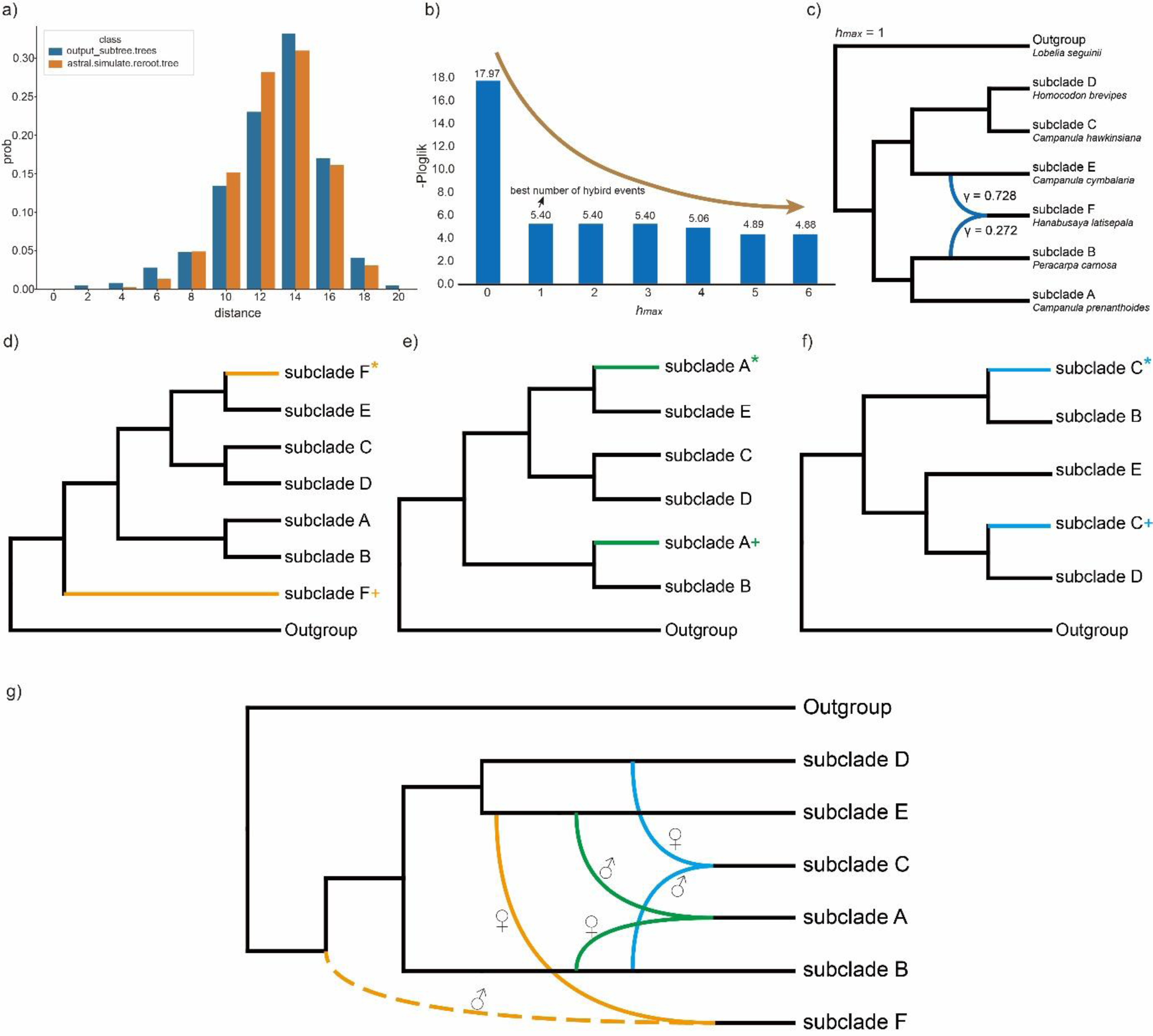
Phylogenetic network exploration of the six monophyletic groups (subclades) within the subtribe Phyteumatinae. **a)** Coalescent simulation analysis showcasing the distribution of distances between empirical gene trees and the species tree. **b)** Bar chart showcasing pseudo-loglikelihood scores (-ploglik) over a spectrum of maximum reticulations, ranging from zero to six. The chart underscores *hmax* = 1 as the optimal count for hybrid events. **c)** Phylogenetic network inferred from SNaQ analysis with *hmax* = 1 as the optimal network. To the right, representative species from various subclades are noted with their names in smaller font sizes, displayed beneath their associated subclades. **d)** The multi-labeled trees (MUL-trees) based on the species tree inferred from nuclear MO dataset, encompassing all taxa within clade I (= subtribe Phyteumatinae). Branches in orange highlight the potential allopolyploid origin of subclade F, further denoted by an orange asterisk or plus sign. **e)** The MUL-trees visualized after excluding subclade F, as depicted in d). Branches colored in green indicate the allopolyploid origin of subclade A, further marked by a green asterisk or plus sign. **f)** The MUL-trees visualized after removing subclades A and F, as referenced in e). Branches in blue highlight the allopolyploid origin of subclade C, complemented by a blue asterisk or plus sign. **g)** Summary of the potential reticulate evolutionary relationships among the six subclades within the subtribe Phyteumatinae. The orange, green, and blue curves depict the allopolyploidy events determined by the GRAMPA analysis, with the dotted line signifying an extinct ancestor. The symbols ♂ and ♀ on the curving branches signify paternal and maternal parents.

The MUL-tree inferred from GRAMPA with the lowest reconciliation score indicated the allopolyploid origin of subclade F, with the subclade E and the common ancestor of clade I as parental lineages (Fig. 5e; Supplementary Fig. S32). After removing subclade F (Fig. 5e orange), the analysis revealed subclade A as an allopolyploid clade, with parental lineages including subclades E and B (Fig. 5f; Supplementary Fig. S33). Finally, upon removing subclade A (Fig. 5f green), the analysis found subclade C to have an allopolyploid origin clade between subclade B and subclade D (Fig. 5g; Supplementary Fig. S34).

### Dating Analyses, Ancestral Area, and Chromosome Number Reconstructions

We constrained nodes with a fossil calibration point to estimate divergence times and diversification dynamics based on three different phylogenies (ML tree and ASTRAL tree inferred from the MO dataset; ML tree inferred from the plastid CDSs) within the subfamily Campanuloideae. The result based on all three phylogenetics indicated that the tribe Campanuleae originated in Europe-South West Asia during the early Eocene, diverging at *c.* 52.32 million years ago (Mya) (95% highest posterior density (HPD): 56.19-48.5 Mya) (Fig. 6a; Supplementary Figs. S35-S37).

**Figure 6.**
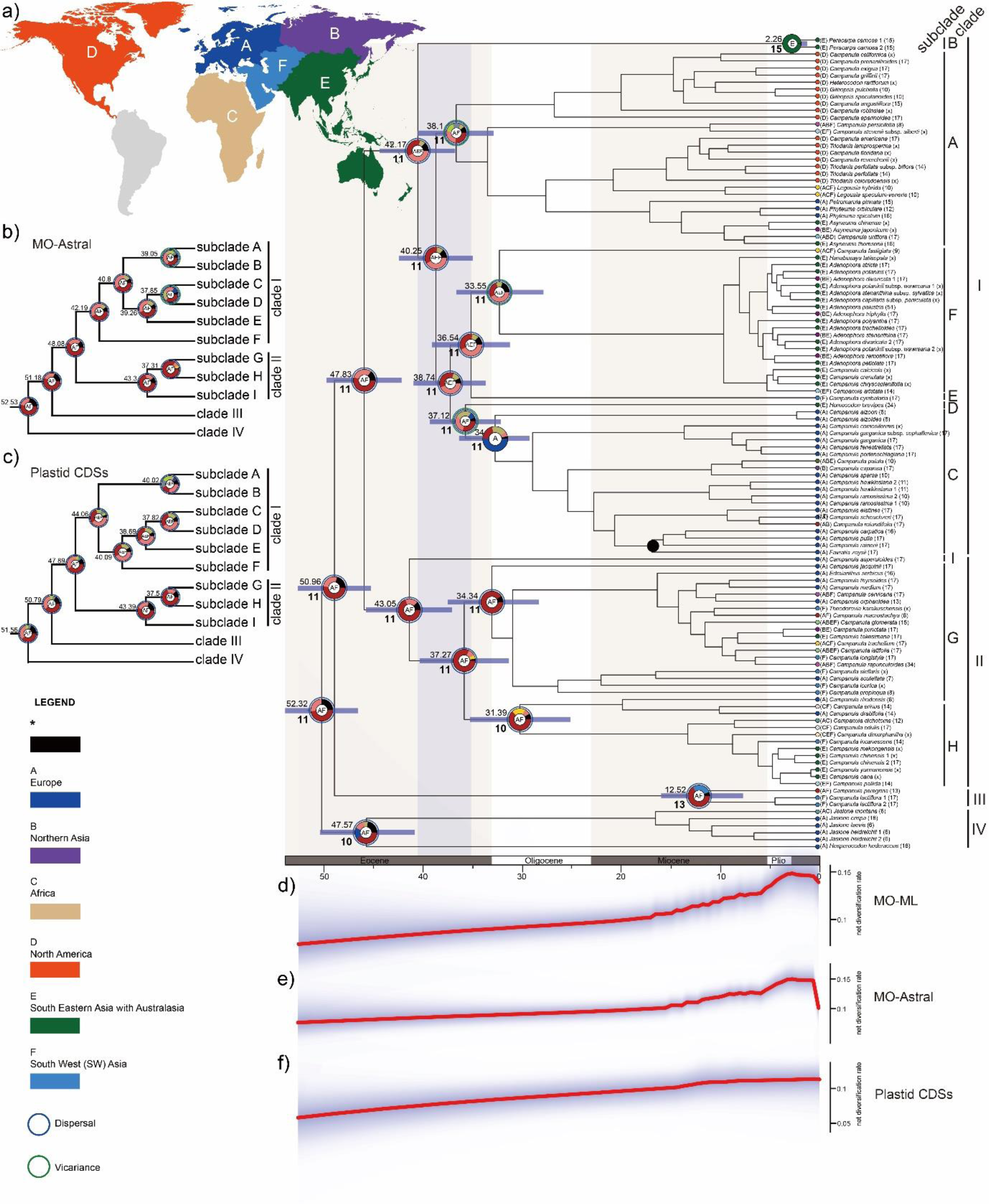
Divergence time estimation and geographical range evolution of the bellflower tribe Campanuleae. **a)** Dated chronogram of the tribe Campanuleae inferred from PAML based on the nuclear MO dataset. Focal nodes feature estimated divergence times and ancestral geographical ranges, with the purple shading highlighting the divergence time range for the six subclades within clade I (equivalent to subtribe Phyteumatinae). The inset map in the upper left outlines the six distribution areas used for geographical analysis: (A) Europe, (B) Northern Asia, (C) Africa, (D) North America, (E) Southern East Asia with Australasia, and (F) South West (SW) Asia. A black circle denotes the fossilized seed constraint. The gametophytic chromosome count was displayed in brackets next to the relevant species name, and the boldface numbers represent the reconstructed ancestral chromosome counts. **b)** The ASTRAL-estimated species tree of the nuclear MO dataset with estimated divergence time and geographical range marked on focal nodes. **c)** The plastid CDSs-inferred topology with estimated divergence time and geographical range marked on focal nodes.**d)** Net diversification rate (species/my) of Campanuleae estimated from the Maximum Likelihood (ML) phylogeny using the nuclear MO dataset. The red line represents the median, while the shaded blue area demarcates the 95% credible intervals for the rate. **e)** The net diversification rate estimated from the species tree of ASTRAL-III based on the nuclear MO dataset. **f)** The net diversification rate from the plastid phylogeny.

Furthermore, the majority of clades underwent divergence during the Eocene to the Oligocene period. Clade I had its origins in Europe-South Eastern Asia with Australasia-South West Asia by the middle Eocene, and diverged at *c.* 42.17 Mya (95% HPD: 46.22-38.22 Mya) based on the ML-trees inferred from the MO dataset and plastid CDSs (Fig. 6a-c; Supplementary Figs. S35-S43). In contrast, the result based on the ASTRAL-tree inferred from the MO dataset indicated Europe-South West Asia as the origin of clade I (Fig. 6b). Within clade I, six subclades underwent divergence during the Eocene by around *c*. 42.17-36.54 Mya based on the MO dataset (Fig. 6a light purple shadow, 6b), but a little earlier based on the plastid CDSs which indicated the divergence by *c*. 44.06-37.82 Mya (Fig. 6c). In addition, clade II diverged around *c*. 43.05 Mya (95% HPD: 47.62-38.55 Mya), and clade IV followed suit at *c*. 47.57 Mya (95% HPD: 52.5-42.48 Mya). In contrast, clade III experienced diversification in the Miocene, occurring approximately *c*. 12.52 Mya (95% HPD: 16.46-7.9 Mya) (Fig. 6a). The diversification rates, as inferred from both the MO dataset and the plastid CDSs, exhibited an increasing trend from the Eocene to the present. Moreover, the rates inferred from the MO dataset showed sudden increases in the late Miocene, which indicated rapid speciation (Fig. 6d-f).

The model CONST_RATE_DEMI_EST, as determined by ChromEvol, was identified as the best fit for chromosome evolution (AIC = 503.2). Using this model, we deduced that the most probable ancestral haploid chromosome number (n = 11) was shared by the subfamily Campanuloideae and the tribe Campanuleae. This count was consistently observed in the MRCA of the tribes Cyanantheae, Wahlenbergieae, and Campanuleae. Within the tribe Campanuleae, the MRCA of both clades I and II maintained the ancestral haploid chromosome number 11. In contrast, the MRCA of clade III presented an ancestral chromosome number of 13, and clade IV had a count of 10. For most subclades within clades I and II, the ancestral number remained 11, with the exception of subclade B, which exhibited a chromosome number of 15 (Fig. 6a; Supplementary Fig. S44).

## Discussion

By integrating multiple genomic data sources and various phylogenetic inference methods, we produced a robust phylogenetic backbone for the bellflower tribe Campanuleae within the subfamily Campanuloideae. Our analyses of deep reticulate evolution suggest that hybridization and allopolyploidization have played a pivotal role in the diversification of the Campanuleae tribe, especially in the early diversification of the subtribe Phytemtinae. We demonstrated that fully untangling deep reticulation is optimally accomplished by leveraging data from multiple genomic sources, especially when inferring the phylogenetic backbone of non-model plant lineages. Additionally, thorough taxon sampling is also critically important to fully dissect deep reticulation. Moreover, it is essential to note that conflicting topologies derived from separate genomic datasets and inference methods can significantly influence downstream inferences, such as dating analyses, ancestral area determination, and diversification rate estimates.

### Exploring the Reticulate Phylogenetic Backbone of the Bellflower Tribe Campanuleae

In traditional phylogenetic models, evolutionary relationships are portrayed as bifurcating branches, suggesting that species or lineages diverge and proceed to evolve independently. However, this strictly branching framework often fails to capture the complexity of evolutionary dynamics as they occur in nature (Jin et al. 2023). In many instances, evolutionary processes are better represented as being network-like, where species or lineages may engage in various forms of hybridization or experience intricate, web-like interactions (Stull et al. 2023). Such network-like patterns can arise from various complex processes involving multiple and ongoing events, such as ILS, introgression, hybridization, and/or allopolyploidization.

In this study, we undertook a comprehensive approach integrating extensive taxon sampling with diverse genomic data types, including DGS, Hyb-Seq, RNA-Seq, and WGS. Utilizing state-of-the-art automated tree-based orthology inference methods (Yang and Smith 2014; Morales-Briones et al. 2022), we carefully estimated paralogs and generated three alternative nuclear data matrices (1to1, MO, and RT), along with a complete dataset of 79 plastid CDSs for phylogenomic analysis. Employing various phylogenomic inference methods, such as concatenated and coalescent-based methods, we produced 16 phylogenetic trees—12 nuclear and four plastid. These trees confirmed the classification of three tribes as previously outlined by Hong and Wang (2015) and recovered four strongly supported major clades—namely Clade I, II, III, and IV—in the tribe Campanuleae (Fig. 2; Supplementary Figs. S3-S14, S23-S26). Nevertheless, our findings also highlighted the ambiguous phylogenetic position of Clade IV (or the subtribe Jasioninae), which appears to be either a sister group to the other three clades in Campanuleae (Fig. 2; Supplementary Figs. S3-S14, S23, S25 and S26) or related to a combined clade (Clade I + II, as shown in Supplementary Fig. S24). We hypothesized that the evolutionary relationships among these four major clades in the Campanuleae tribe may not be strictly bifurcating but could exhibit a more network-like structure. This hypothesis was corroborated by our SplitsTree network analysis (Fig. 4a).

Tree incongruence has been a prevalent and challenging issue in molecular phylogenetic studies (Wendel and Doyle 1998; Som 2015; Kapli et al. 2020; Steenwyk et al. 2023). Such incongruence can arise from both biological and non-biological factors. Non-biological factors primarily encompass limited taxon and/or gene sampling, the use of paralogous genes, and errors in sequence alignment (Steenwyk et al. 2023). In our study, these factors seem insufficient to explain the observed incongruence. Our extensive taxon sampling and use of genomic-level data largely rule out stochastic uncertainties from insufficient taxon and/or gene sampling. Furthermore, we used three strategies of orthology inference to reduce the effect of paralogs, and it is unlikely the observed topological inconsistencies among individual gene trees can be attributed solely to paralogy. In addition, sequence alignments have undergone meticulous visual examination and are mostly well-aligned. Tools like trimAl help trim low-quality alignments, and TreeShrink filters out samples with aberrantly long branches. This suggests that our sequence alignment is unlikely a major factor in tree incongruence. Biologically, tree incongruences can result from ILS, hybridization/introgression, or horizontal gene transfer (HGT; Kapli et al. 2020). While HGT is more common in bacteria and archaea, it is less so in eukaryotes (Kurland et al. 2003; Keeling and Palmer 2008; Boto 2010), making it an unlikely cause of the tree incongruence in our case. On the other hand, both ILS and hybridization/introgression are established contributors to tree incongruence across the tree of life (Degnan and Rosenberg 2009; Mallet et al. 2016). Thus, it is plausible that these factors, individually or in combination, are behind the observed tree incongruence in Campanuleae.

Distinguishing between ILS and hybridization/introgression based solely on patterns of tree incongruence is challenging. Both processes can produce similar incongruence patterns, complicating the identification of the specific cause of incongruence in phylogenetic studies (Degnan and Rosenberg 2009; Degnan 2018). To further dissect the potential events contributing to this network-like evolution, we performed step-by-step analyses to distinguish the possible events in this net-like evolution. Coalescent simulation analyses indicated that ILS alone could not account for the observed reticulate patterns (Fig. 4f). Subsequent PhyloNetwork analyses using the SNaQ algorithm and polyploidy assessments with the GRAMPA reconciliation algorithm were carried out to explore the potential roles of hybridization and polyploidy.

Given the strongly supported gene tree and species tree discordance for the phylogenetic position of clade III and IV in different datasets and inference methods (Fig. 1), the phylogenetic position of subtribe Jasioninae remains a subject of debate. As we delved into the diversification of the three tribes and four subtribes, we pinpointed an allopolyploidy event (Fig. 4b,c; Supplementary Fig. S31) and a distinct hybridization event (Fig. 4d; Supplementary Fig. S30). The GRAMPA analysis unveiled the allopolyploid origin of subtribe Jasioninae, resulting from the MRCA of the tribe Campanuleae (acting as the maternal lineage) and the MRCA of the subtribe Musschiinae (serving as the paternal lineage, Fig. 4d,e). Our analyses of dating and ancestral area reconstruction consistently identified the likely occurrence of this allopolyploidization event in Europe and SW Asia (around the Tethys Sea) during the early Eocene, as supported by both nuclear (52.32 Mya from the ASTRAL species tree and 52.53 Mya from the ML tree) and plastid (51.56 Mya) topologies (Supplementary Figs. S35-S43). Further, our PhyloNetwork results confirmed the hybrid origin of *Jasione montana* between the MRCA of *Hanabusaya latisepala* (paternal lineage) and the MRCA of *Hesperocodon hederaceus* (maternal lineage). Nevertheless, the dating analyses of the stem clade of *Jasione montana* showed disparities between nuclear (17.08 Mya estimated from the ML tree and 16.53 Mya from the ASTRAL species tree) and plastid data (2.08 Mya). Considering these strongly supported but discordant time estimates, we hypothesize that the MRCA of *Jasione montana* might have captured the chloroplast genome from a particular *Jasione* species, an event possibly occurred in Europe during the early Quaternary.

Within the subtribe Phyteumatinae, we conducted thorough taxon sampling encompassing all 15 clades, as confirmed by Xu and Hong (2021). Significant nuclear/plastid gene trees and cytonuclear discordance were detected among the 16 generated trees (Fig. 2; Supplementary Figs. S3-S14, S23-S26). We classified the 15 clades of subtribe Phyteumatinae into six monophyletic groups, labeled as subclades A-F (Figs. 3, 5). We utilized an iterative strategy to estimate the potential polyploidy origin for each subclade. The first GRAMPA reconciliation identified an allopolyploid origin of subclade F, arising between the MRCA of subclade E and the MRCA of subtribe Phyteumatinae (Fig. 5d). The close relationship of subclade F with subclade E in the plastid topologies (Fig. 3a) suggests that the MRCA of subclade E likely served as the maternal parent, while the MRCA of subtribe Phyteumatinae may have acted as the paternal parent. This allopolyploidization is partially consistent with the hybrid origin proposed by the SNaQ analysis, designating subclade E (γ = 0.728) as the maternal parent and subclade B (γ = 0.272) as the paternal parent (Fig. 5b,c). Interestingly, these two methodologies inferred different paternal origins. This discrepancy, marked by an uneven inheritance probability (γ = 0.728 vs. 0.272), could clarify the conflicting topologies of subclade F in both the nuclear concatenated-based analyses and species tree (Fig. 3b,c). As a result, we concluded that the origin of the subclade F could be traced to multiple, undistinguishable hybrid swarms involving the MRCA of subclade E and the MRCA of subtribe Phyteumatinae, all of which were accompanied by a whole genome duplication (WGD) event. After excluding subclades F and A, the origins of subclades A and C were identified as allopolyploid, arising from the combination of subclades E and B, and B and D, respectively (Fig. 5e,f). Our dating analysis, based on three distinct topologies depicted in Figure 6abc, supports the rapid radiation hypothesis for the six subclades (A to F) highlighted in the purple-shaded area of Figure 6a. This radiation possibly took place during the Middle to Late Eocene, with date ranges of 42.17-36.54 Mya from the nuclear ML topology, 42.19-37.85 Mya from the nuclear species tree, and 44.06-37.82 Mya from the plastid ML topology. Notably, discrepancies were observed between our nuclear and plastid topologies regarding the ancestral areas of the subtribe Phyteumatinae. The nuclear topologies suggest Europe & SW Asia (the Ancient Mediterranean region) as the ancestral areas, while the plastid topology points to a broader region encompassing Europe, SW Asia, and Southern East Asia and Australasia—essentially spanning much of Southern Eurasia. Given that plastid data may have a narrower genetic diversity (due to uniparental inheritance), they might not capture the full range of ancestral distributions compared to nuclear data. We concluded that the possible ancestral areas of the subtribe Phyteumatinae included the Ancient Mediterranean region. The European and Southwest Asian regions experienced pivotal transitions in both climate and tectonics. This era marked a departure from the extremely warm in the early Eocene to a slightly cooler, yet still warm global environment. It was also a time of significant tectonic events; the collision between the African, Eurasian, and Anatolian tectonic plates continued during this period, resulting in the formation of the Alpine mountain chain and uplift of mountain ranges like the Taurus and the Zagros. Additionally, the closure of the Tethys Sea also had significant impacts on ocean circulation patterns and climate. The extreme climate and tectonic changes may have promoted the rapid diversification of the six major clades in subtribe Phyteumatinae.

The notable cytonuclear inconsistency in dating analyses, ancestral area reconstruction, net diversification rates analyses (Fig. 6) underscores a critical lesson: relying solely on a singular genomic data source might not furnish a comprehensive or precise depiction of biogeographic evolutions for potential reticulate lineages (Dong et al. 2022; Liu et al. 2022). This disparity underscores the need for integrative and multi-faceted approaches in phylogenomic research, ensuring a more holistic understanding of evolutionary histories and events. Utilizing multiple data sources can bridge knowledge gaps, validate findings, and offer richer insights into the complexities of lineage diversification and adaptation.

### Multi-source Genomic Data for Phylogenomic Analyses of Non-model Plants

Phylogenomics has revolutionized our understanding of the evolutionary relationships among organisms by incorporating hundreds to thousands of nuclear genes in advanced phylogenetic inference methods. However, applying phylogenomics to non-model plants presents several challenges. Non-model plants display vast diversity, making it difficult to obtain representative samples that encompass the entire phylogenetic and geographical diversity of a particular group. Researchers from specific regions may have an advantage in collecting taxon samples from their areas, but conducting comprehensive species sampling, especially for cosmopolitan lineages, becomes a challenge for a single laboratory. As a result, previous phylogenomic studies have mainly focused on resolving the phylogenetic backbone of major lineages at the family or even higher level, such as Xiang et al. (2017), Zhao et al. (2021), Huang et al. (2022), Hu et al. (2023), and Zhang et al. (2023), due to the ease of conducting comprehensive taxon sampling covering most major clades. However, incomplete taxon sampling has substantially hindered shallow-level phylogenomic studies of non-model plants, except for some ecologically and economically important lineages, such as apples (Liu et al. 2022) and *Rhododendron* (Xia et al. 2022). Moreover, limited sampling can lead to biased inferences and inaccurate estimation of evolutionary relationships (Heath et al. 2008; Nabhan and Sarkar 2011; Young et al. 2020). Limited or biased sampling can be particularly problematic when reticulation is prevalent because recent hybridization can obscure accurate inference of more ancient reticulation events.

To address these challenges, Guo et al. (2021) proposed several criteria for an optimal sequencing strategy for non-model plants, which recommended generation of hundreds to thousands of SCNs from a large number of samples. As Next-Generation Sequencing (NGS) technologies have developed, researchers in plant taxonomy, evolutionary biology, and horticulture have sequenced various types of genomic-level data from different species within their respective countries. Most of these raw data have been deposited in public data repositories, such as Sequence Read Archive (SRA) in the National Center for Biotechnology Information (NCBI), National Genomics Data Center (NGDC), and European Nucleotide Archive (ENA). Since a single sequencing technology does not entirely fulfill the requirements for non-model plant phylogenomics, our objective is to propose an alternative method by integrating all available genomic data and employing a feasible bioinformatic pipeline. Liu et al. (2021, 2022) introduced a practical and innovative approach for assembling SCN genes and plastomes, as well as conducting phylogenomic discordance analyses in non-model plants. The researchers identified three primary data types that proved to be particularly suitable for their methodology: DGS, WGS, and RNA-Seq data. Each data type offered unique advantages in contributing to a comprehensive understanding of the genomic landscape and evolutionary history of non-model plants. DGS data, which were generated with approximately 10× coverage, and WGS data, with higher coverage at around 20×, were both derived from whole genome DNA libraries. These two approaches complemented each other, with DGS providing a cost-effective means of obtaining broader genomic coverage and WGS offering higher sequencing depth for more accurate variant calling and detection of rare genetic variations. The combination of both DGS and WGS data allowed for a robust analysis of genomic diversity and evolutionary patterns within the studied plant lineages. In addition to DGS and WGS, the researchers utilized RNA-Seq data, which represented the complete set of RNA transcripts of expressing coding regions. Moreover, the focus on nuclear protein-coding genes, particularly nuclear SCN genes, in current phylogenomic studies offered an effective strategy to resolve the phylogenetic relationships of non-model plants. The conserved nature of nuclear SCN gene ensures their suitability for inferring evolutionary relationships across a wide range of plant species. Leveraging these genes provided a stable and well-supported phylogenetic backbone for the studied lineages, allowing researchers to confidently infer the evolutionary history of non-model plants.

Collectively, the integration of DGS, WGS, and RNA-Seq data, along with the emphasis on nuclear SCN genes, has opened up new avenues of research in the phylogenomics of non-model plants. This integration of multi-source genomic data approach has empowered researchers to explore the rich genomic diversity and evolutionary patterns of diverse plant lineages more comprehensively and accurately. By leveraging the various types of available data and employing innovative bioinformatic pipelines, researchers can now gain deeper insights into the evolutionary relationships, historical biogeography, and functional implications of genetic variations in non-model plants, ultimately advancing our understanding of plant evolution and diversification. Continued advancements in sequencing technologies and analytical methods are expected to further enhance the potential of phylogenomics, opening up exciting possibilities for unraveling the mysteries of plant biodiversity and adaptation.

### An updated infra-tribal classification of the tribe Campanuleae

**Campanulaceae** Juss., Gen. Pl. [Jussieu] 163 (1789), nom. cons. Type: *Campanula* L. Subfamily **Campanuloideae** Burnett, Outlines Bot. 942, 1094, 1110 (1835). Type: *Campanula* L.

Tribe **Campanuleae** Dumort., Fl. Belg. 58. 1827. Type: *Campanula* L.

= tribe Jasioneae Dumort., Fl. Belg. 59. 1827. Type: *Jasione* L.

= tribe Phyteumateae Dumort., Fl. Belg. 59. 1827, ‘Phyteumeae’. Type: *Phyteuma* L.

= tribe Peracarpeae Fed., Fl. URSS 24: 471. 1957. Type: *Peracarpa* Hook.f. & Thomson.

= tribe Michauxieae Fed., Fl. URSS 24: 472. 1957. Type: *Michauxia* L’Hér.

= tribe Edraiantheae Fed., Fl. URSS 24: 475. 1957. Type: *Edraianthus* A.DC.

= tribe Annaeeae Kolak., Bot. Zhurn. (Moscow & Leningrad) 72(12): 1575. 1987, ‘Annaea’. Type: *Annaea* Feer.

= tribe Azorineae Kolak., Bot. Zhurn. (Moscow & Leningrad) 72(12): 1575. 1987. Type: *Azorina* Feer.

= tribe Echinocodoneae Kolak., Bot. Zhurn. (Moscow & Leningrad) 72(12): 1575.

1987, nom. illeg. Type: *Echinocodon* Kolak., Soobshch. Akad. Nauk Gruz. SSR, 121(2): 387. 1986, hom. illeg. non D.Y. Hong, Acta Phytotax. Sin., 22(3): 183. 1984.

= tribe Gadellieae Kolak., Bot. Zhurn. (Moscow & Leningrad) 72(12): 1576. 1987. Type: *Gadellia* Schulk.

= tribe Muehlbergelleae Kolak., Bot. Zhurn. (Moscow & Leningrad) 72(12): 1575. 1987. Type: *Muehlbergella* Feer.

= tribe Musschieae Kolak., Bot. Zhurn. (Moscow & Leningrad) 72(12): 1575. 1987. Type: *Musschia* Dumort.

= tribe Mzymteleae Kolak., Bot. Zhurn. (Moscow & Leningrad) 72(12): 1578. 1987. Type: *Mzymtella* Kolak.

= tribe Neocodoneae Kolak., Bot. Zhurn. (Moscow & Leningrad) 72(12): 1577. 1987. Type: *Neocodon* Kolak. & Serdyuk.

= tribe Sachokieleae Kolak., Bot. Zhurn. (Moscow & Leningrad) 72(12): 1578. 1987. Type: *Sachokiella* Kolak.

= tribe Sergieae Kolak., Bot. Zhurn. (Moscow & Leningrad) 72(12): 1577. 1987. Type: *Sergia* Fed.

= tribe Theodorovieae Kolak., Bot. Zhurn. (Moscow & Leningrad) 72(12): 1575. 1987. Type: *Theodorovia* Kolak.

= tribe Echinococonieae Kolak., Bot. Zhurn. (Moscow & Leningrad) 79(1): 114. 1994. Type: *Echinocodonia* Kolak.

= tribe Pseudocampanuleae Kolak., Bot. Zhurn. (Moscow & Leningrad) 79(1): 115. 1994. Type: *Pseudocampanula* Kolak.

1. subtribe **Campanulinae** R.Schönland (clade II) in H.G.A. Engler & K.A.E. Prantl, Nat. Pflanzenfam. IV, 5(48). 1889. Type: *Campanula* L.

Included genera: *Azorina*, *Campanula*, *Edraianthus*, *Michauxia*, *Muehlbergella*, *Sachokiella*, *Theodorovia*, *Trachelium*, and *Zeugandra*.

II. subtribe **Phyteumatinae** Caruel (clade I), Epit. Fl. Europ. 2: 248. 1894, ‘Phyteumeae’. Type: *Phyteuma* L.

Included genera: *Adenophora*, *Astrocodon*, *Asyneuma*, *Brachycodonia, Cryptocodon*, *Cylindrocarpa*, *Decaprisma*, *Eastwoodiella*, *Favratia*, *Githopsis*, *Hanabusaya*, *Hayekia*, *Heterocodon*, *Homocodon*, *Legousia*, *Loreia*, *Melanocalyx*, *Palustricodon*, *Peracarpa*, *Petromarula*, *Physoplexis*, *Phyteuma*, *Poolea*, *Protocodon*, *Ravenella*, *Rotanthella*, *Sergia*, *Smithiastrum*, and *Triodanis*.

III. subtribe **Musschiinae** B.B.Liu (clade III), **stat. nov.** basionym: tribe Musschieae Kolak. Bot. Zhurn. (Moscow & Leningrad) 72: 1575. 1987. Type: *Musschia* L. Included genera: *Echinocodonia*, *Gadellia*, and *Musschia*.
IV. subtribe **Jasioninae** Endl. (clade IV), Gen. Pl. [Endlicher]. 514. 1838, ‘Jasioneae’. Type: *Jasione* L.

Included genera: *Feeria*, *Hesperocodon*, and *Jasione*.

## Conclusions

We focused on the bellflower tribe Campanuleae, a non-model plant lineage known for its extensive history of hybridization and introgression. We presented a comprehensive and versatile framework for deciphering the network-like phylogenetic relationships within such lineages, relying on various genomic data sources and analytical methods. Our results unveiled compelling evidence supporting the role of allopolyploidization and hybridization in promoting the early diversification of the Campanuleae tribe. Notably, the rapid radiation of six major subclades in the subtribe Phyteumatinae may have been driven by multiple and continuous allopolyploidization events, taking place in the ancient Mediterranean region during the Middle to Late Eocene epochs. Furthermore, our study emphasizes a significant challenge in evolutionary biology research: conflicting tree topologies derived from different genomic datasets and phylogenetic inference methods can lead to substantial variation in our downstream estimates of the timing of evolutionary events, ancestral geographic origins, and diversification rates.

## Supplementary Material

Data available from the Dryad Digital Repository: https://doi.org/10.5061/dryad.hqbzkh1nz.

## Acknoledgements

All the phylogenomic analyses have been run on the PhyloAI supercomputer, owned by B.B.L., Institute of Botany, Chinese Academy of Sciences.

## Author Contributions

B.B.L. designed the project and supervised the study. B.B.L., Z.T.J., S.Y.X., and C.X. wrote the draft manuscript. Z.T.J., S.Y.X., and Y.Z. carried out the phylogenomic analyses. C.X. performed the deep genome skimming sequencing. B.L. provided suggestions for taxonomic classification. G.N.L., H.W., X.H.L., R.G.J.H., D.K.M., S.H.J., L.Z., C.R., and D.Y.H. provided suggestions for structuring the paper. All the authors contributed to the writing and interpreting of the results and approved the final manuscript.

---

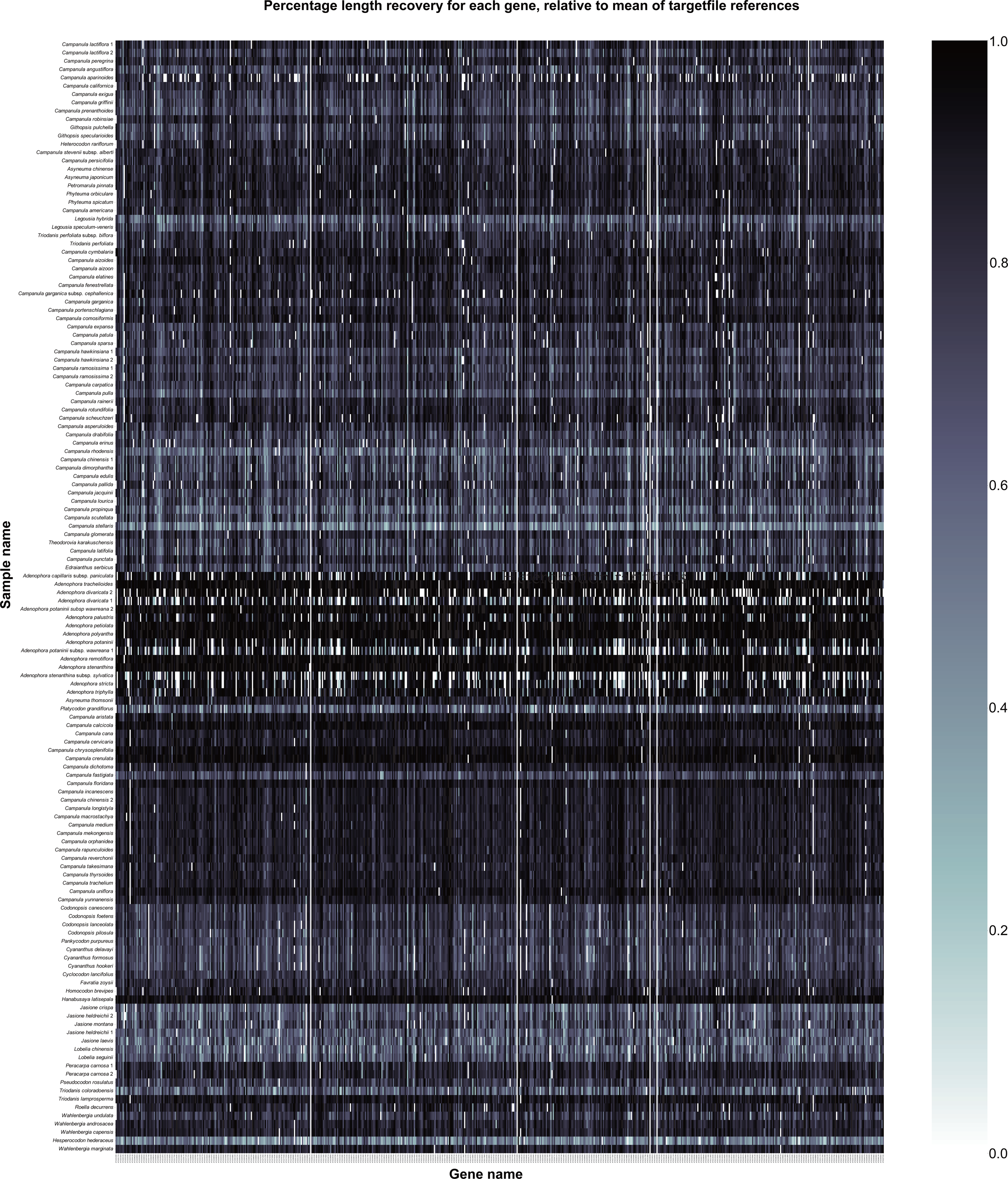

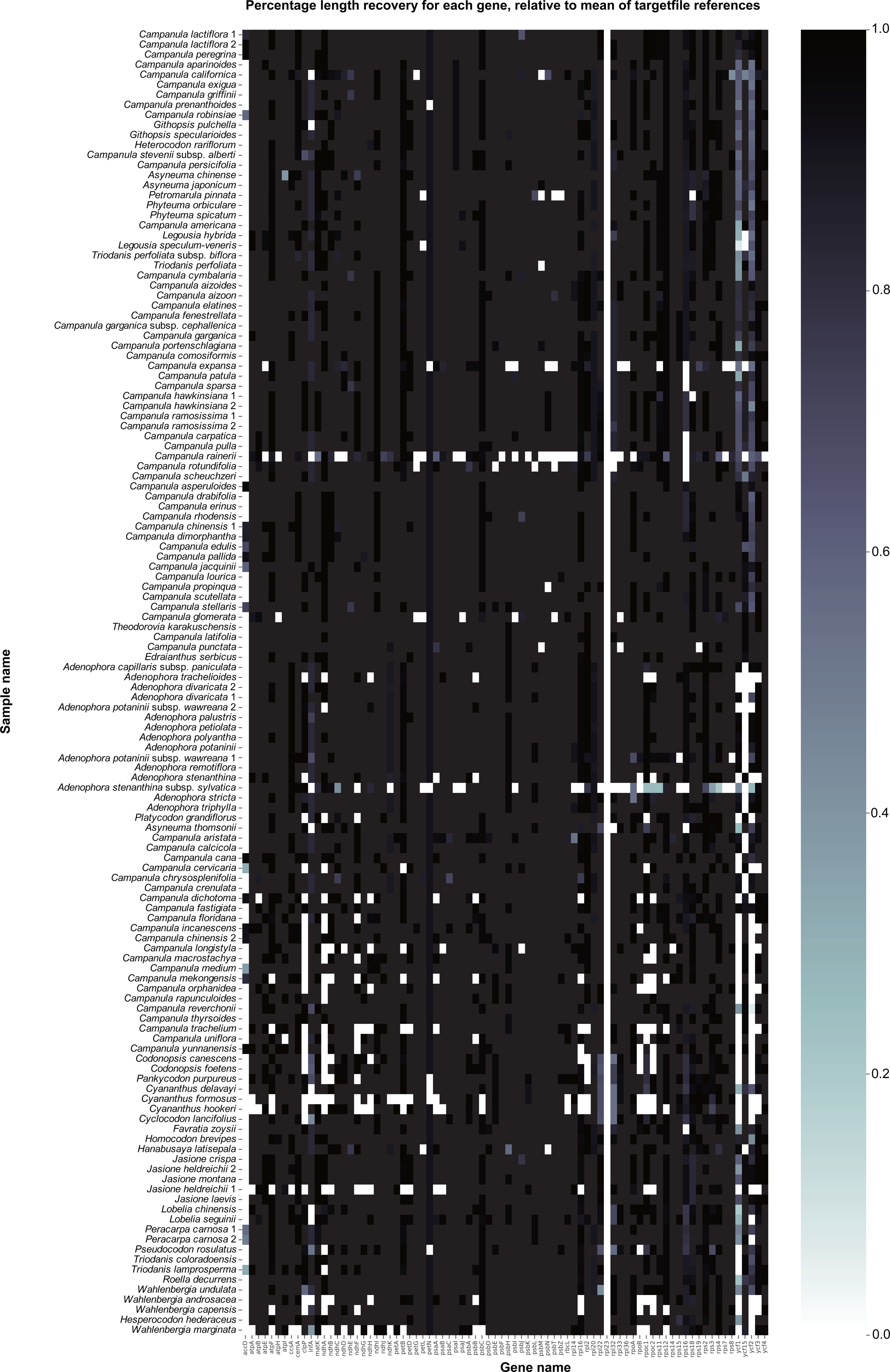

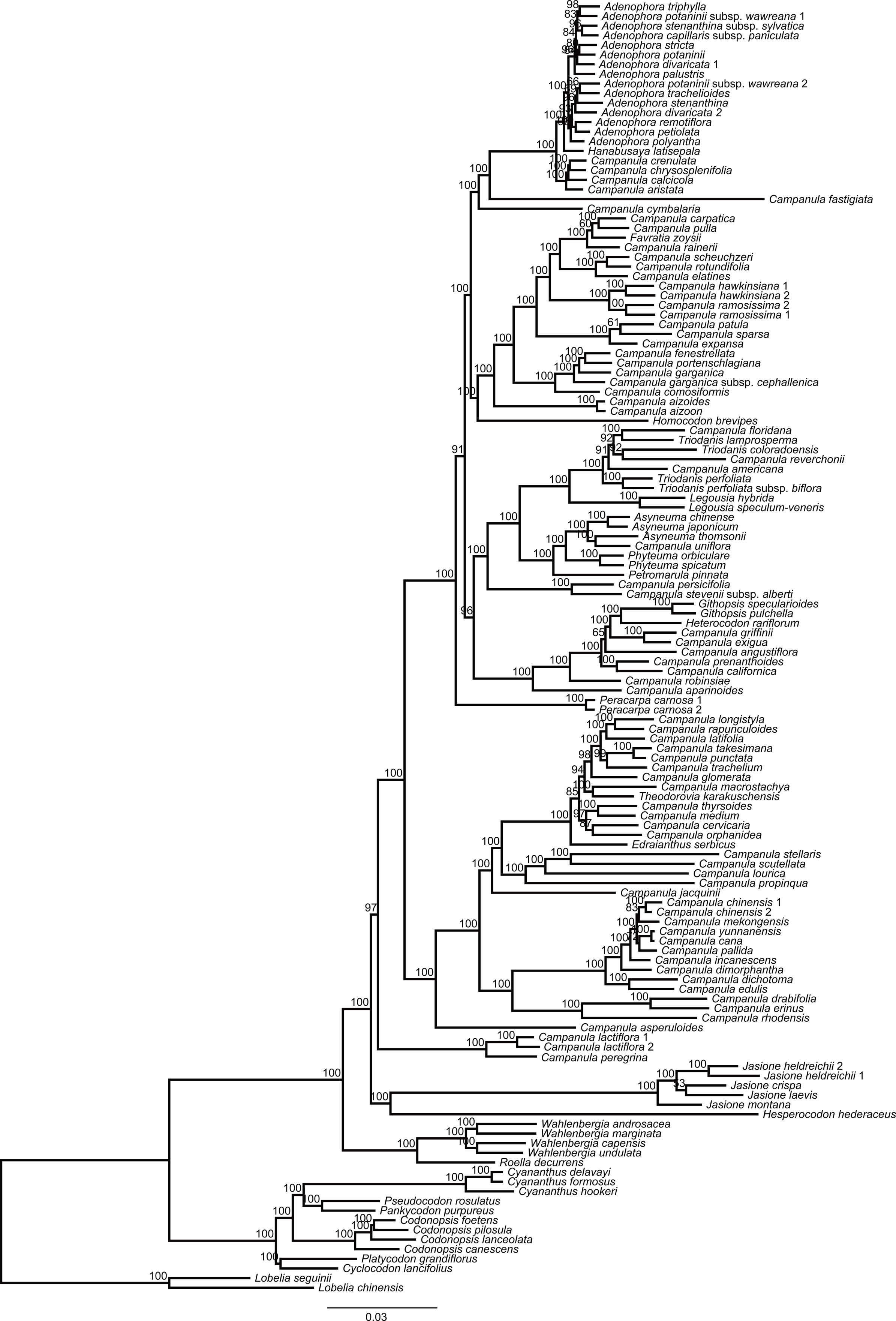

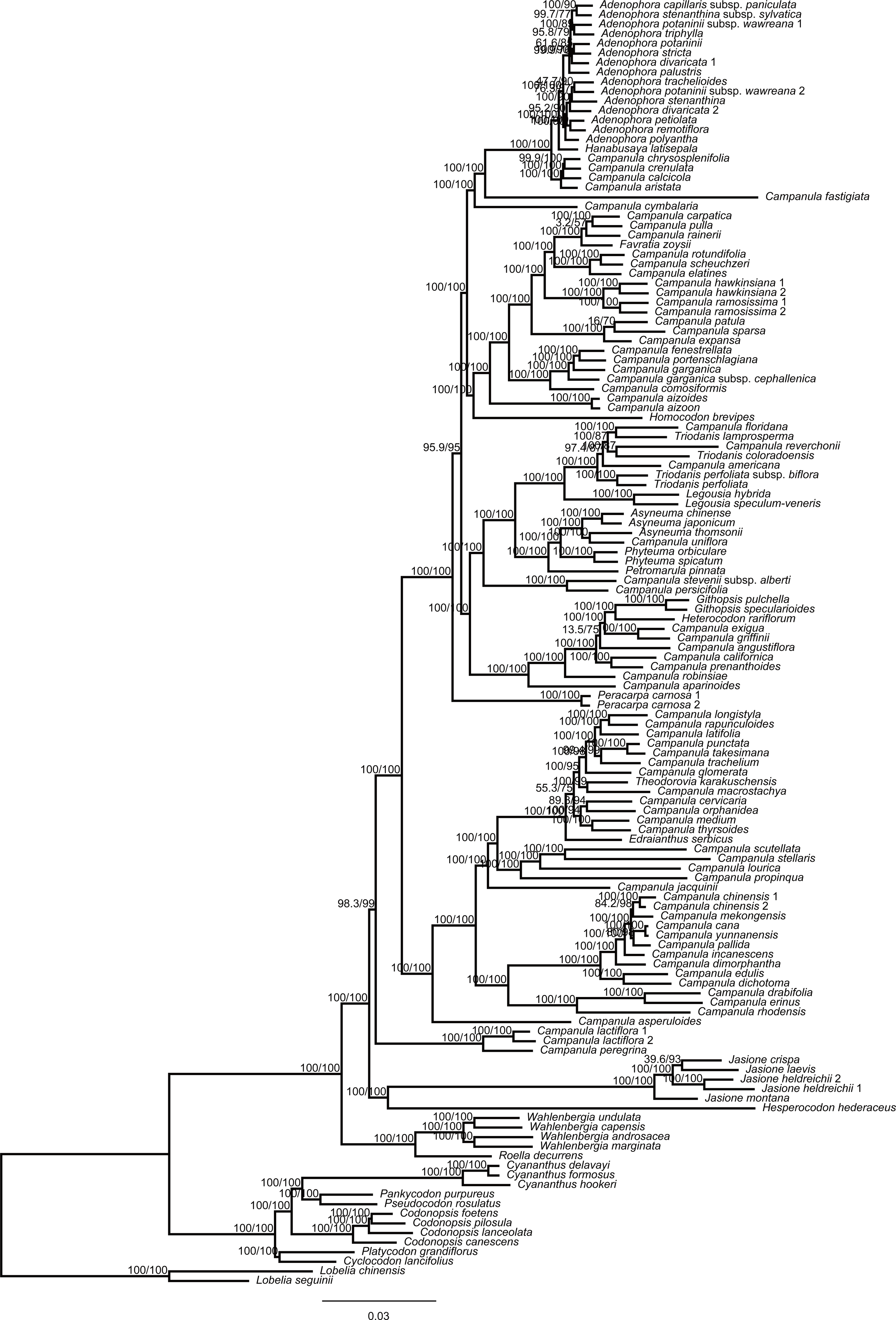

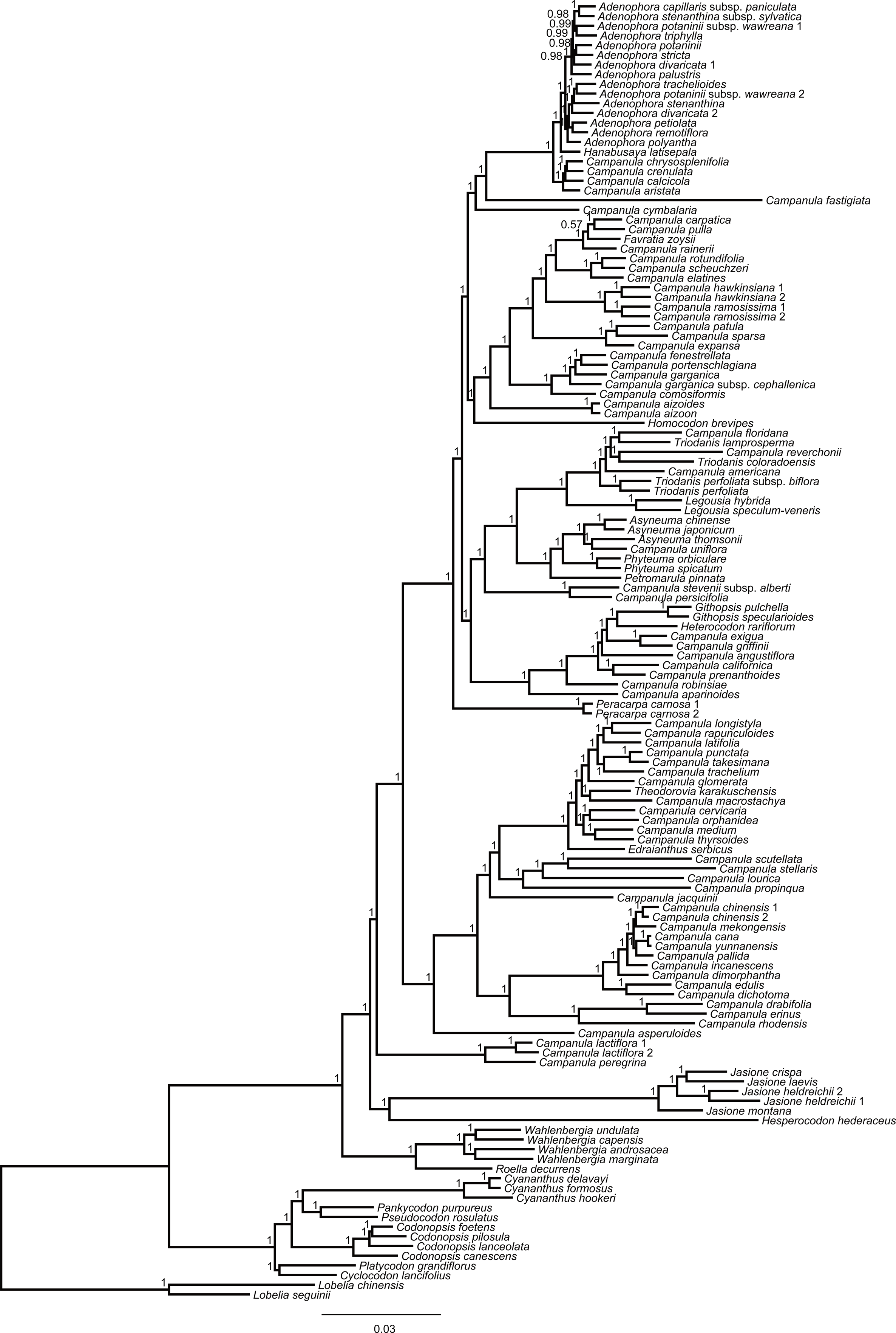

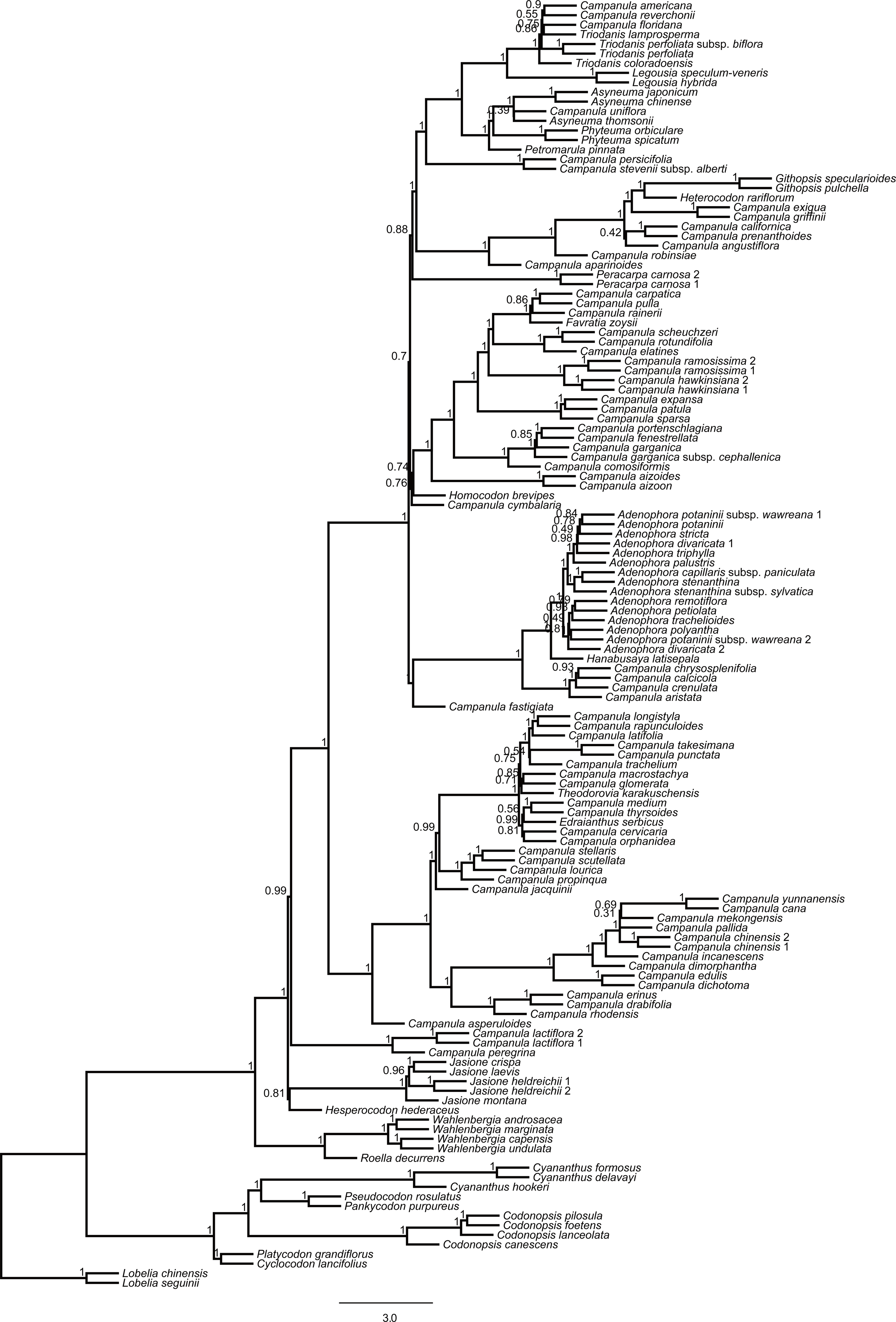

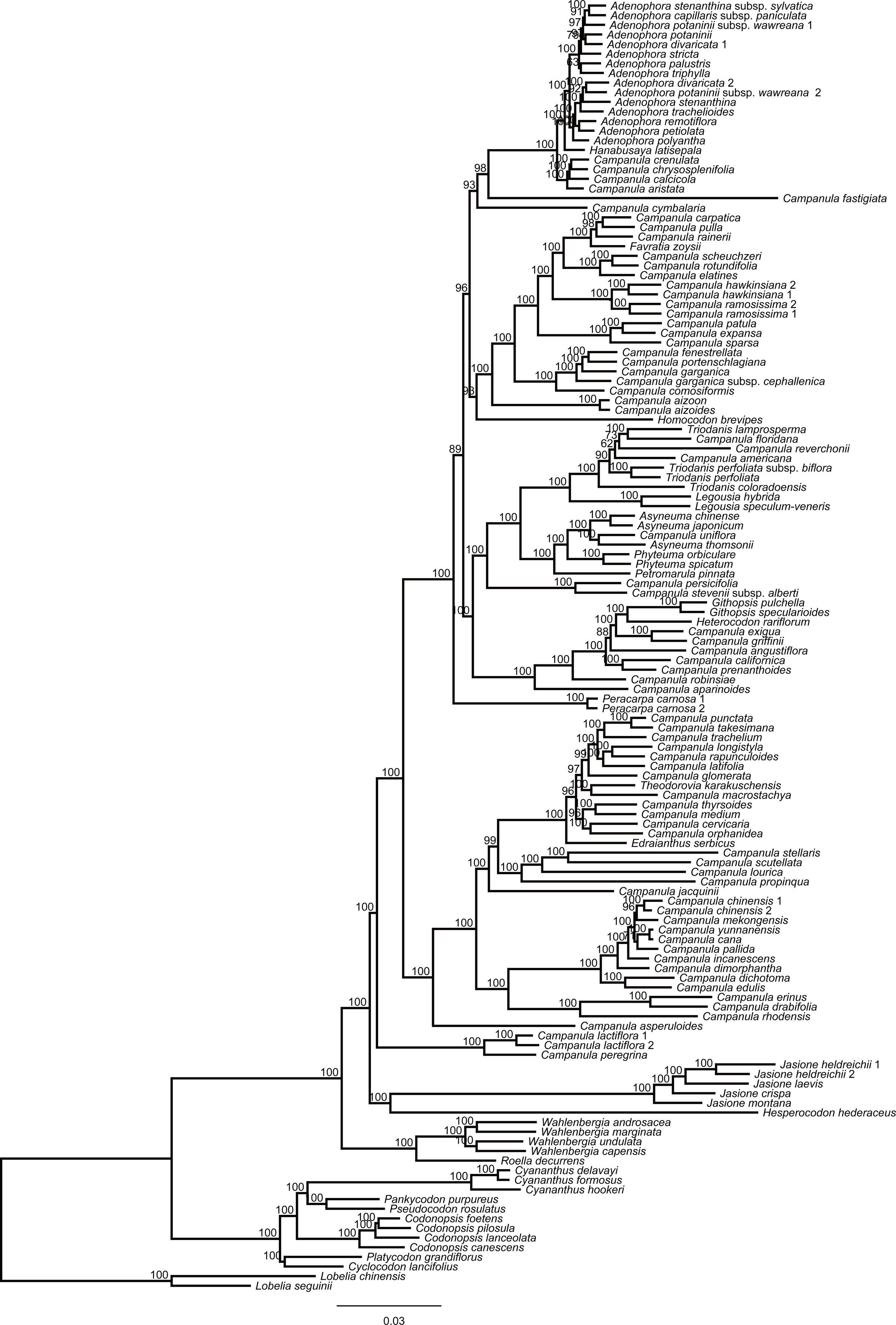

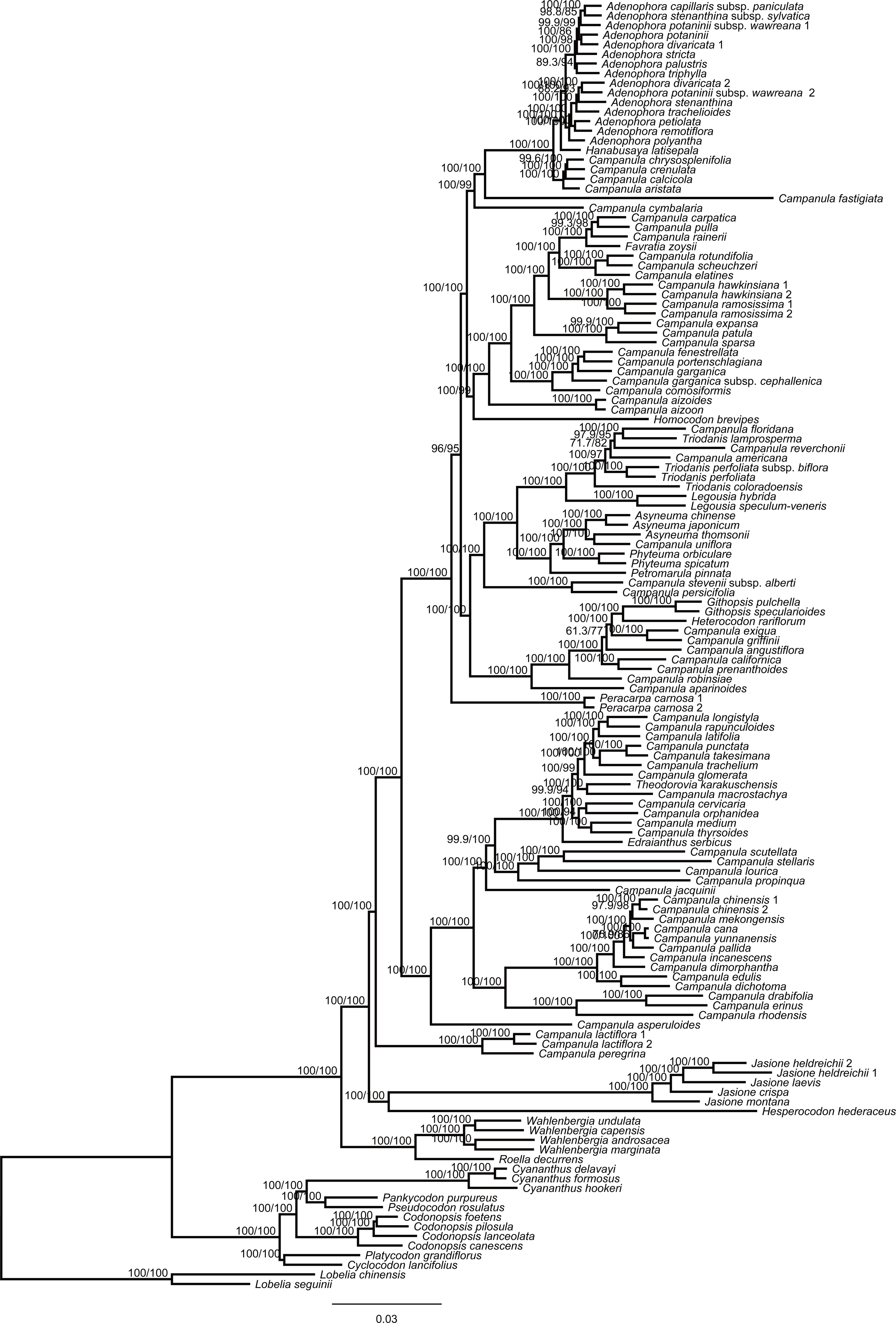

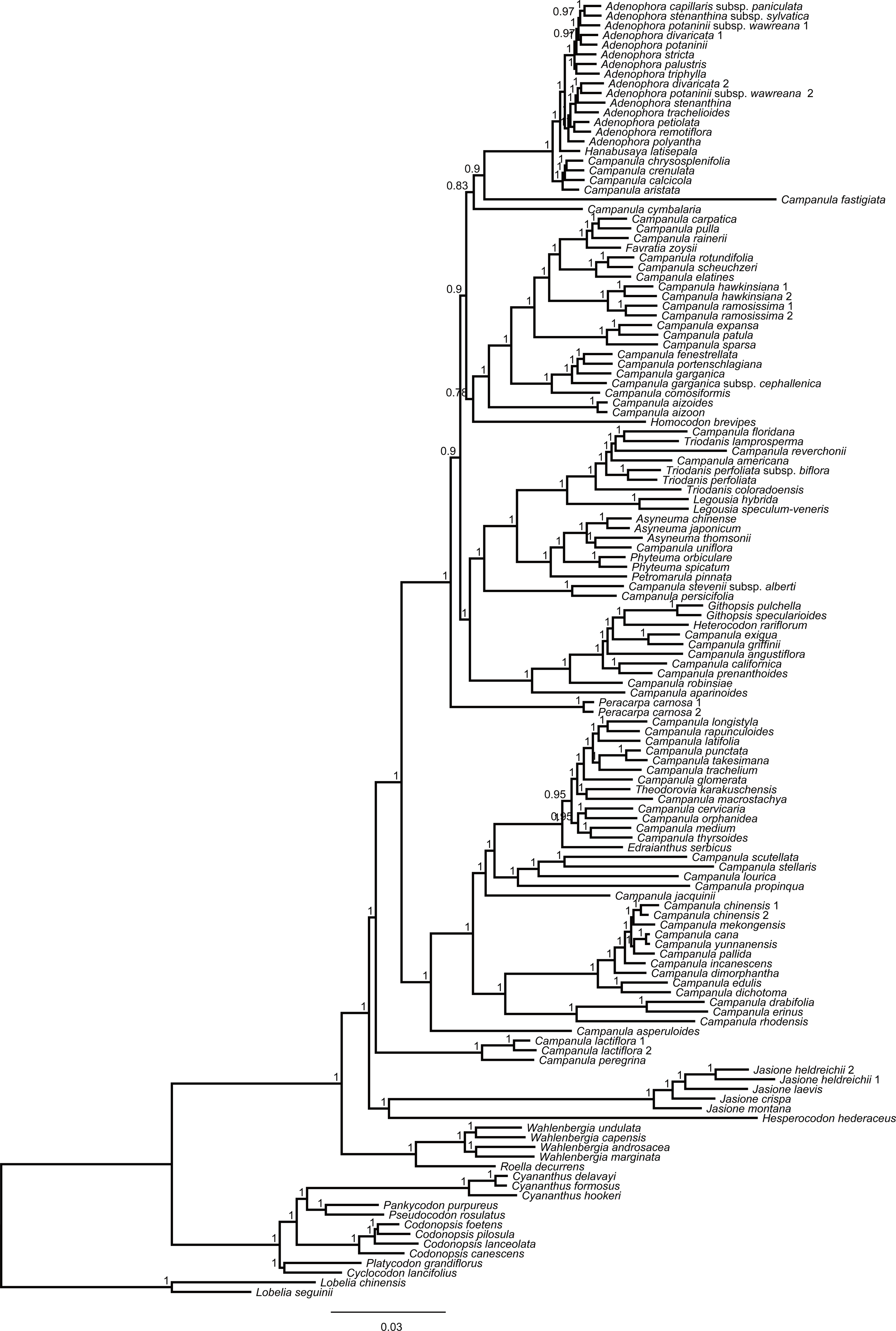

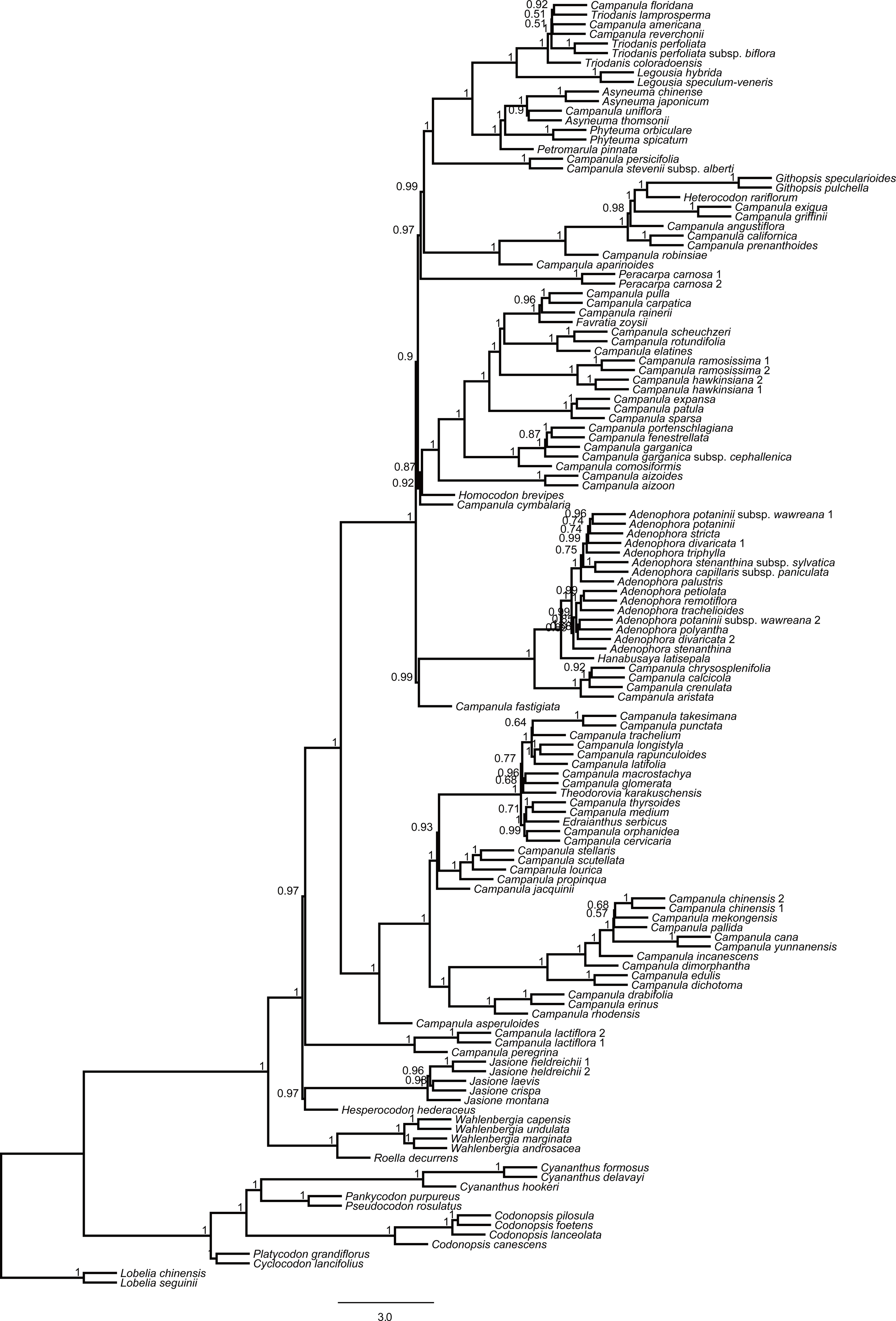

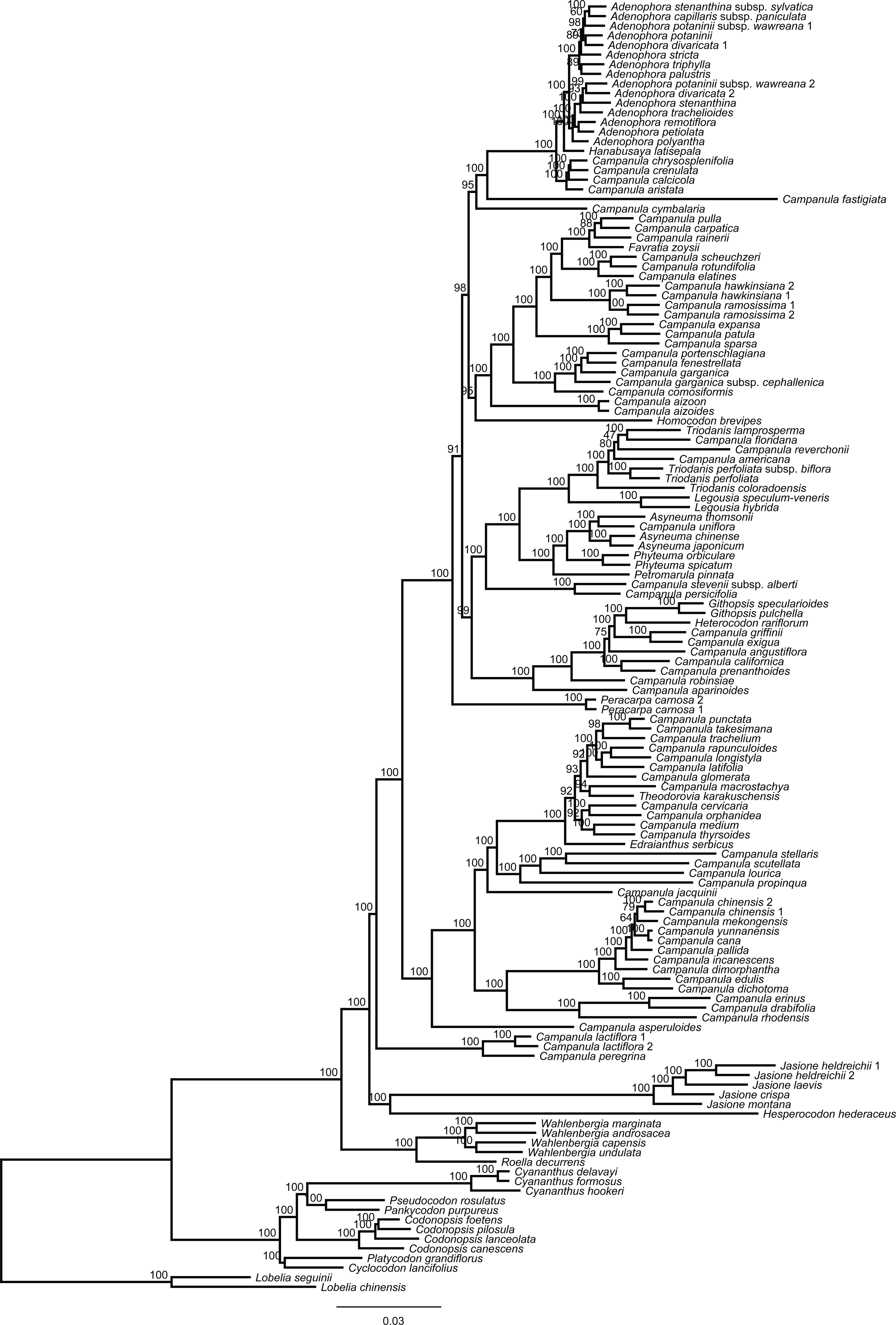

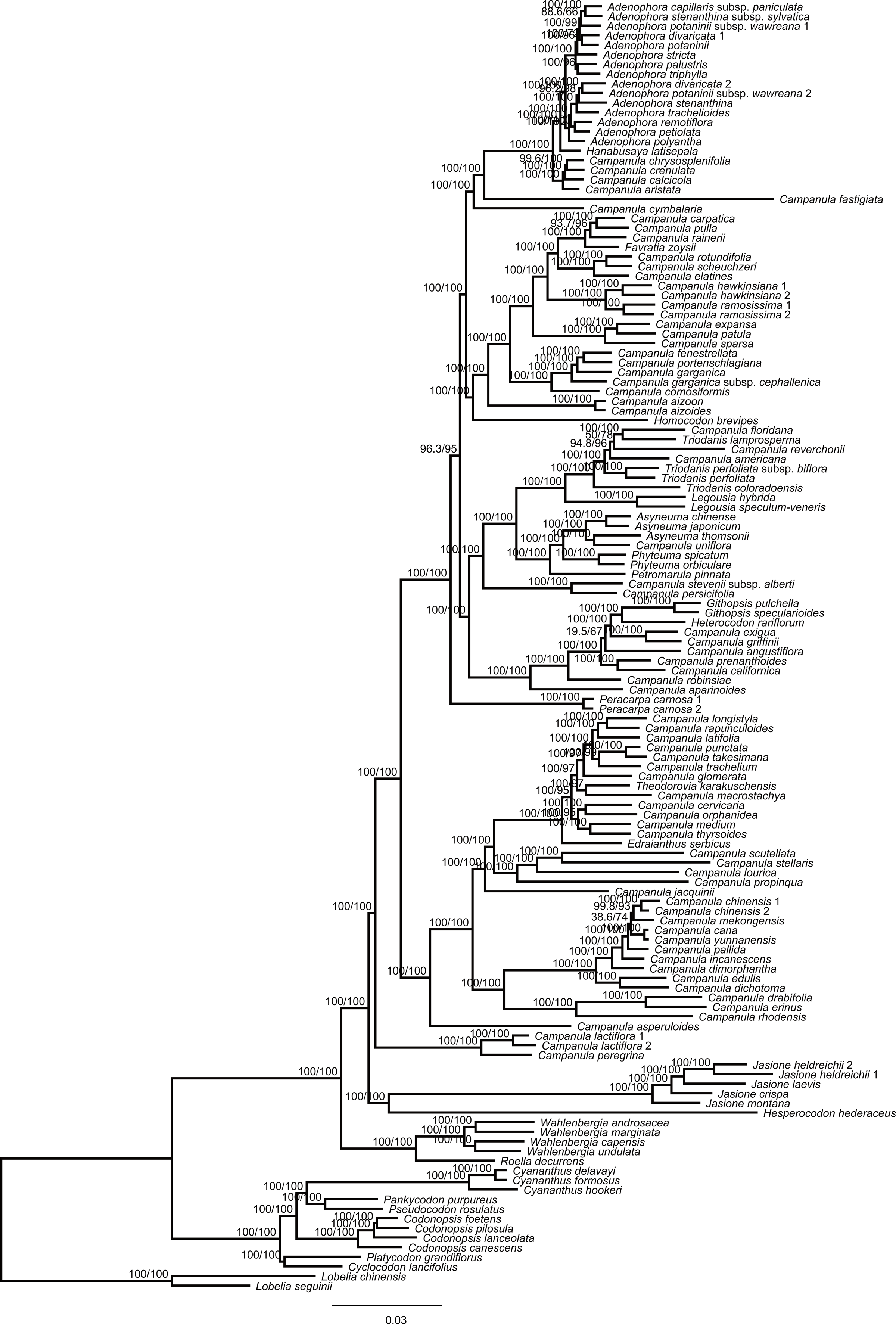

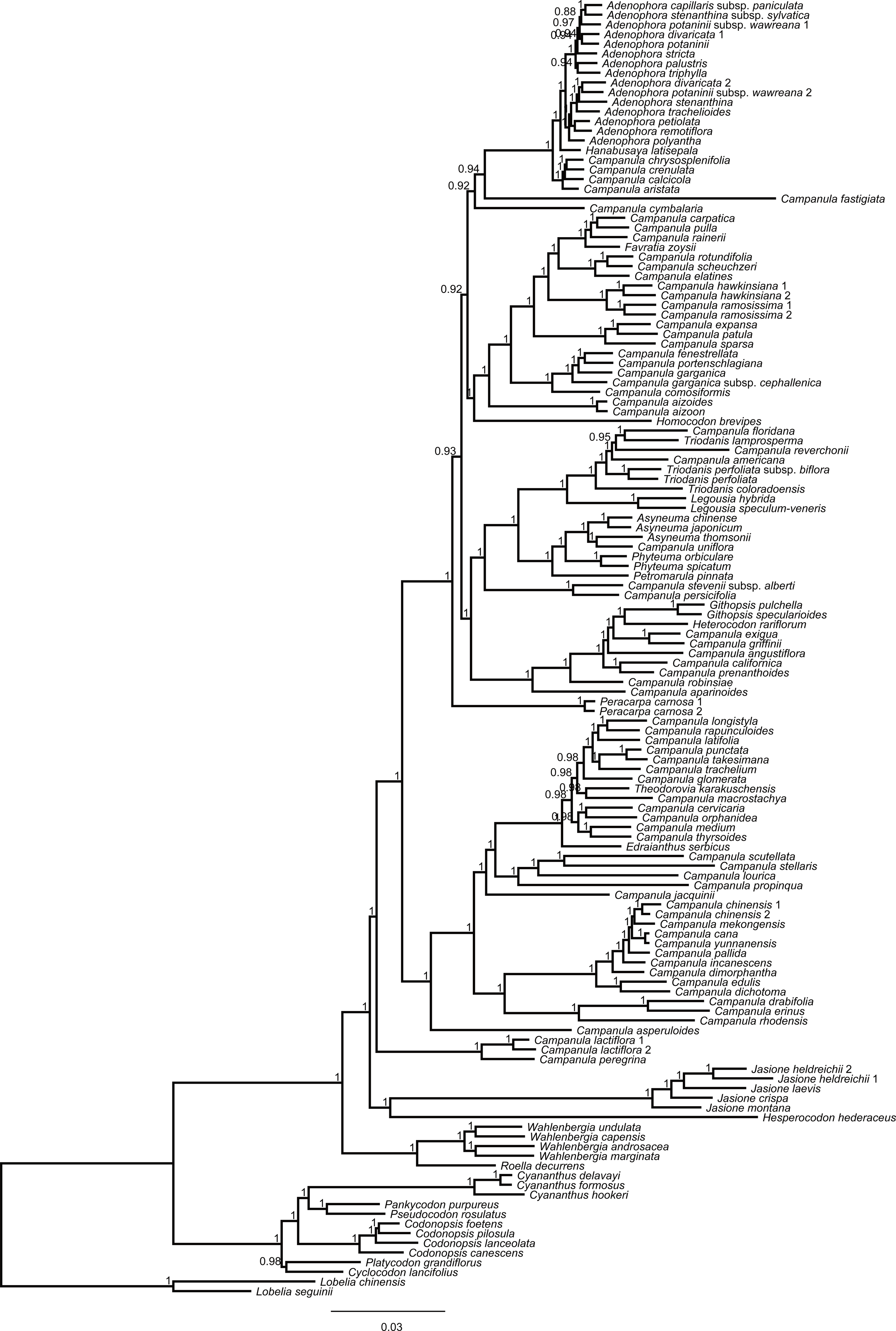

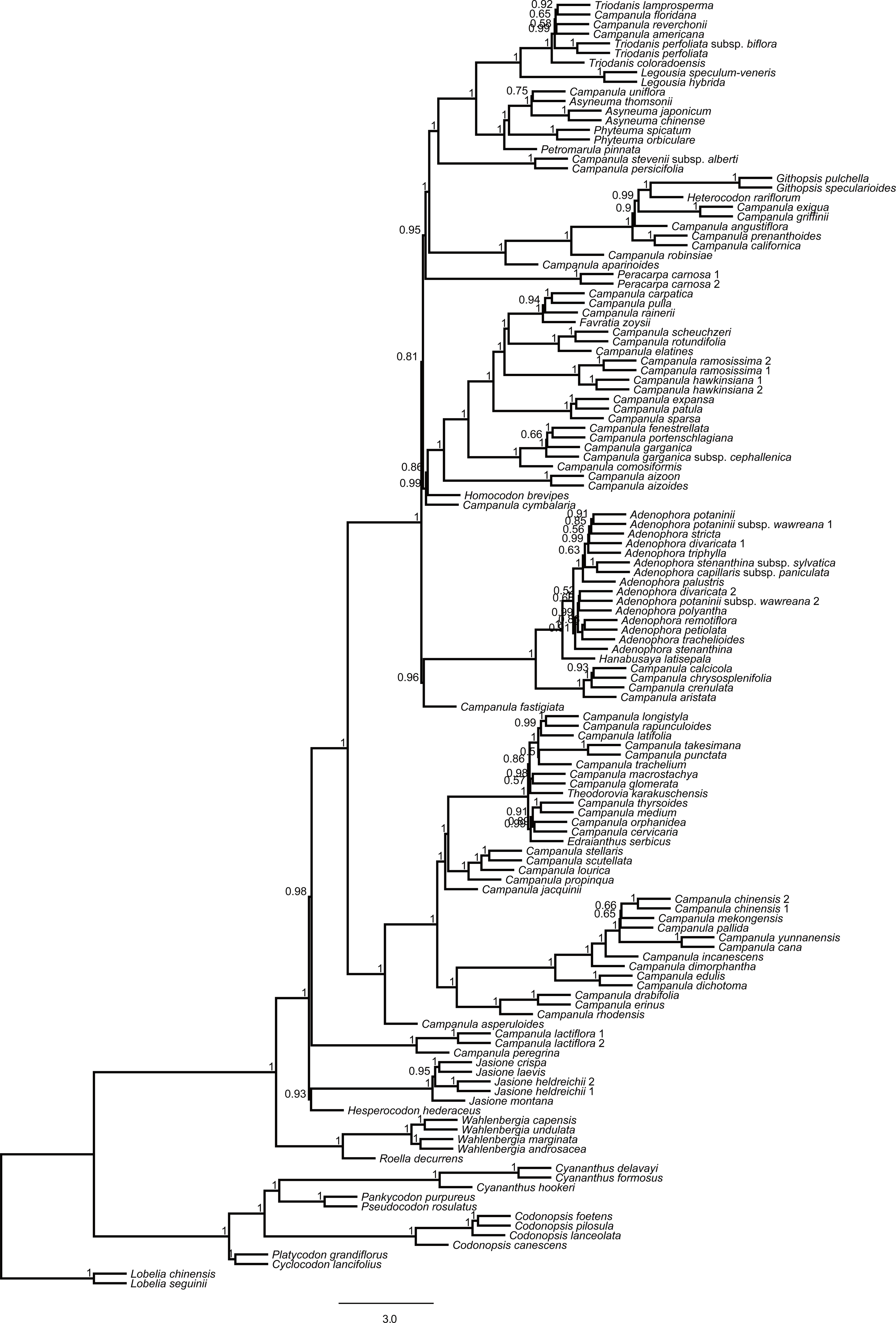

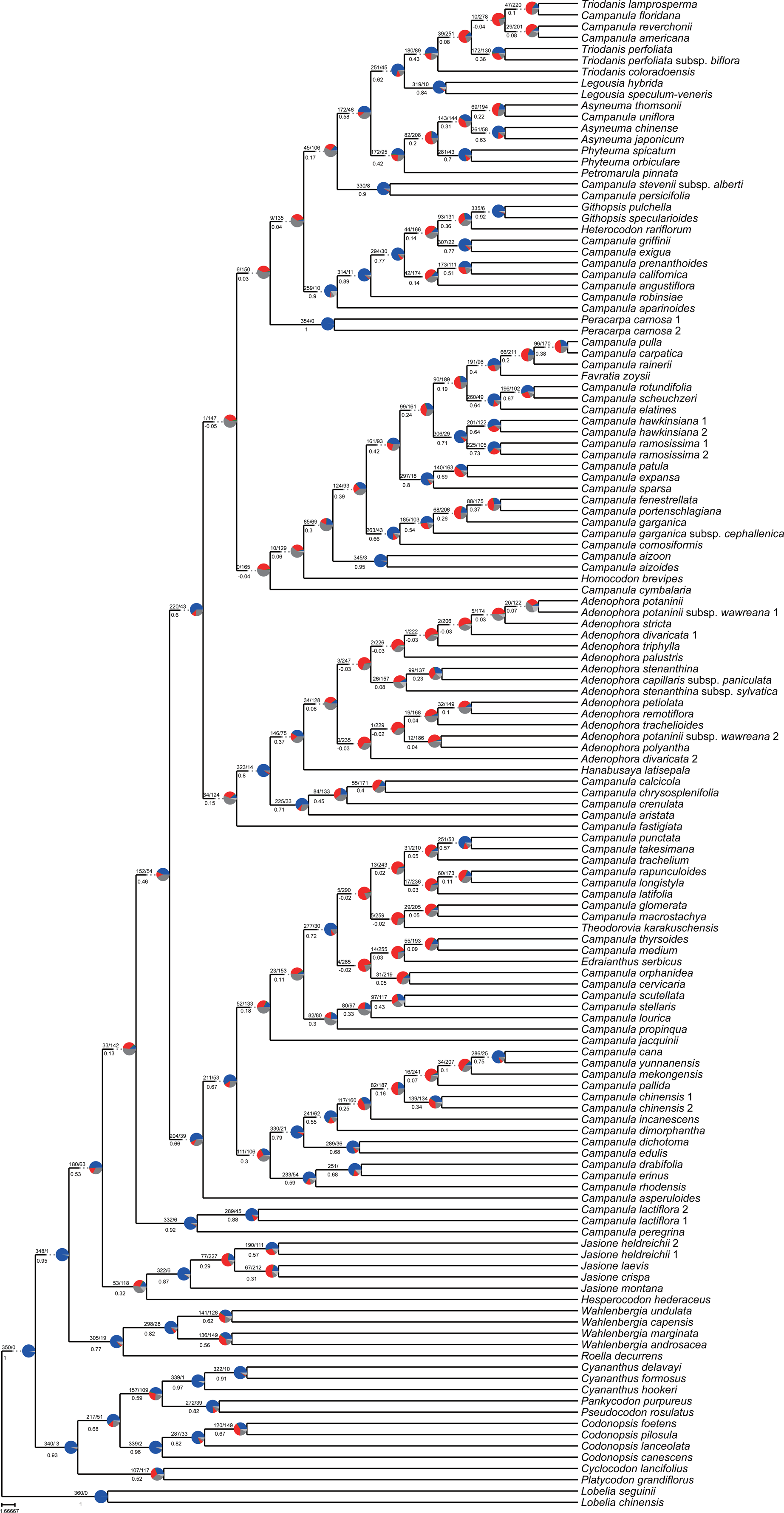

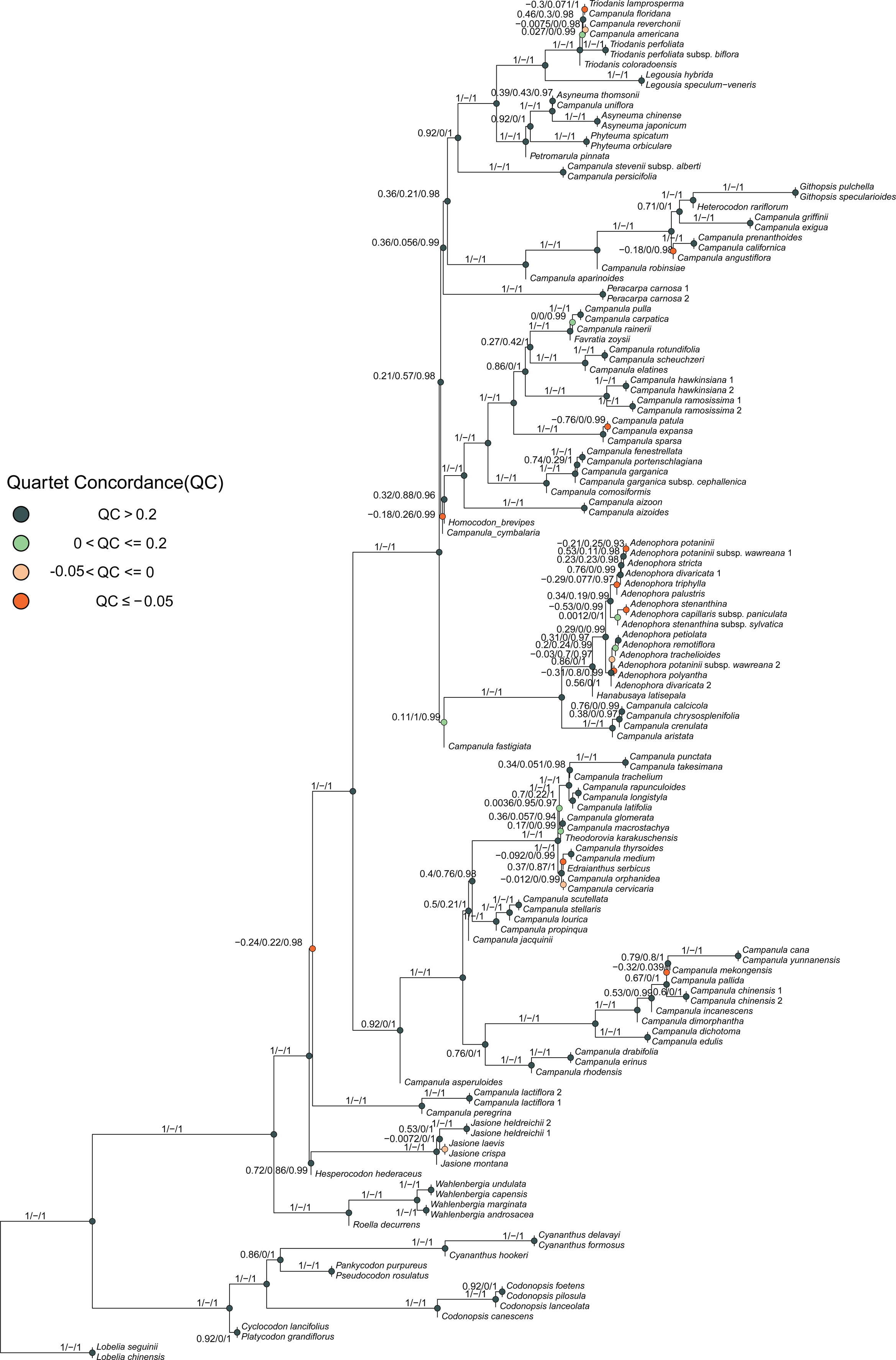

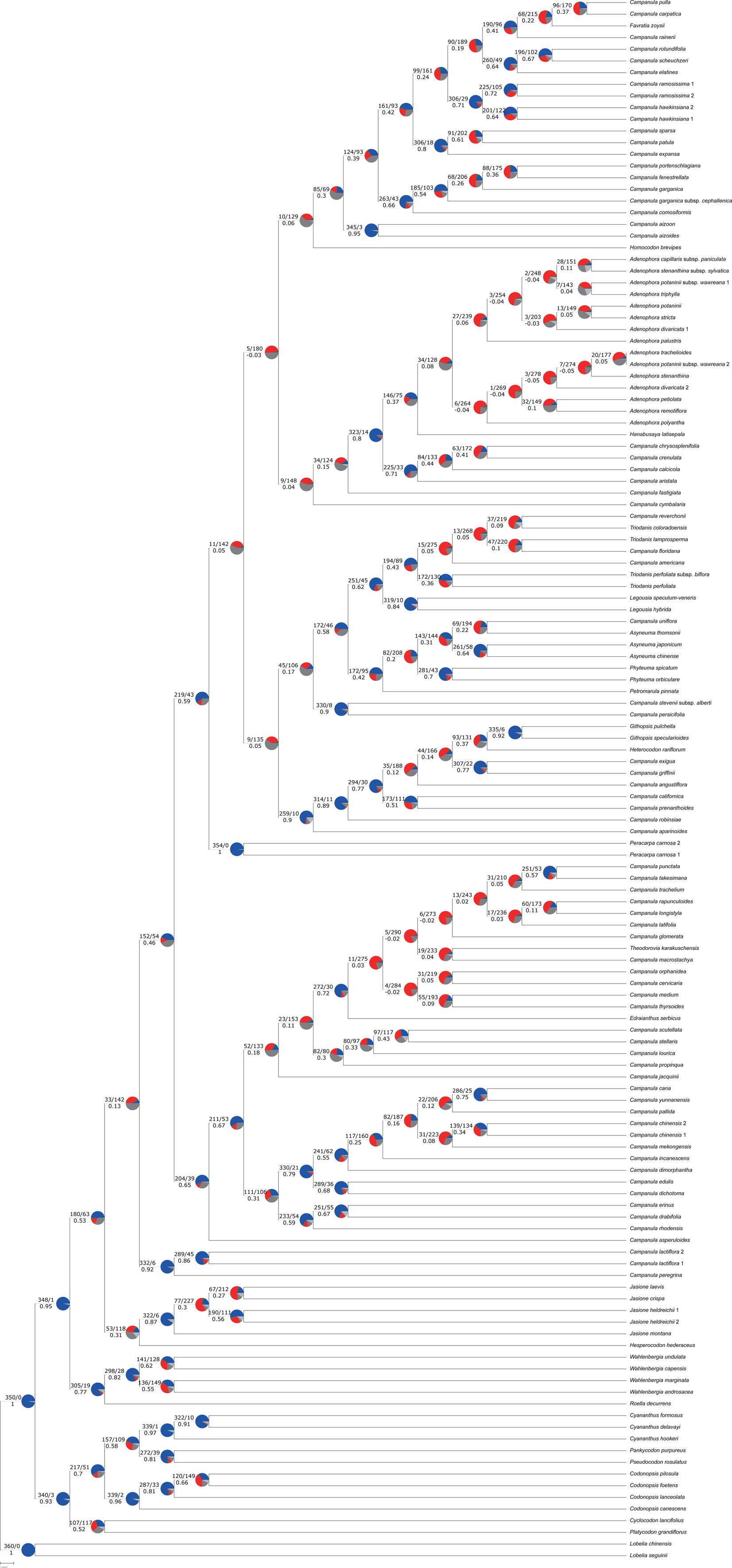

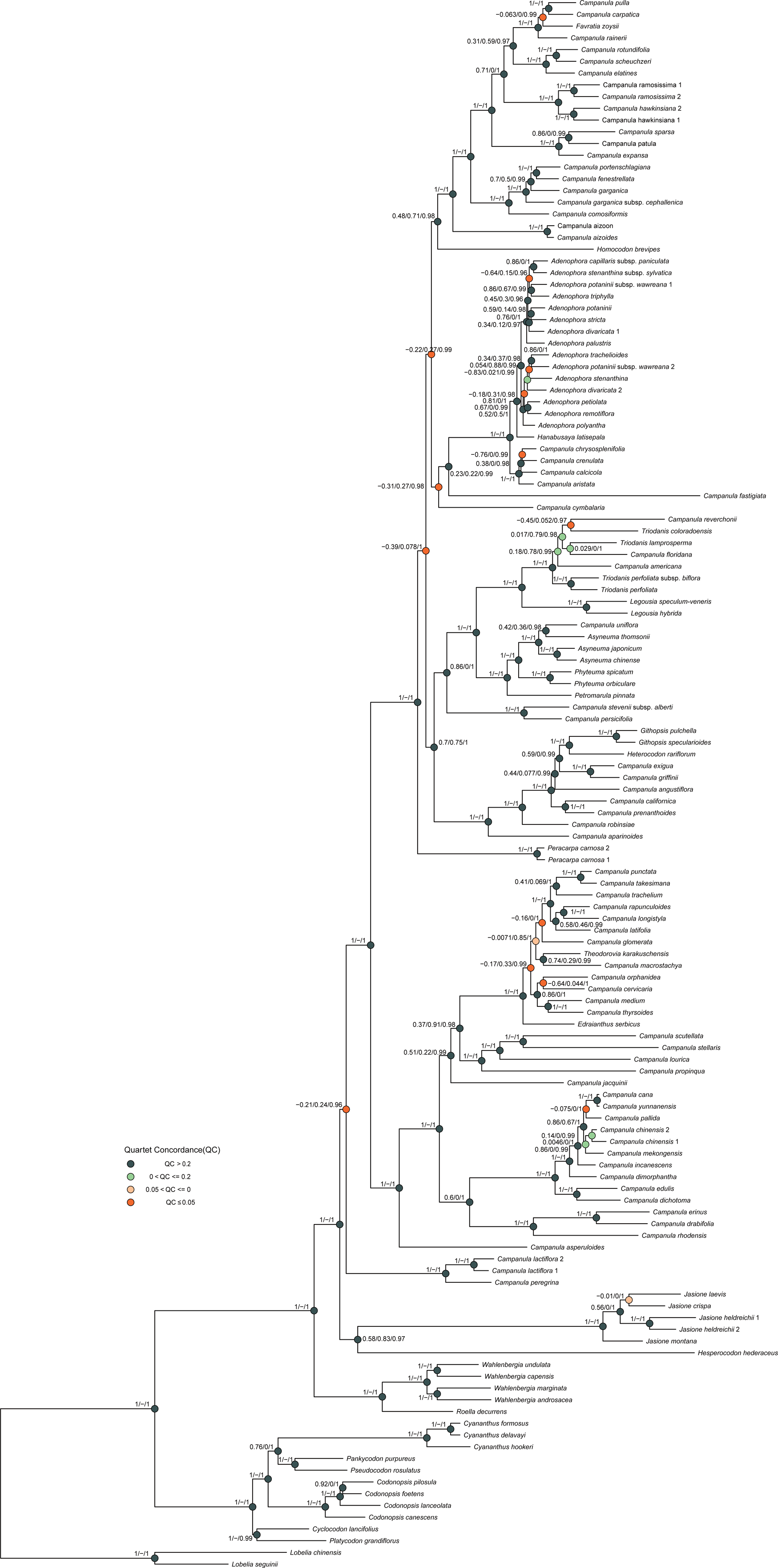

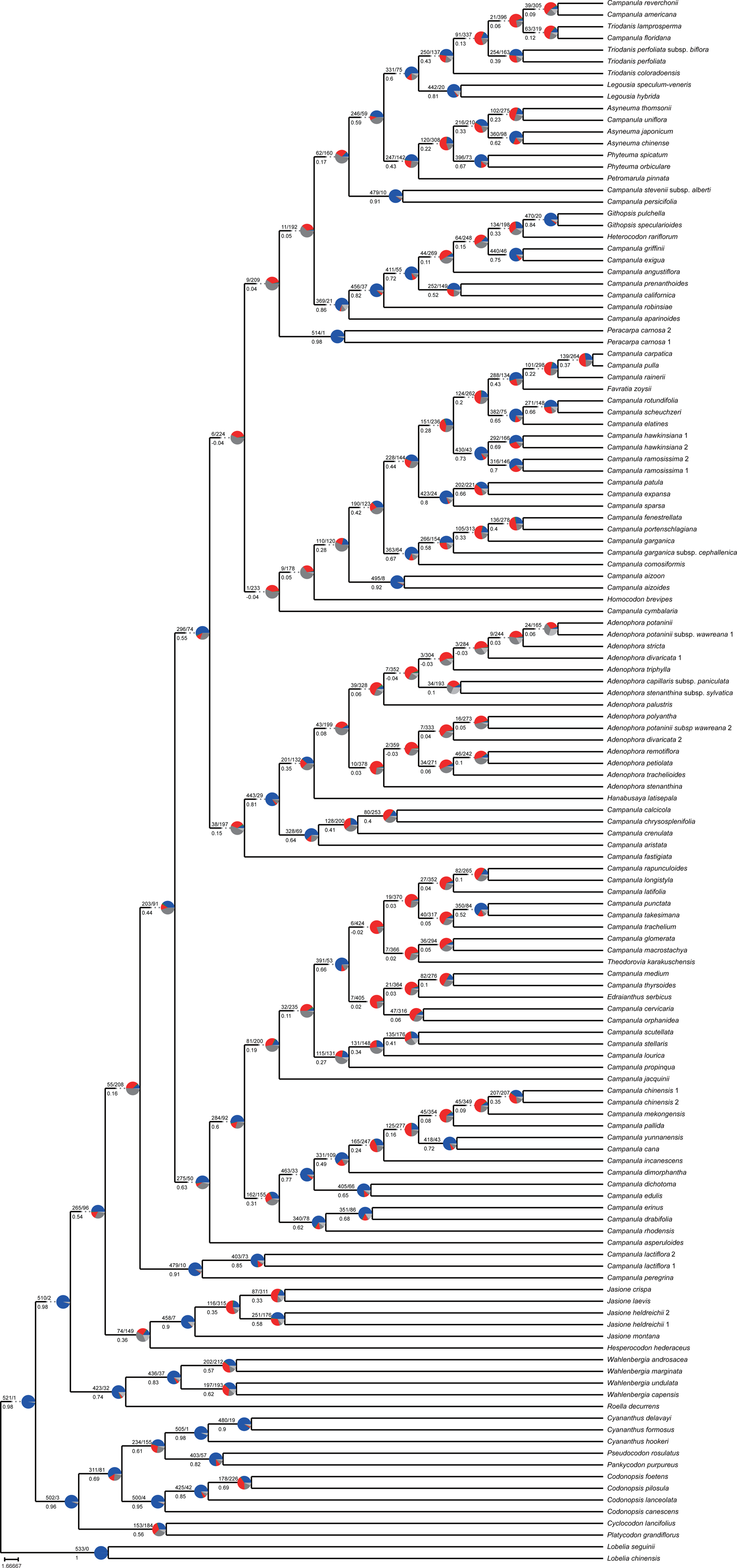

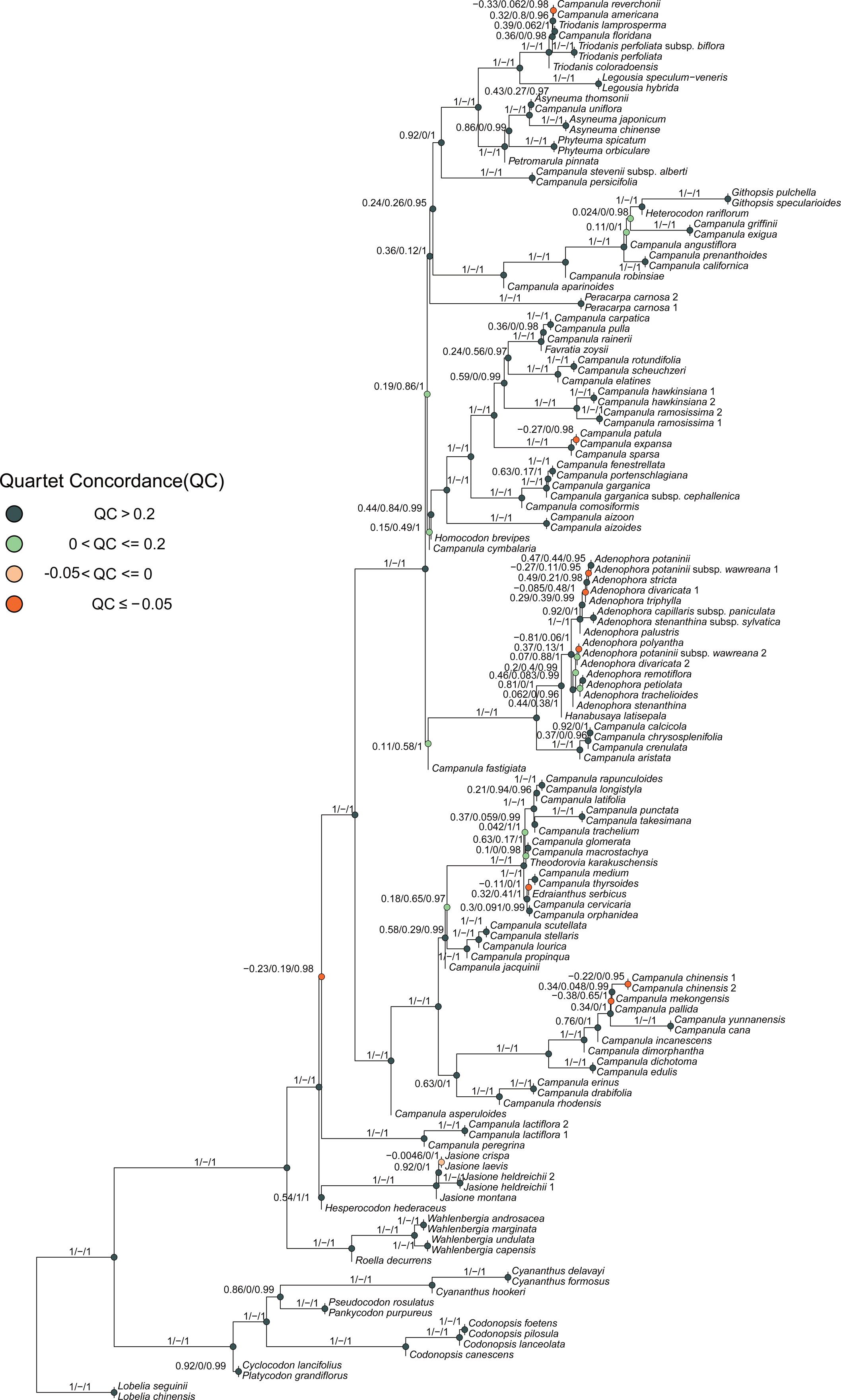

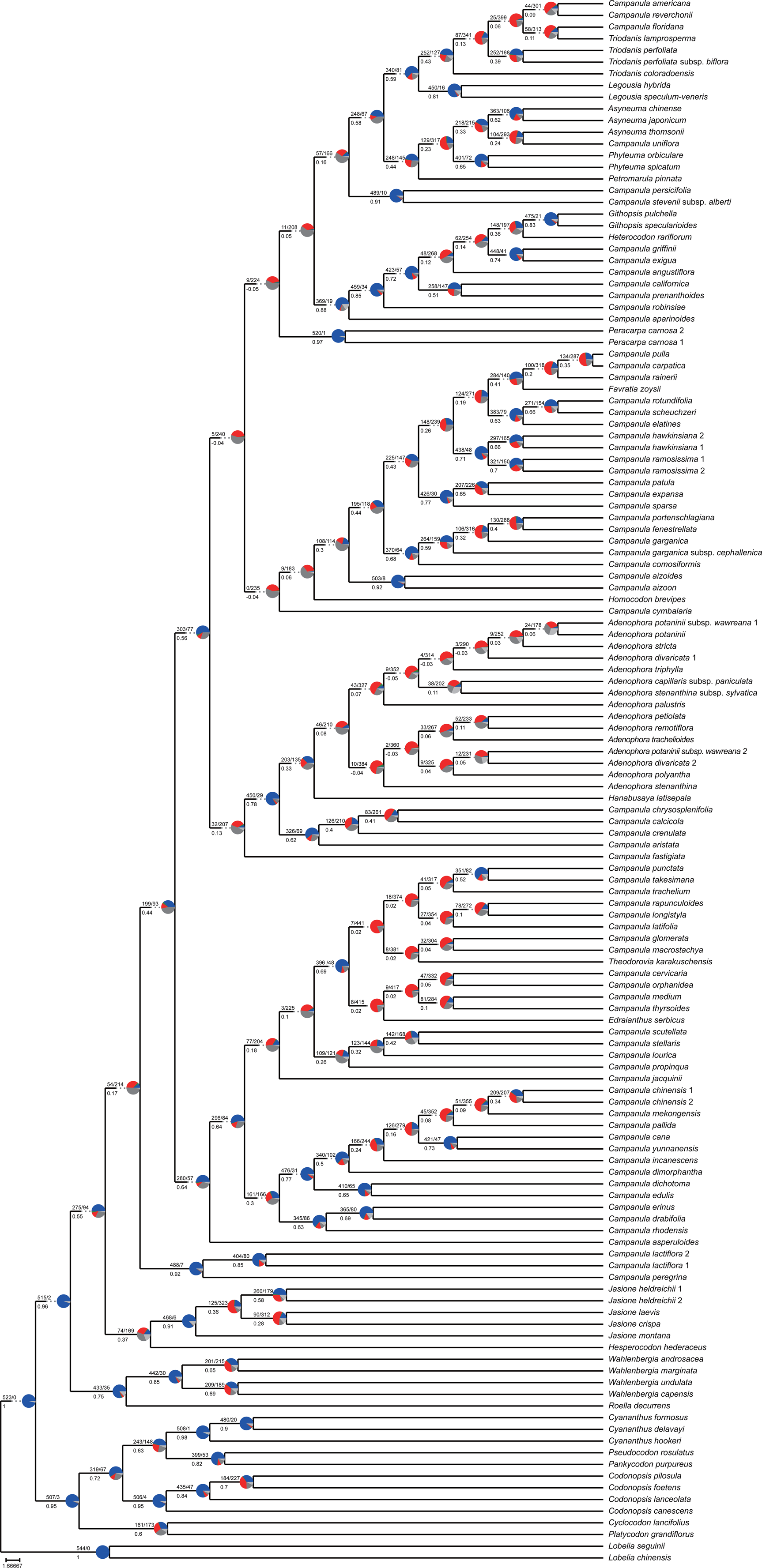

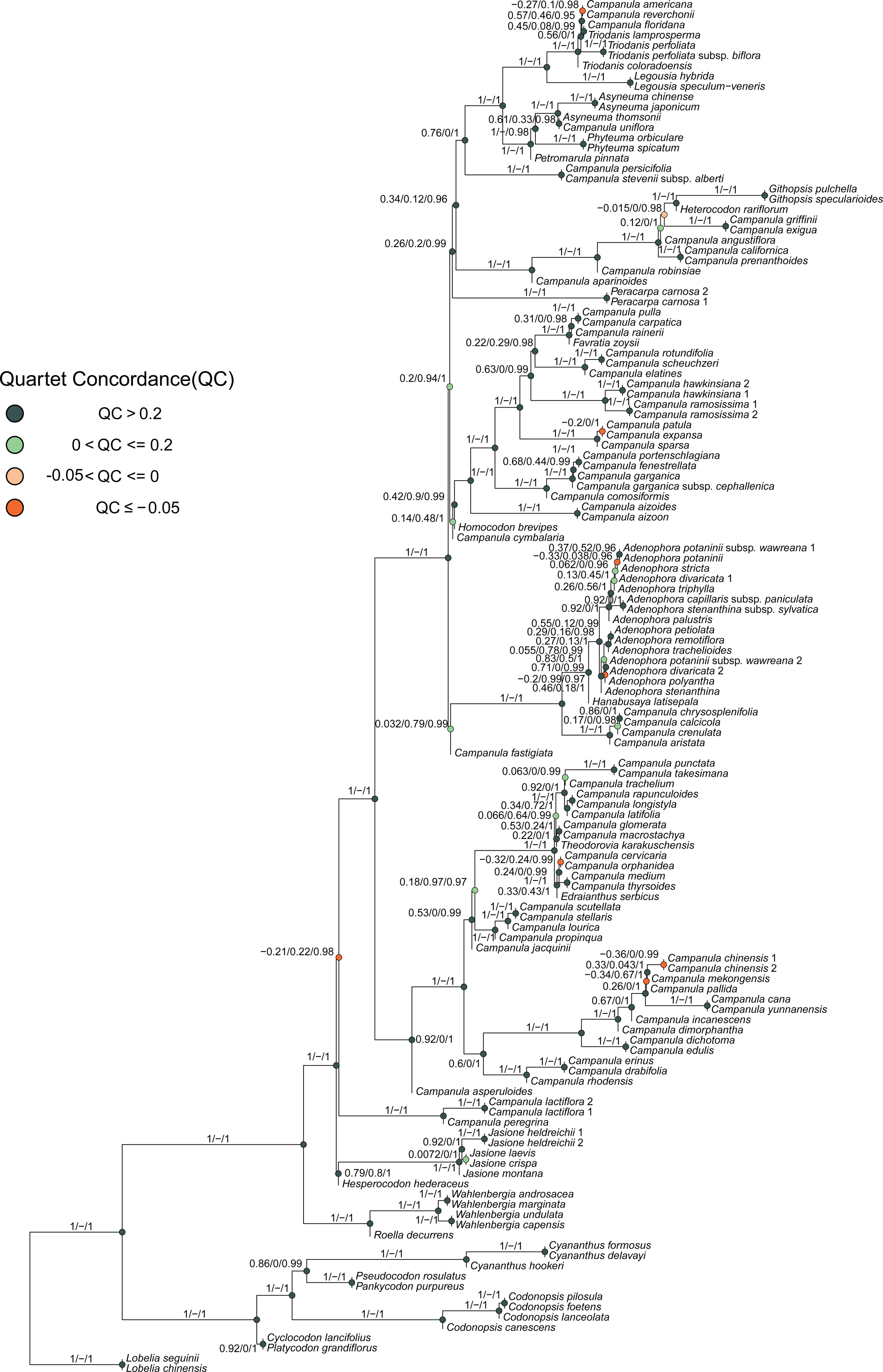

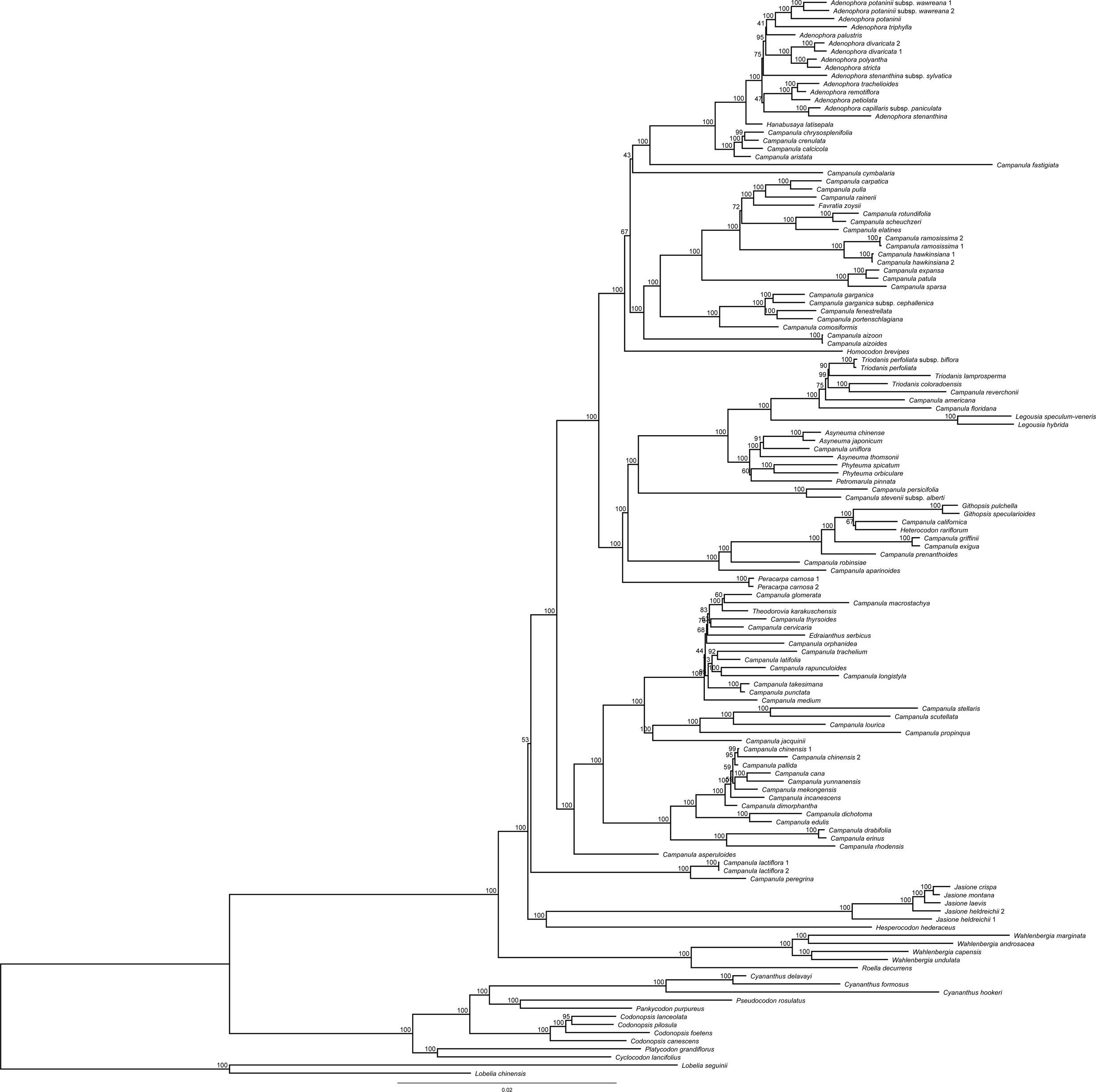

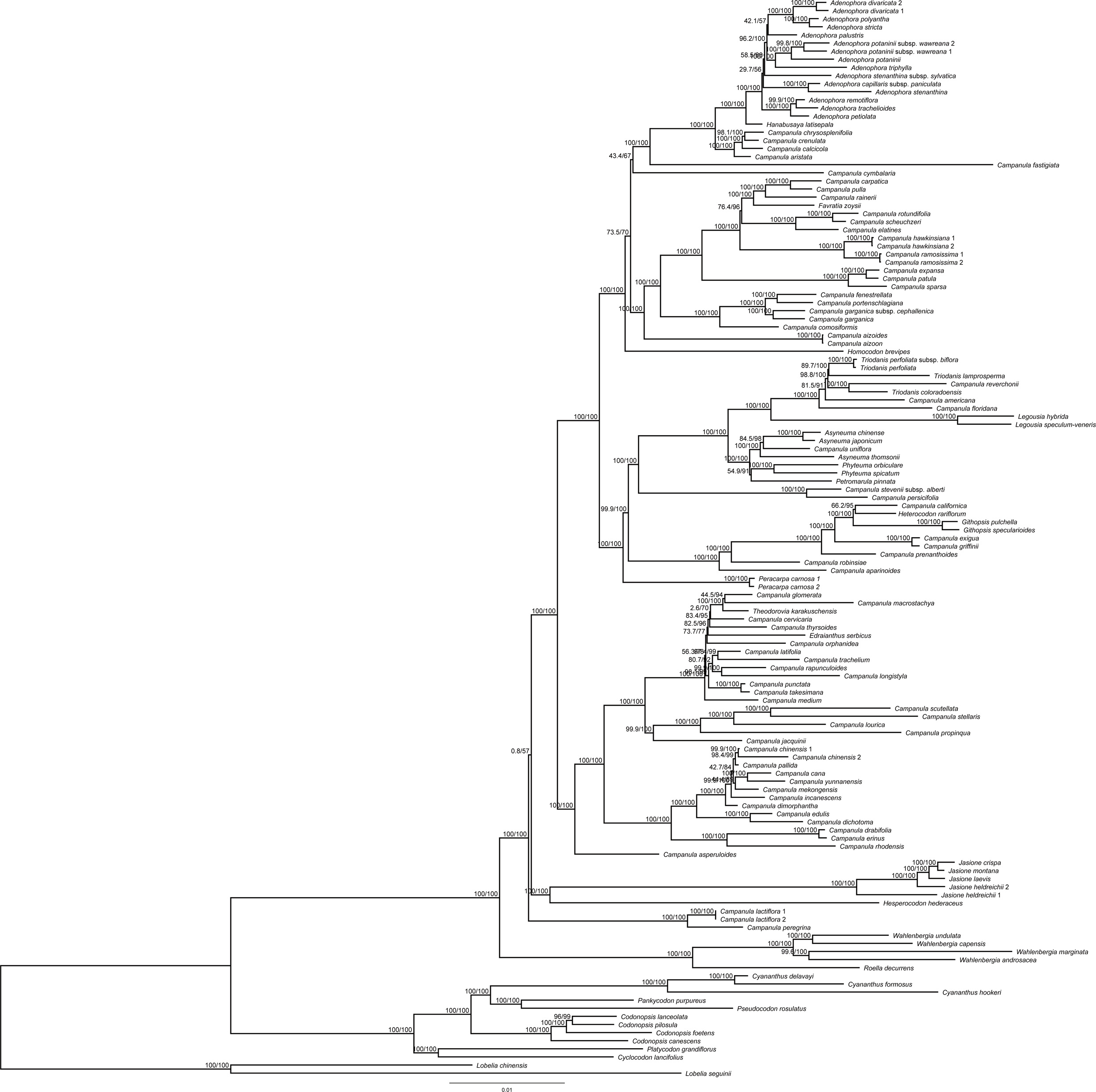

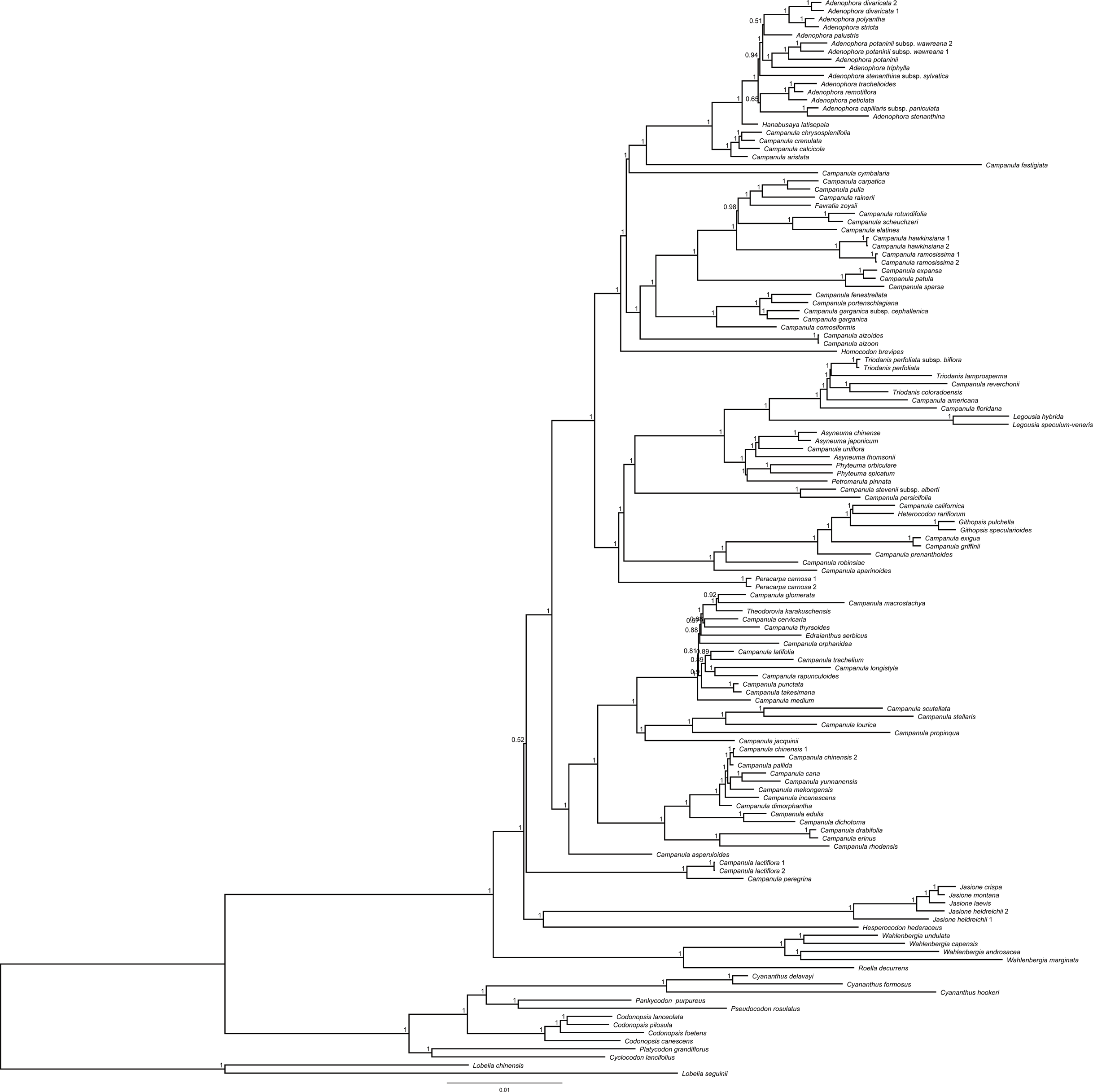

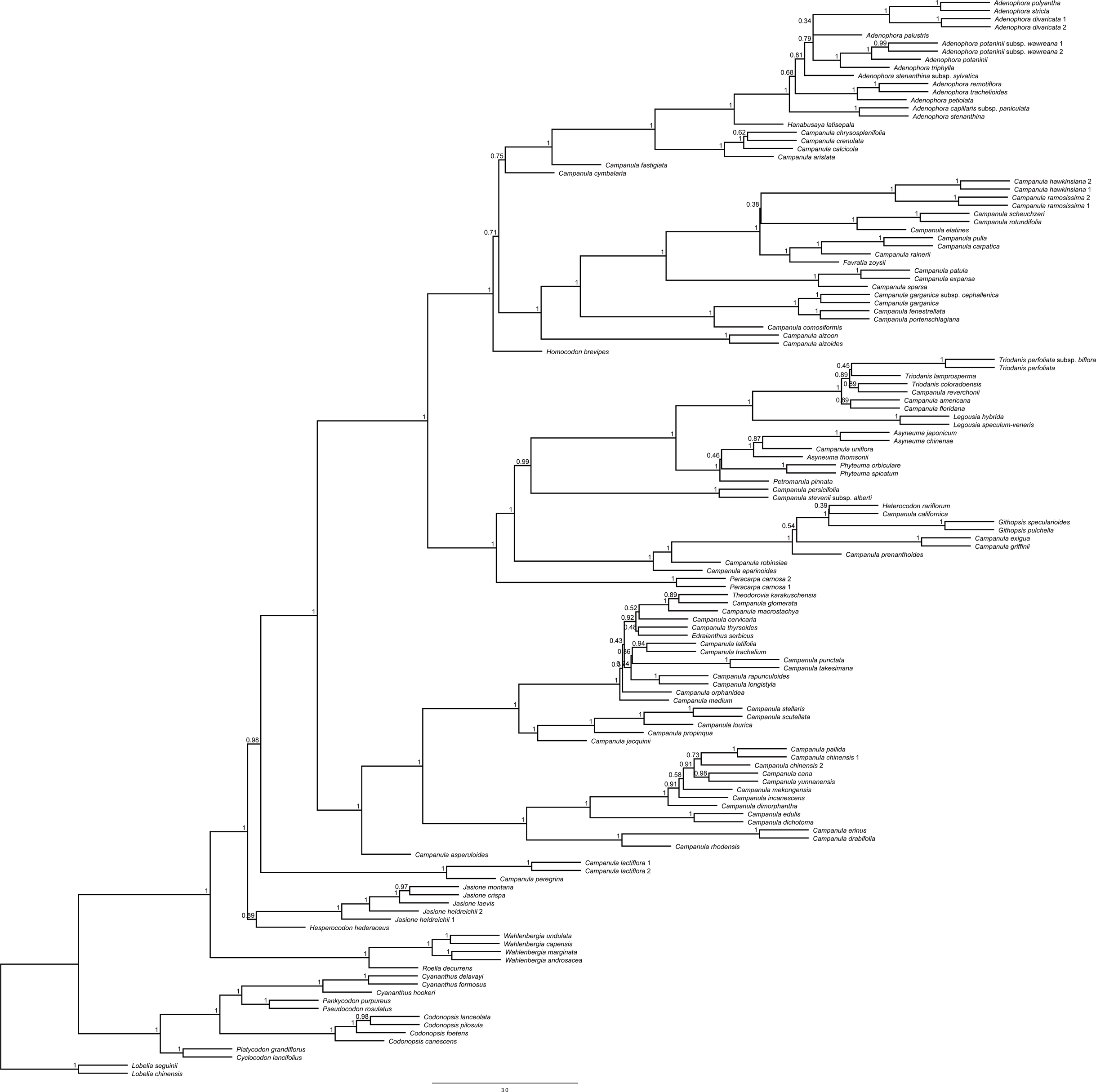

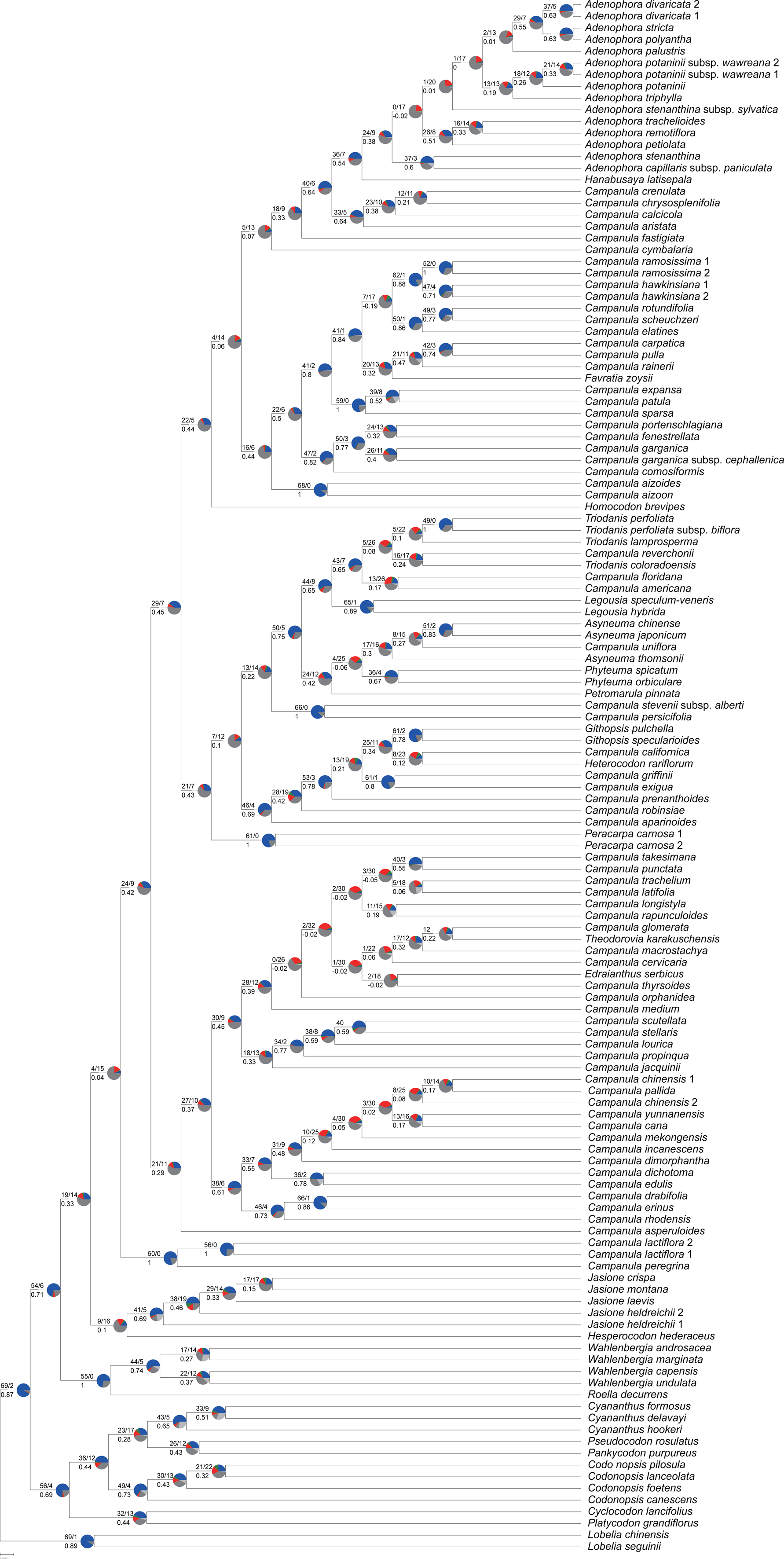

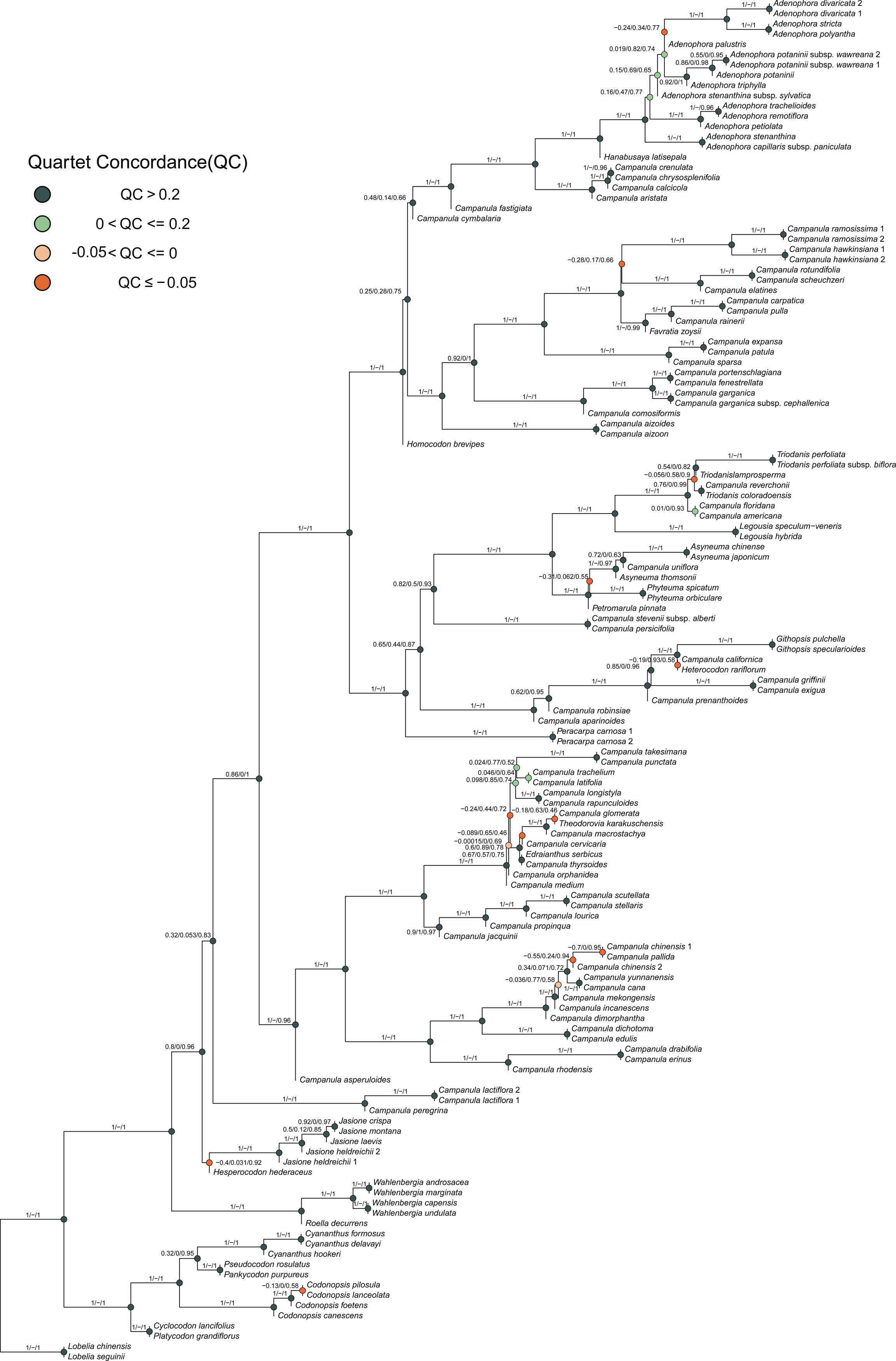

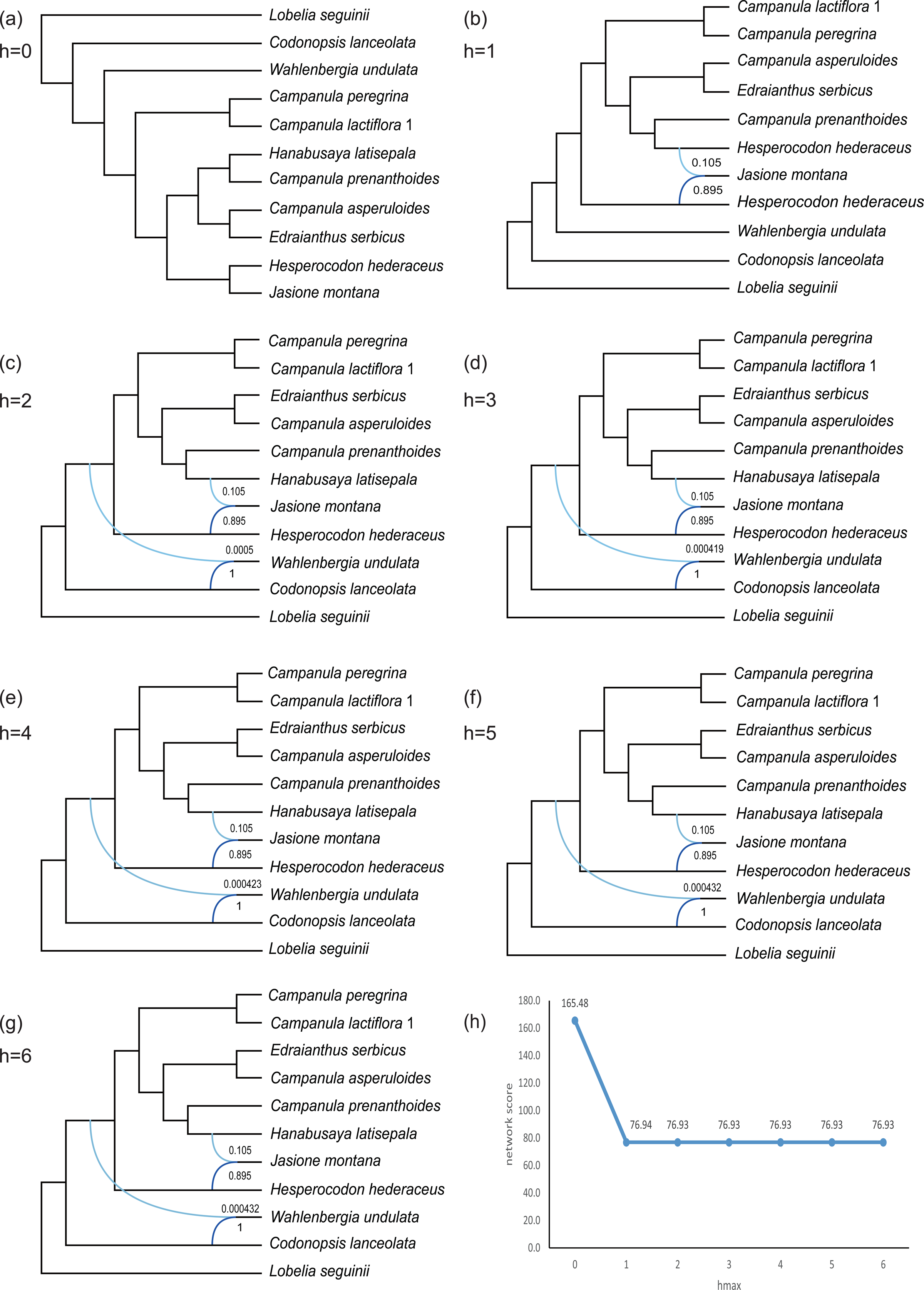

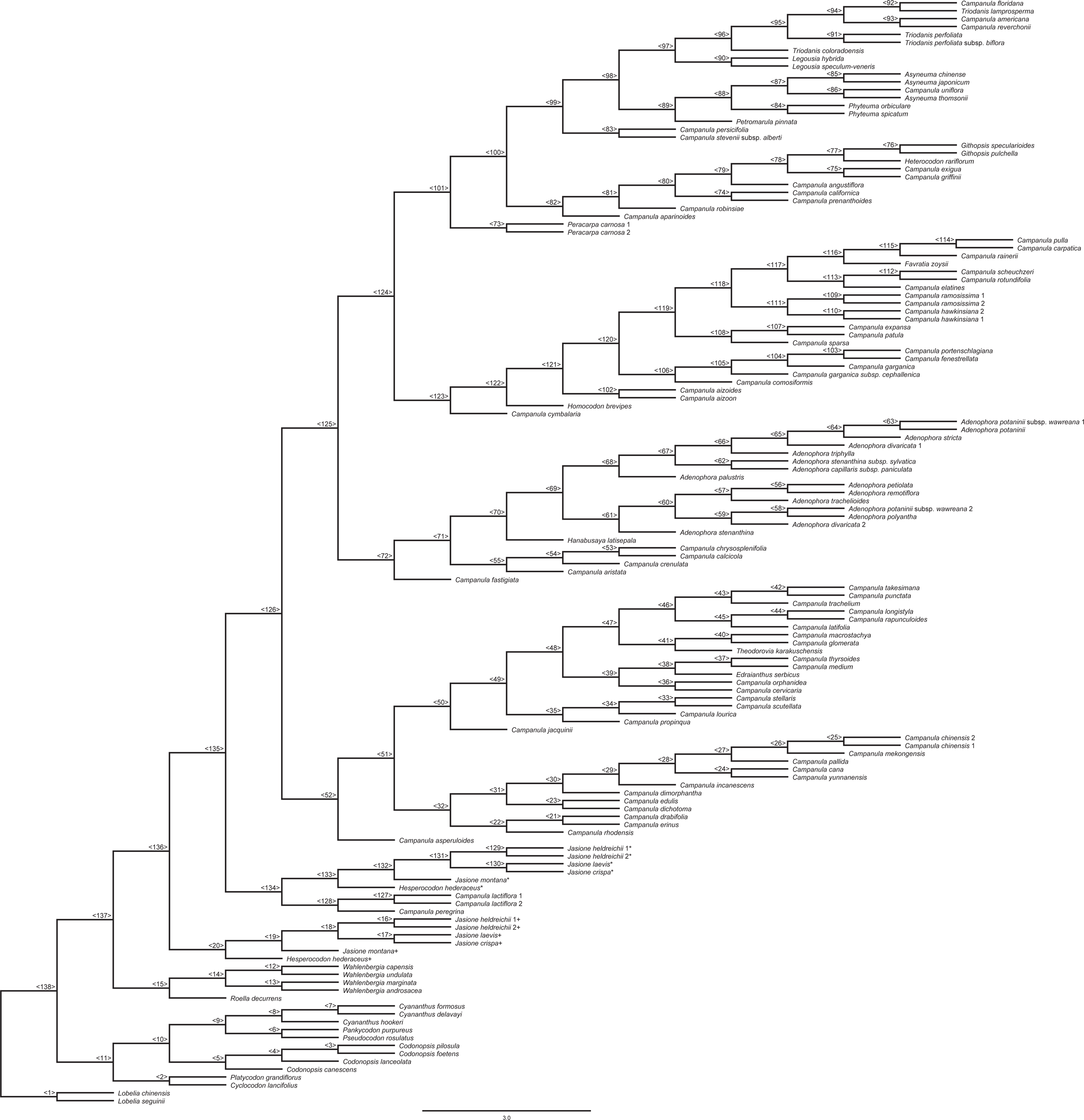

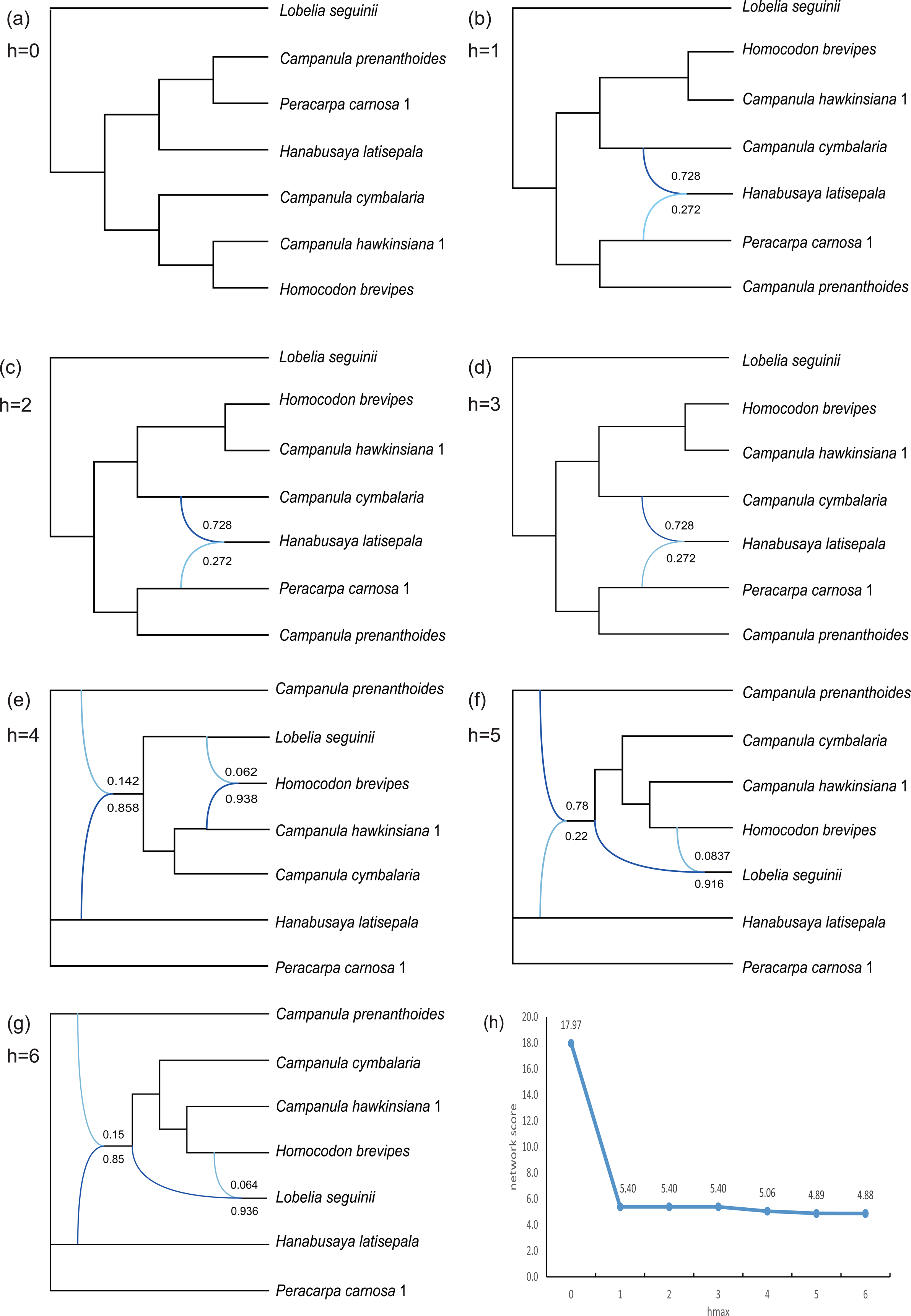

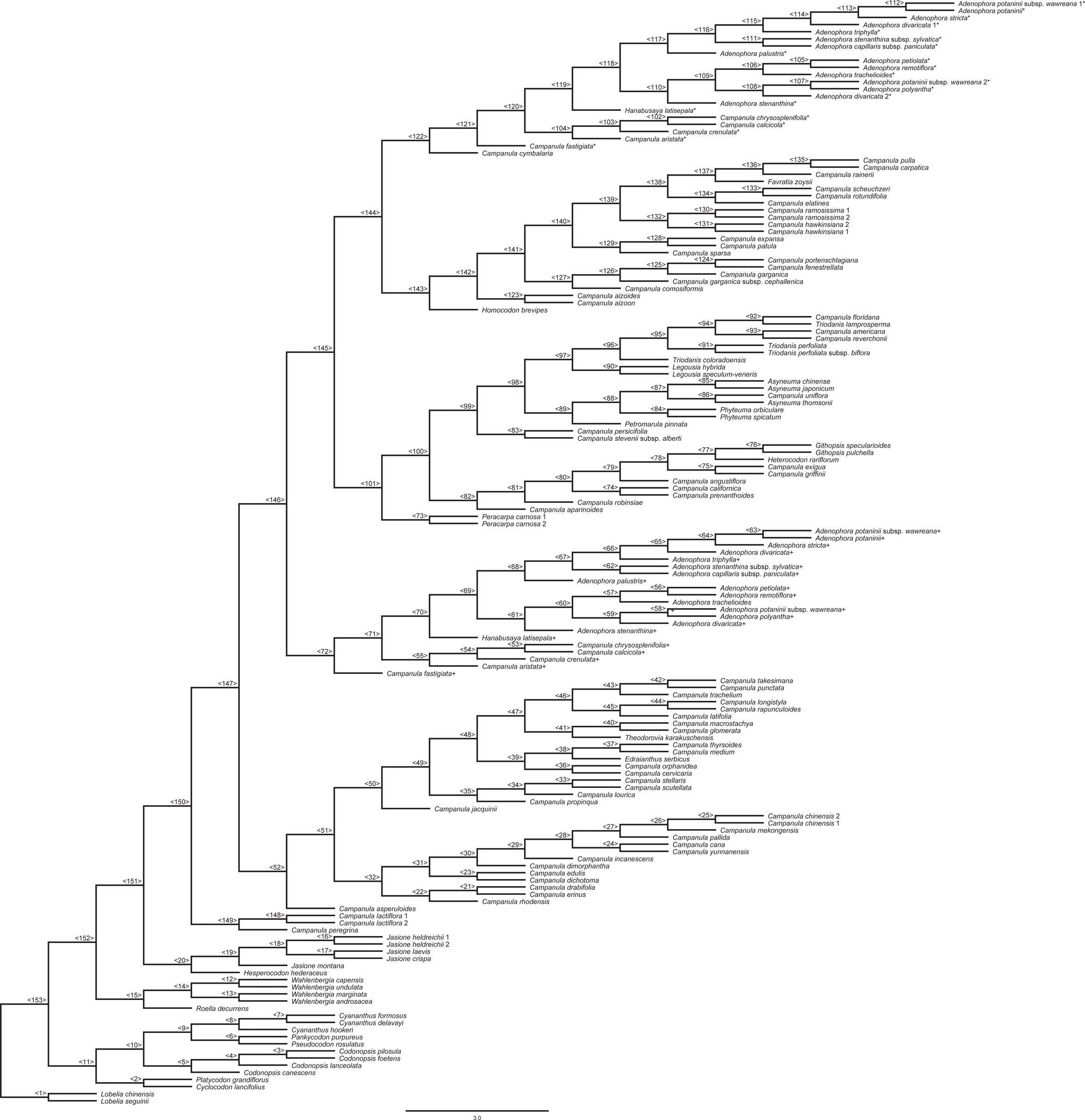

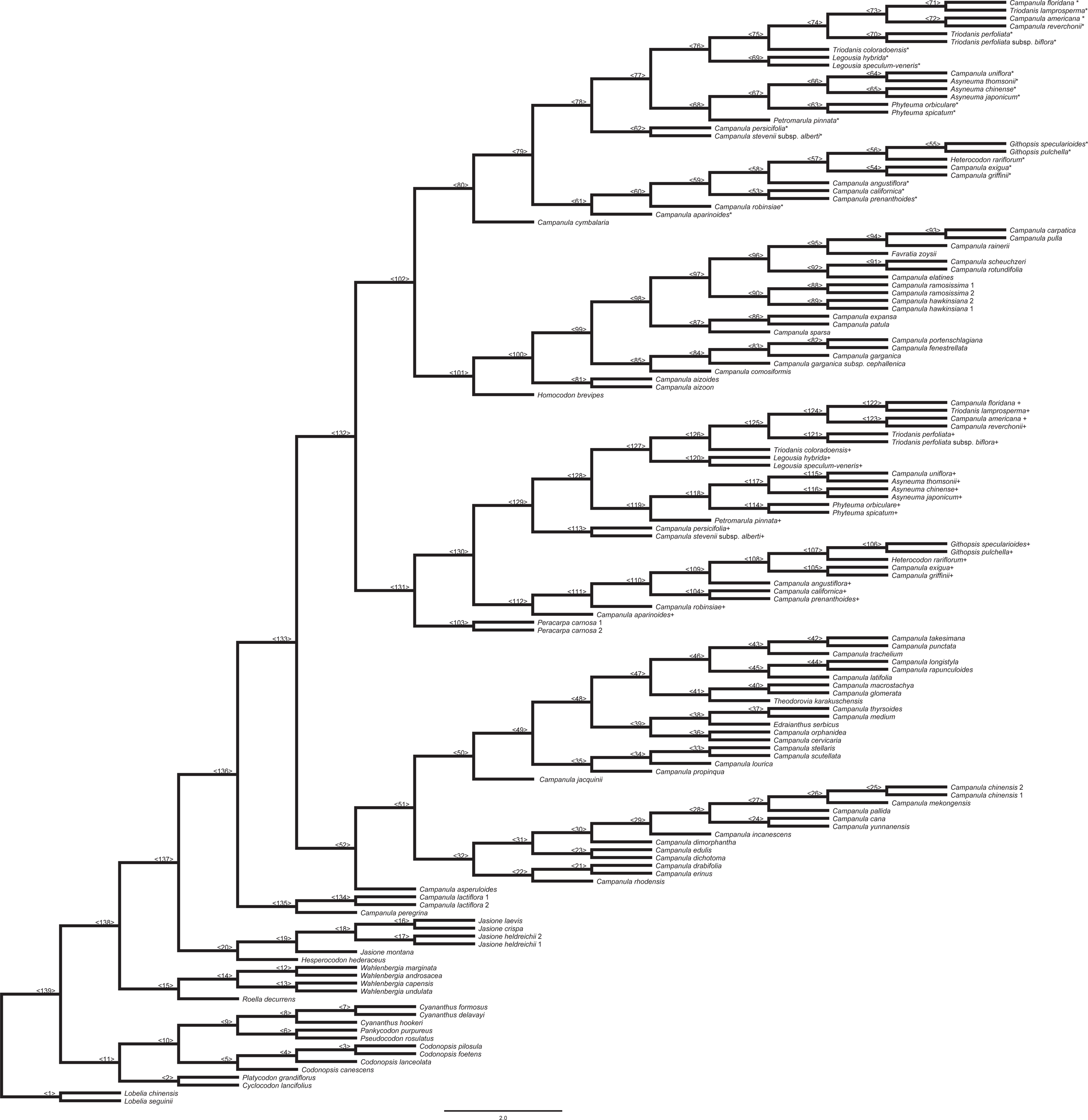

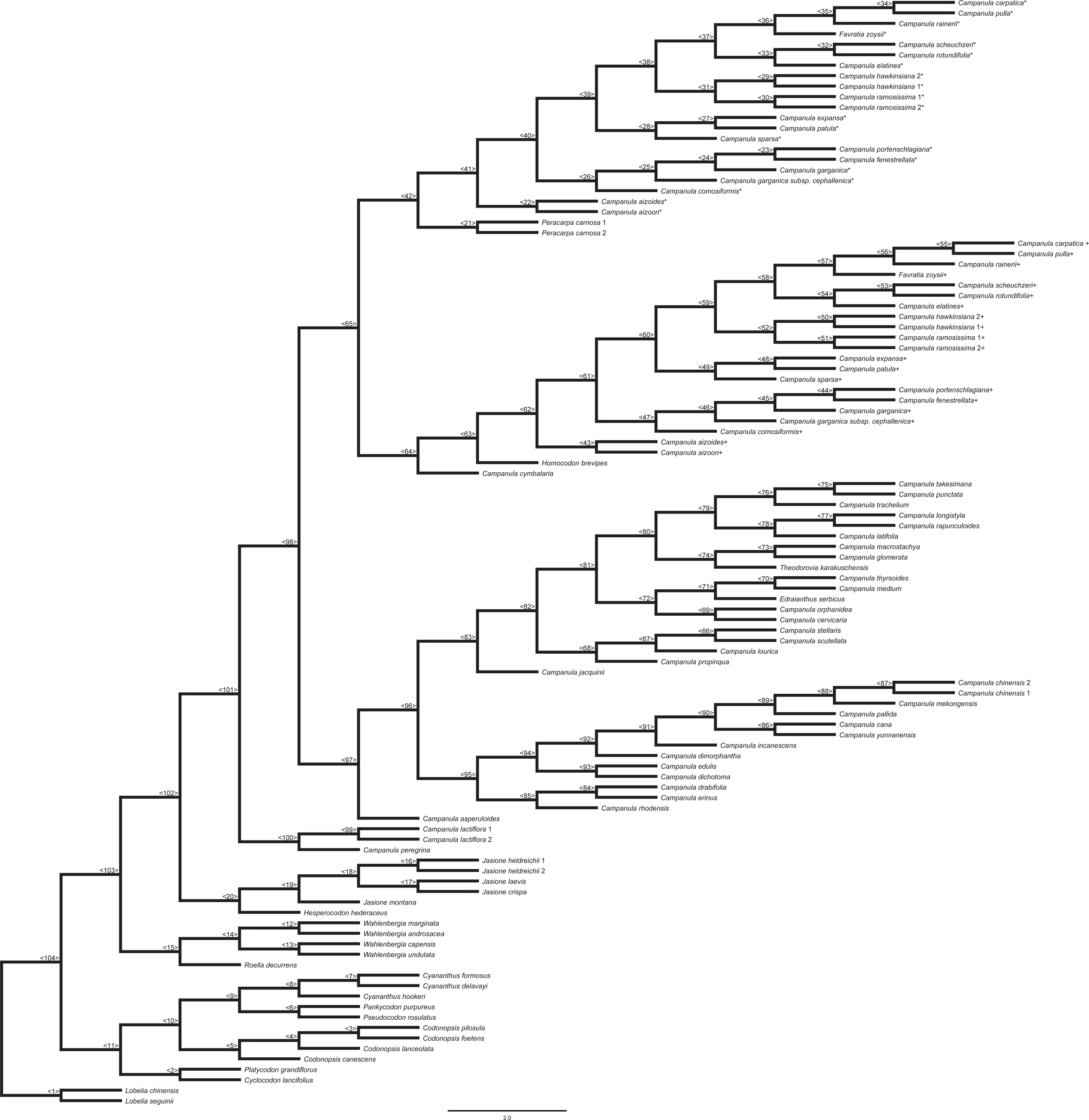

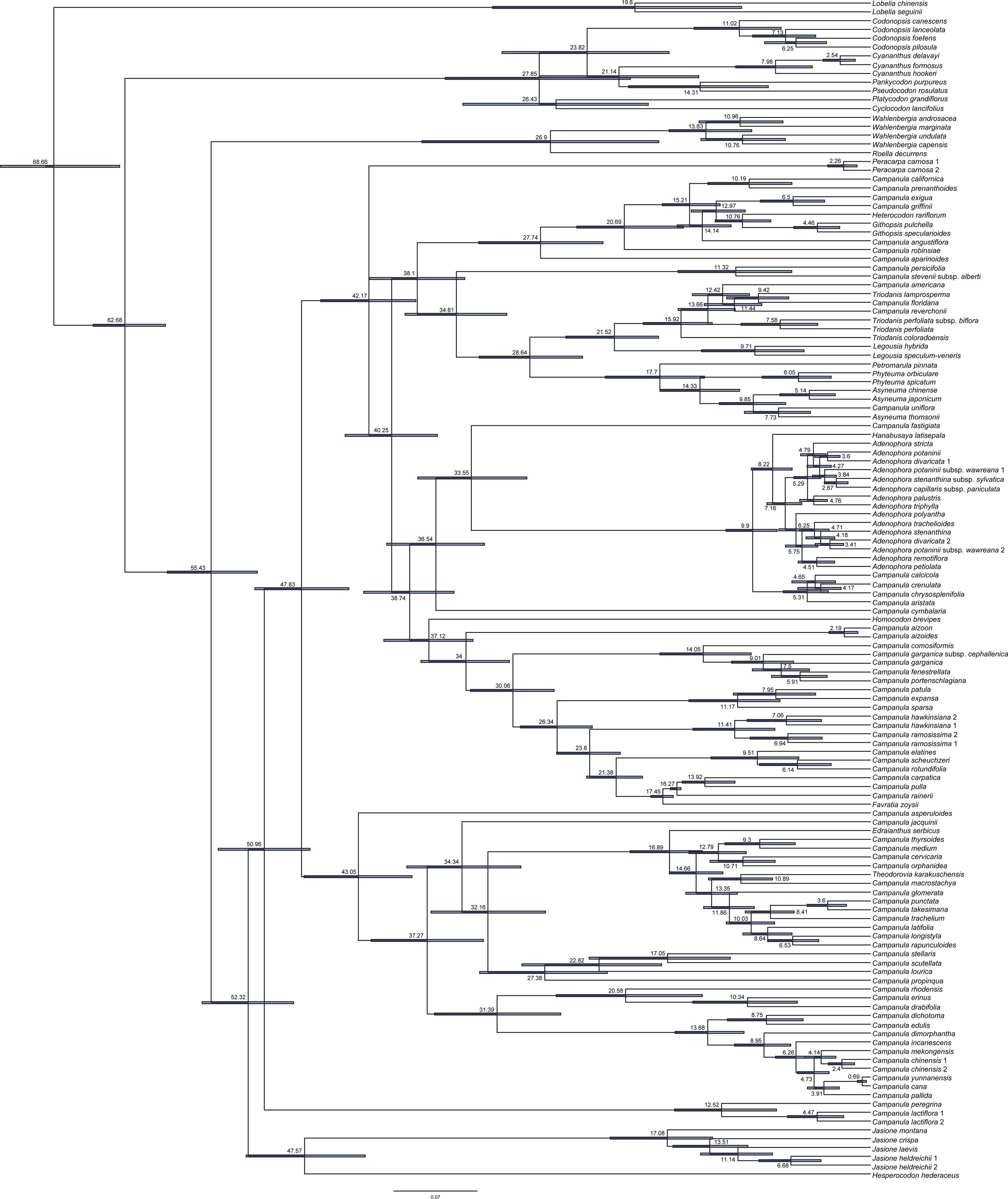

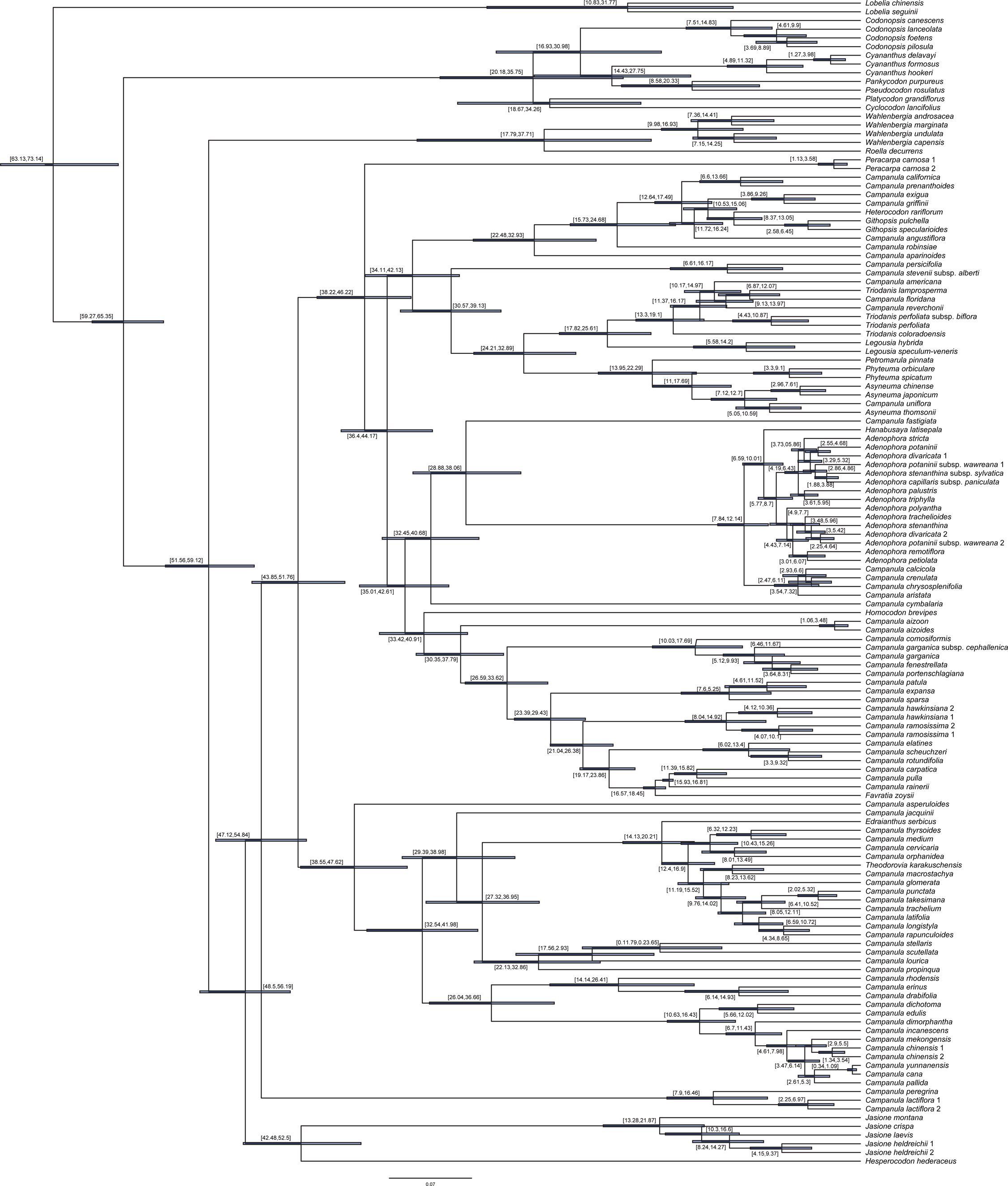

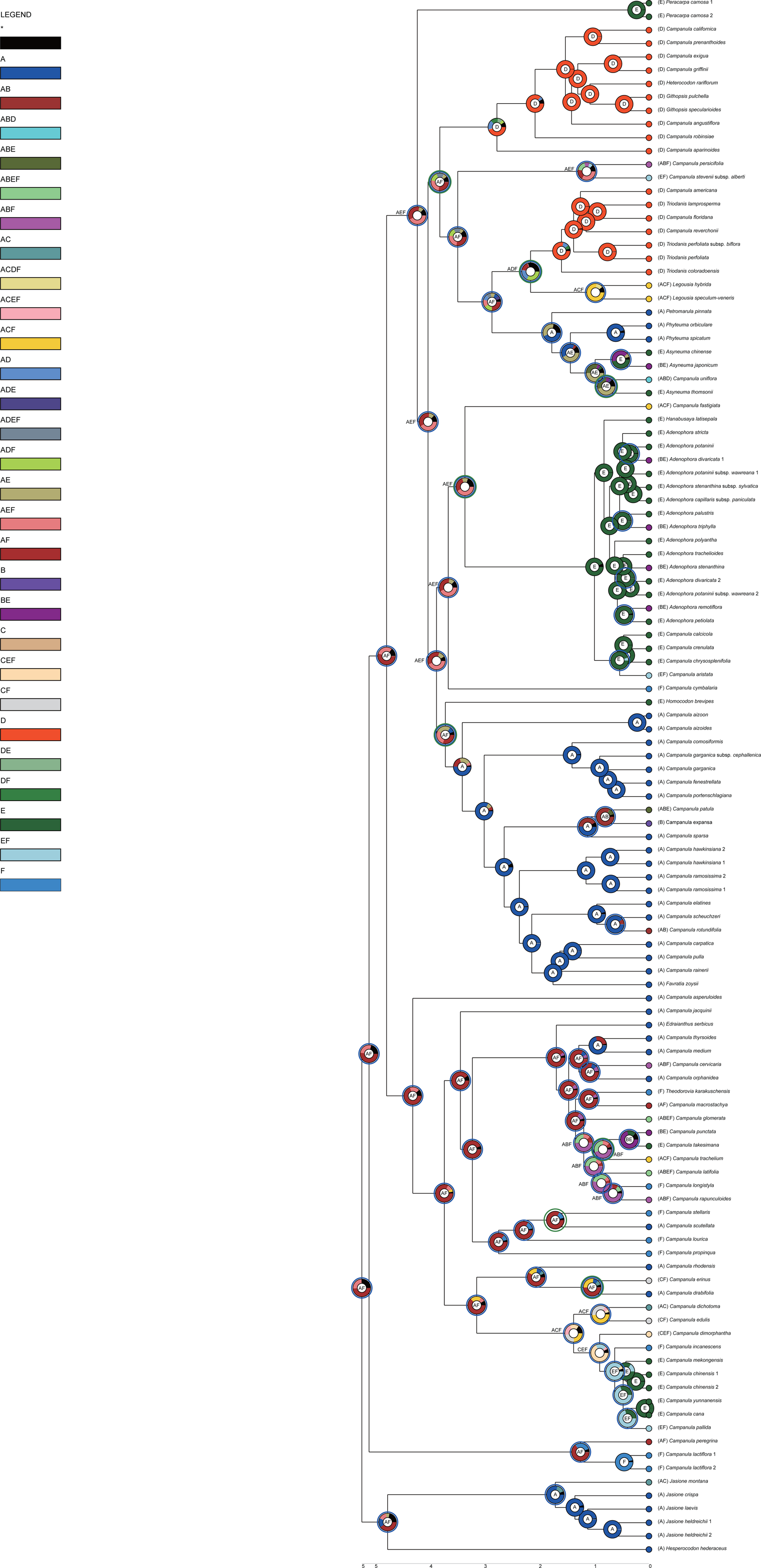

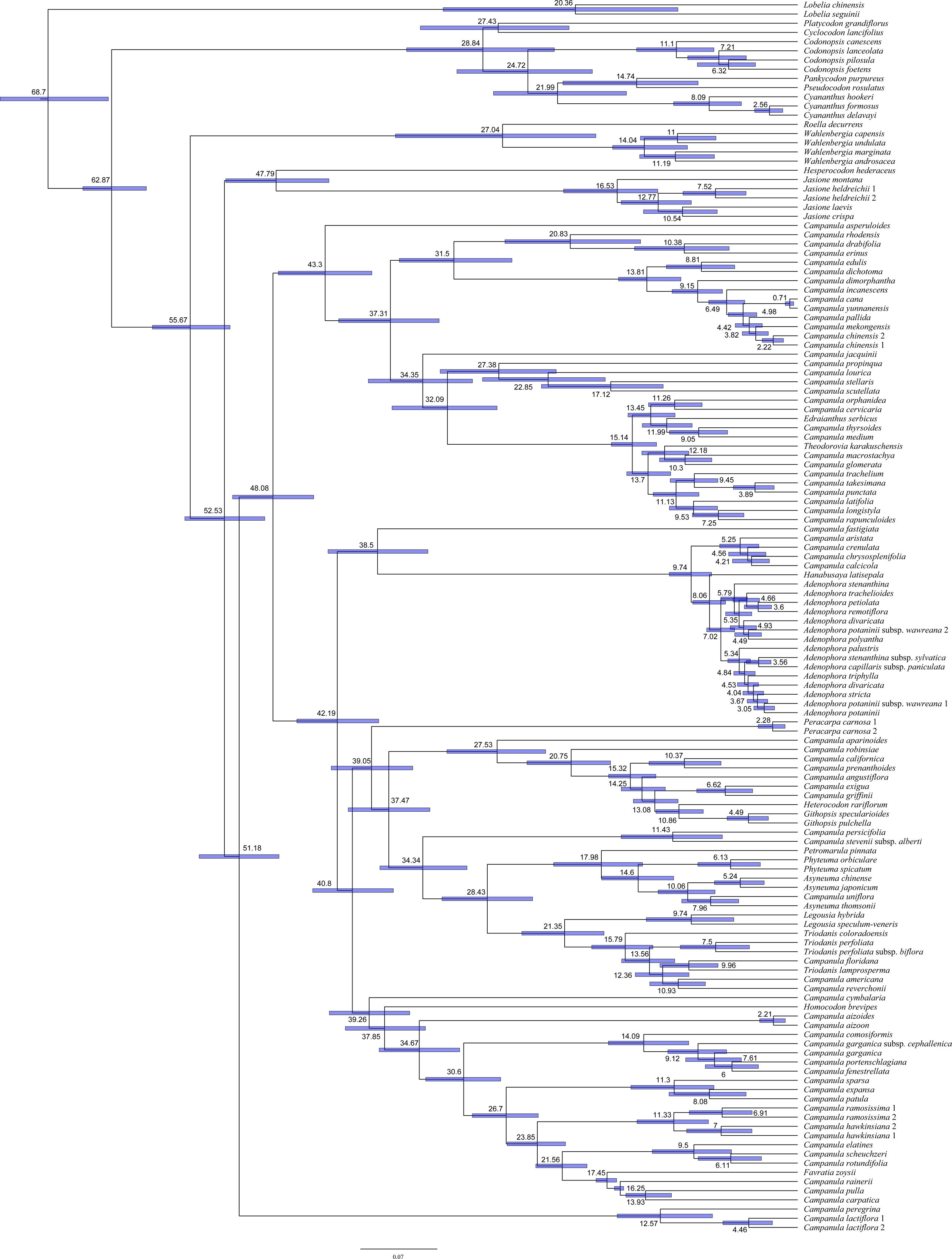

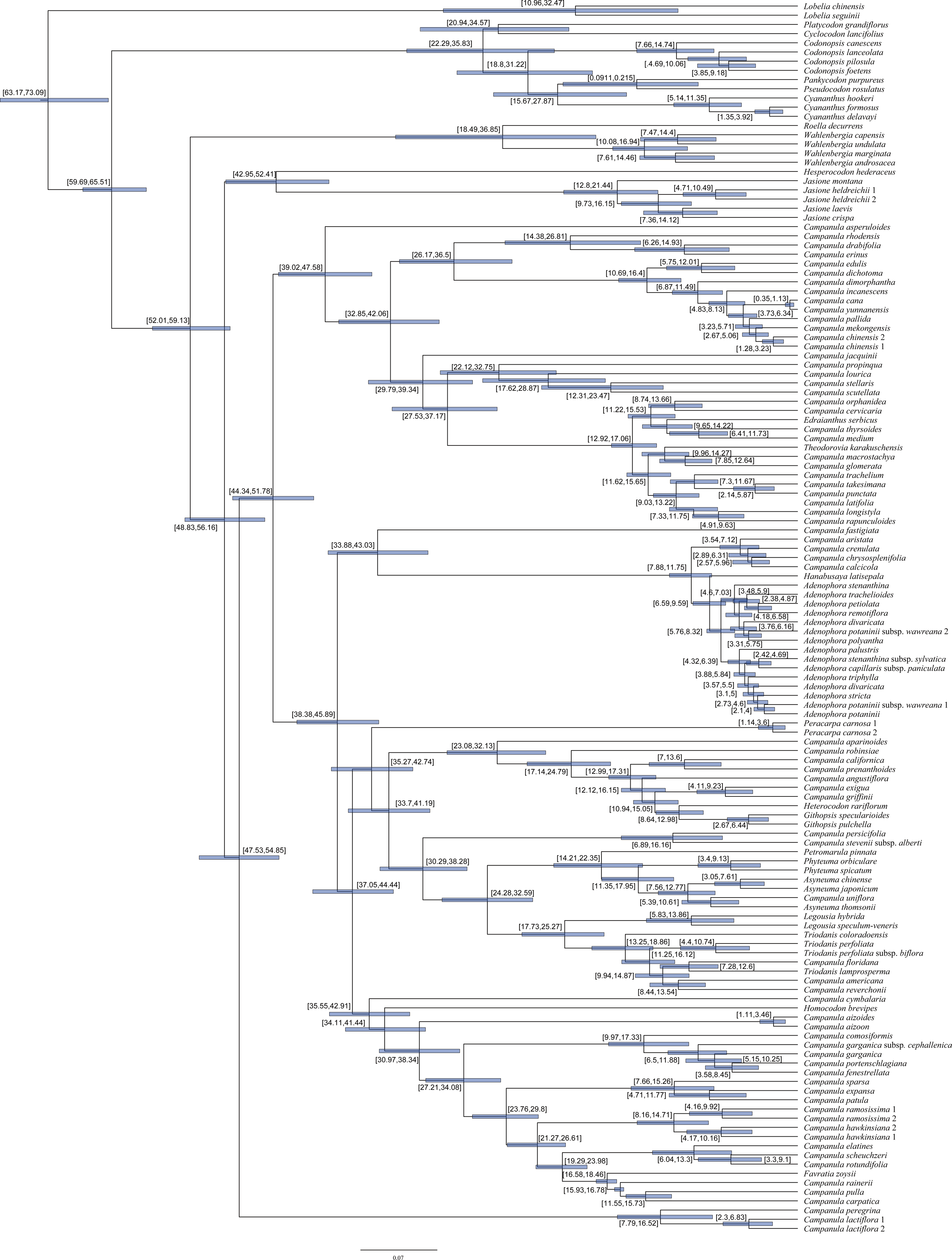

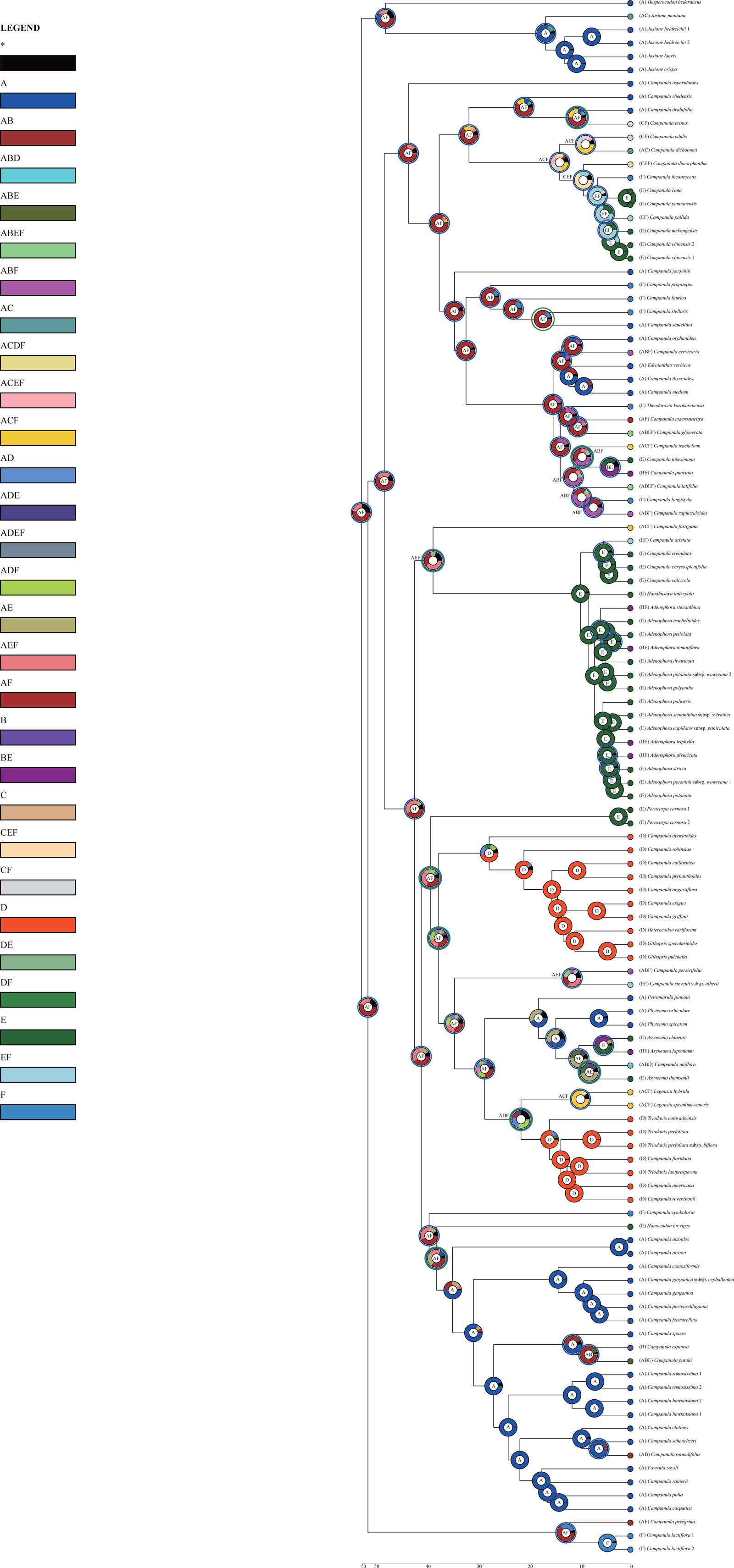

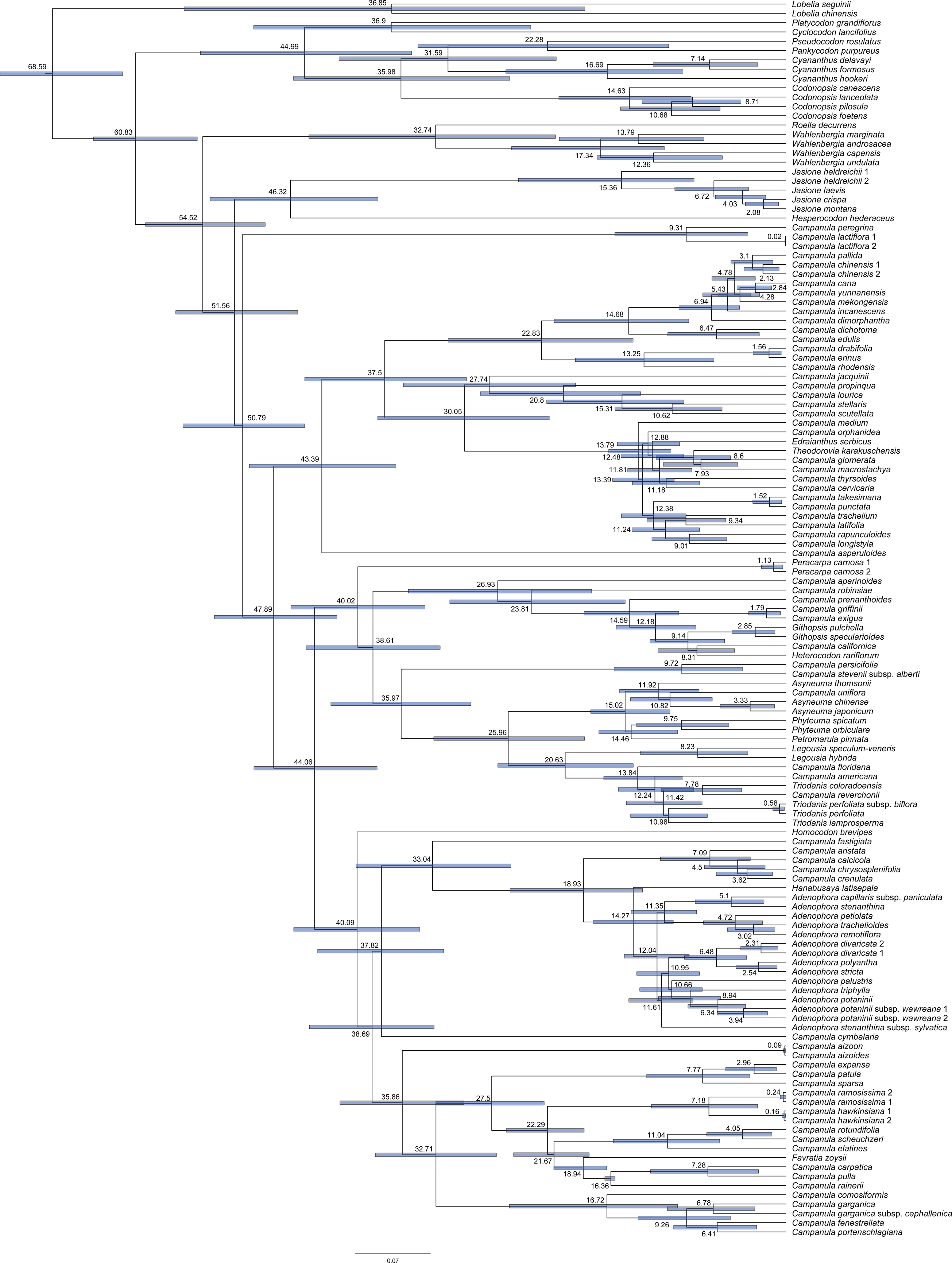

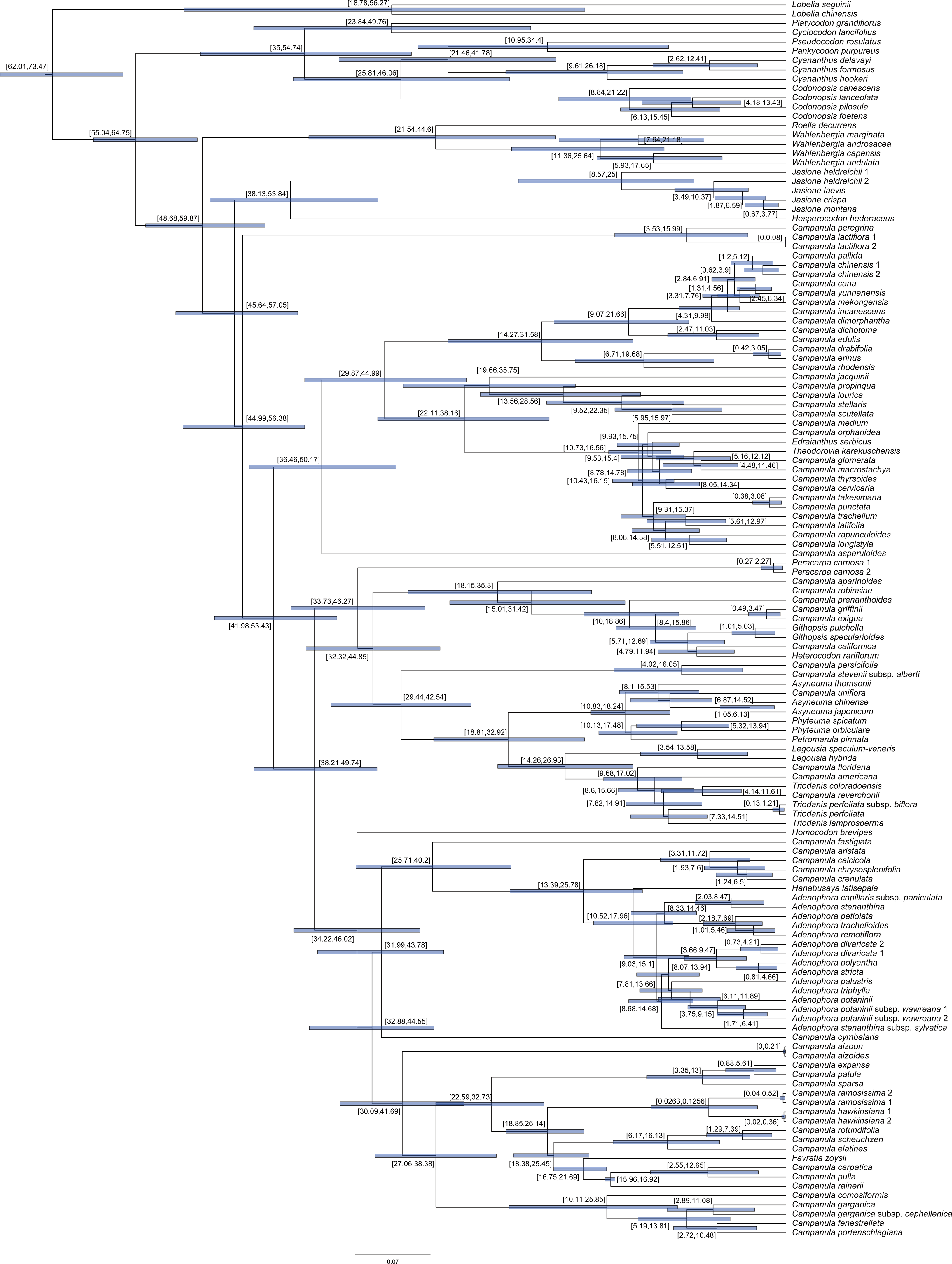

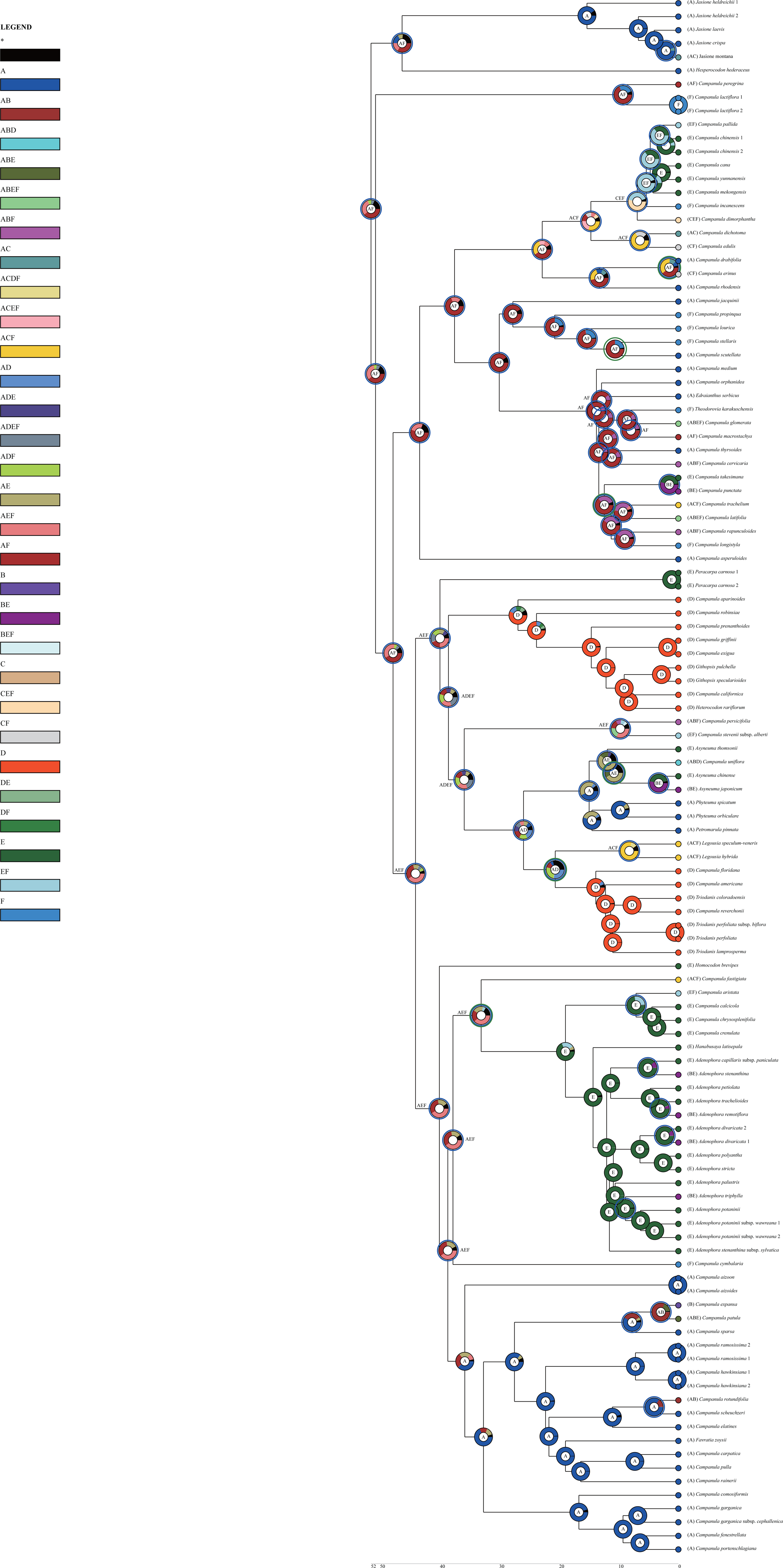

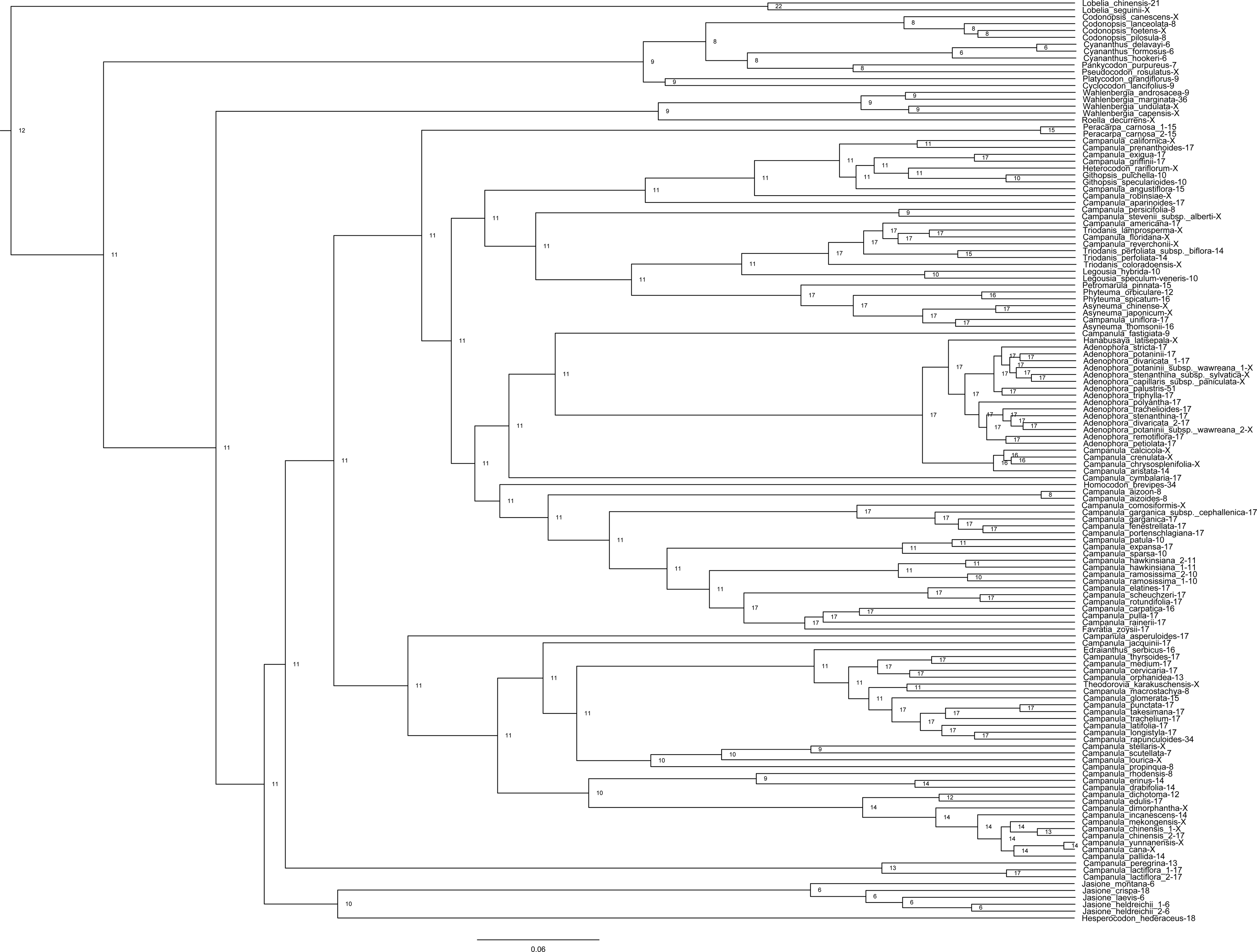

