## Supplementary material for "Dense Sampling of Taxa and Genomes Untangles the Phylogenetic Backbone of a Non-model Plant Lineage Rife with Deep Hybridization and Allopolyploidy": Table S1

| Taxonomic_Hierarchy | subtribe_in_Campanuleae | BAMM_analysis |
|---|---|---|
| Campanuloideae-Cyanantheae | Outgroup | Outgroup |
| Campanuloideae-Campanuleae | Campanulinae | Clade I |
| Campanuloideae-Campanuleae | Campanulinae | Clade H |
| Campanuloideae-Campanuleae | Campanulinae | Clade H |
| Campanuloideae-Campanuleae | Campanulinae | Clade H |
| Campanuloideae-Campanuleae | Campanulinae | Clade H |
| Campanuloideae-Campanuleae | Campanulinae | Clade H |
| Campanuloideae-Campanuleae | Campanulinae | Clade G |
| Campanuloideae-Campanuleae | Campanulinae | Clade G |
| Campanuloideae-Campanuleae | Campanulinae | Clade G |
| Campanuloideae-Campanuleae | Campanulinae | Clade G |
| Campanuloideae-Campanuleae | Campanulinae | Clade H |
| Campanuloideae-Campanuleae | Campanulinae | Clade G |
| Campanuloideae-Campanuleae | Campanulinae | Clade G |
| Campanuloideae-Campanuleae | Campanulinae | Clade H |
| Campanuloideae-Campanuleae | Campanulinae | Clade G |
| Campanuloideae-Campanuleae | Campanulinae | Clade G |
| Campanuloideae-Campanuleae | Campanulinae | Clade G |
| Campanuloideae-Campanuleae | Campanulinae | Clade G |
| Campanuloideae-Campanuleae | Jasioninae | Clade K |
| Campanuloideae-Campanuleae | Jasioninae | Clade K |
| Campanuloideae-Campanuleae | Jasioninae | Clade K |
| Campanuloideae-Campanuleae | Musschiinae | Clade J |
| Campanuloideae-Campanuleae | Musschiinae | Clade J |
| Campanuloideae-Campanuleae | Musschiinae | Clade J |
| Campanuloideae-Campanuleae | Phyteumatinae | Clade F |
| Campanuloideae-Campanuleae | Phyteumatinae | Clade A |
| Campanuloideae-Campanuleae | Phyteumatinae | Clade A |
| Campanuloideae-Campanuleae | Phyteumatinae | Clade C |
| Campanuloideae-Campanuleae | Phyteumatinae | Clade C |
| Campanuloideae-Campanuleae | Phyteumatinae | Clade A |
| Campanuloideae-Campanuleae | Phyteumatinae | Clade A |
| Campanuloideae-Campanuleae | Phyteumatinae | Clade A |
| Campanuloideae-Campanuleae | Phyteumatinae | Clade F |
| Campanuloideae-Campanuleae | Phyteumatinae | Clade F |
| Campanuloideae-Campanuleae | Phyteumatinae | Clade A |
| Campanuloideae-Campanuleae | Phyteumatinae | Clade C |
| Campanuloideae-Campanuleae | Phyteumatinae | Clade F |
| Campanuloideae-Campanuleae | Phyteumatinae | Clade C |
| Campanuloideae-Campanuleae | Phyteumatinae | Clade F |
| Campanuloideae-Campanuleae | Phyteumatinae | Clade E |
| Campanuloideae-Campanuleae | Phyteumatinae | Clade C |
| Campanuloideae-Campanuleae | Phyteumatinae | Clade A |
| Campanuloideae-Campanuleae | Phyteumatinae | Clade C |
| Campanuloideae-Campanuleae | Phyteumatinae | Clade C |
| Campanuloideae-Campanuleae | Phyteumatinae | Clade C |
| Campanuloideae-Campanuleae | Phyteumatinae | Clade C |
| Campanuloideae-Campanuleae | Phyteumatinae | Clade A |
| Campanuloideae-Campanuleae | Phyteumatinae | Clade C |
| Campanuloideae-Campanuleae | Phyteumatinae | Clade C |
| Campanuloideae-Campanuleae | Phyteumatinae | Clade C |

| | | |
|---|---|---|
| Campanuloideae-Campanuleae | Phyteumatinae | Clade A |
| Campanuloideae-Campanuleae | Phyteumatinae | Clade C |
| Campanuloideae-Campanuleae | Phyteumatinae | Clade A |
| Campanuloideae-Campanuleae | Phyteumatinae | Clade C |
| Campanuloideae-Campanuleae | Phyteumatinae | Clade C |
| Campanuloideae-Campanuleae | Phyteumatinae | Clade C |
| Campanuloideae-Campanuleae | Phyteumatinae | Clade C |
| Campanuloideae-Campanuleae | Phyteumatinae | Clade A |
| Campanuloideae-Campanuleae | Phyteumatinae | Clade C |
| Campanuloideae-Campanuleae | Phyteumatinae | Clade C |
| Campanuloideae-Campanuleae | Phyteumatinae | Clade C |
| Campanuloideae-Campanuleae | Phyteumatinae | Clade A |
| Campanuloideae-Campanuleae | Phyteumatinae | Clade C |
| Campanuloideae-Campanuleae | Phyteumatinae | Clade A |
| Campanuloideae-Campanuleae | Phyteumatinae | Clade A |
| Campanuloideae-Campanuleae | Phyteumatinae | Clade F |
| Campanuloideae-Campanuleae | Phyteumatinae | Clade A |
| Campanuloideae-Campanuleae | Phyteumatinae | Clade D |
| Campanuloideae-Campanuleae | Phyteumatinae | Clade A |
| Campanuloideae-Campanuleae | Phyteumatinae | Clade A |
| Campanuloideae-Campanuleae | Phyteumatinae | Clade B |
| Campanuloideae-Campanuleae | Phyteumatinae | Clade B |
| Campanuloideae-Campanuleae | Phyteumatinae | Clade A |
| Campanuloideae-Campanuleae | Phyteumatinae | Clade A |
| Campanuloideae-Campanuleae | Phyteumatinae | Clade A |
| Campanuloideae-Campanuleae | Phyteumatinae | Clade A |
| Campanuloideae-Campanuleae | Phyteumatinae | Clade A |
| Campanuloideae-Cyanantheae | Outgroup | Outgroup |
| Campanuloideae-Wahlenbergieae | Outgroup | Outgroup |
| Campanuloideae-Wahlenbergieae | Outgroup | Outgroup |
| Campanuloideae-Campanuleae | Phyteumatinae | Clade F |
| Campanuloideae-Campanuleae | Phyteumatinae | Clade F |
| Campanuloideae-Campanuleae | Phyteumatinae | Clade F |
| Campanuloideae-Campanuleae | Phyteumatinae | Clade F |
| Campanuloideae-Campanuleae | Phyteumatinae | Clade F |
| Campanuloideae-Campanuleae | Phyteumatinae | Clade F |
| Campanuloideae-Campanuleae | Phyteumatinae | Clade F |
| Campanuloideae-Campanuleae | Phyteumatinae | Clade F |
| Campanuloideae-Campanuleae | Phyteumatinae | Clade F |
| Campanuloideae-Campanuleae | Phyteumatinae | Clade F |
| Campanuloideae-Campanuleae | Campanulinae | Clade H |
| Campanuloideae-Campanuleae | Campanulinae | Clade G |
| Campanuloideae-Campanuleae | Campanulinae | Clade H |
| Campanuloideae-Campanuleae | Campanulinae | Clade H |
| Campanuloideae-Campanuleae | Campanulinae | Clade G |
| Campanuloideae-Campanuleae | Campanulinae | Clade G |
| Campanuloideae-Campanuleae | Campanulinae | Clade G |
| Campanuloideae-Campanuleae | Campanulinae | Clade H |
| Campanuloideae-Campanuleae | Campanulinae | Clade G |
| Campanuloideae-Campanuleae | Campanulinae | Clade G |
| Campanuloideae-Campanuleae | Campanulinae | Clade G |
| Campanuloideae-Campanuleae | Campanulinae | Clade G |

| | | |
|---|---|---|
| Campanuloideae-Campanuleae | Campanulinae | Clade G |
| Campanuloideae-Campanuleae | Campanulinae | Clade H |
| Campanuloideae-Campanuleae | Jasioninae | Clade K |
| Campanuloideae-Campanuleae | Jasioninae | Clade K |
| Campanuloideae-Campanuleae | Jasioninae | Clade K |
| Campanuloideae-Campanuleae | Phyteumatinae | Clade A |
| Campanuloideae-Campanuleae | Phyteumatinae | Clade F |
| Campanuloideae-Campanuleae | Phyteumatinae | Clade A |
| Campanuloideae-Campanuleae | Phyteumatinae | Clade A |
| Campanuloideae-Campanuleae | Phyteumatinae | Clade A |
| Campanuloideae-Campanuleae | Phyteumatinae | Clade A |
| Campanuloideae-Campanuleae | Phyteumatinae | Clade A |
| Campanuloideae-Cyanantheae | Outgroup | Outgroup |
| Campanuloideae-Cyanantheae | Outgroup | Outgroup |
| Campanuloideae-Cyanantheae | Outgroup | Outgroup |
| Campanuloideae-Cyanantheae | Outgroup | Outgroup |
| Campanuloideae-Cyanantheae | Outgroup | Outgroup |
| Campanuloideae-Cyanantheae | Outgroup | Outgroup |
| Campanuloideae-Cyanantheae | Outgroup | Outgroup |
| Campanuloideae-Cyanantheae | Outgroup | Outgroup |
| Campanuloideae-Cyanantheae | Outgroup | Outgroup |
| Campanuloideae-Wahlenbergieae | Outgroup | Outgroup |
| Campanuloideae-Wahlenbergieae | Outgroup | Outgroup |
| Campanuloideae-Wahlenbergieae | Outgroup | Outgroup |
| Lobelioideae | Outgroup | Outgroup |
| Lobelioideae | Outgroup | Outgroup |
| Campanuloideae-Campanuleae | Campanulinae | Clade H |
| Campanuloideae-Campanuleae | Phyteumatinae | Clade F |
| Campanuloideae-Campanuleae | Phyteumatinae | Clade F |
| Campanuloideae-Campanuleae | Phyteumatinae | Clade F |
| Campanuloideae-Campanuleae | Phyteumatinae | Clade F |
| Campanuloideae-Campanuleae | Phyteumatinae | Clade F |
| Campanuloideae-Campanuleae | Phyteumatinae | Clade F |
| Campanuloideae-Campanuleae | Phyteumatinae | Clade F |
| Campanuloideae-Campanuleae | Phyteumatinae | Clade F |

| Clades_sensu Xu et al. (2021) | species name | Number of reads |
|---|---|---|
| Outgroup | *Codonopsis pilosula* | 746553662 |
| Clade cam13 | *Campanula asperuloides* | 8868476 |
| Clade cam15 | *Campanula chinensis 1* | 9396344 |
| Clade cam15 | *Campanula dimorphantha* | 8679256 |
| Clade cam14 | *Campanula drabifolia* | 6604762 |
| Clade cam15 | *Campanula edulis* | 5375438 |
| Clade cam14 | *Campanula erinus* | 16135058 |
| Clade cam17 | *Campanula glomerata* | 9813304 |
| Clade cam16 | *Campanula jacquinii* | 7634783 |
| Clade cam17 | *Campanula latifolia* | 5305022 |
| Clade cam16 | *Campanula lourica* | 6533257 |
| Clade cam15 | *Campanula pallida* | 17871338 |
| Clade cam16 | *Campanula propinqua* | 6177320 |
| Clade cam17 | *Campanula punctata* | 12740716 |
| Clade cam14 | *Campanula rhodensis* | 5951508 |
| Clade cam16 | *Campanula scutellata* | 6925265 |
| Clade cam16 | *Campanula stellaris* | 6116894 |
| Clade cam17 | *Edraianthus serbicus* | 6196839 |
| Clade cam17 | *Theodorovia karakuschensis* | 7613878 |
| Clade Jasione | *Jasione crispa* | 8532488 |
| Clade Jasione | *Jasione heldreichii 2* | 9304313 |
| Clade Jasione | *Jasione montana* | 11055573 |
| Clade cam01 | *Campanula lactiflora* 1 | 8982199 |
| Clade cam01 | *Campanula lactiflora* 2 | 10951978 |
| Clade cam01 | *Campanula peregrina* | 11098072 |
| Clade cam06 | *Adenophora polyantha* | 8835041 |
| Clade cam04 | *Asyneuma chinense* | 8055597 |
| Clade cam04 | *Asyneuma japonicum* | 5654433 |
| Clade cam07 | *Campanula aizoides* | 8092251 |
| Clade cam07 | *Campanula aizoon* | 5071216 |
| Clade cam04 | *Campanula americana* | 9199944 |
| Clade cam02 | *Campanula angustiflora* | 7140767 |
| Clade cam02 | *Campanula aparinoides* | 29393669 |
| Clade cam06 | *Campanula aristata* | 4875136 |
| Clade cam06 | *Campanula calcicola* | 6578432 |
| Clade cam02 | *Campanula californica* | 7063504 |
| Clade cam11 | *Campanula carpatica* | 4628872 |
| Clade cam06 | *Campanula chrysosplenifolia* | 6216964 |
| Clade cam08 | *Campanula comosiformis* | 11969460 |
| Clade cam06 | *Campanula crenulata* | 8112949 |
| Clade cam05 | *Campanula cymbalaria* | 8908548 |
| Clade cam08 | *Campanula elatines* | 9533631 |
| Clade cam02 | *Campanula exigua* | 5724737 |
| Clade cam09 | *Campanula expansa* | 3165420 |
| Clade cam08 | *Campanula fenestrellata* | 9082837 |
| Clade cam08 | *Campanula garganica* | 9341262 |
| Clade cam08 | *Campanula garganica* subsp. *cephallenica* | 11611956 |
| Clade cam02 | *Campanula griffinii* | 4010736 |
| Clade cam10 | *Campanula hawkinsiana* 1 | 4505400 |
| Clade cam10 | *Campanula hawkinsiana* 2 | 9397394 |
| Clade cam09 | *Campanula patula* | 11933371 |

| | | |
|---|---|---|
| Clade cam03 | *Campanula persicifolia* | 9053555 |
| Clade cam08 | *Campanula portenschlagiana* | 9054588 |
| Clade cam02 | *Campanula prenanthoides* | 5542547 |
| Clade cam11 | *Campanula pulla* | 1893527 |
| Clade cam11 | *Campanula rainerii* | 7709480 |
| Clade cam10 | *Campanula ramosissima* 1 | 5400324 |
| Clade cam10 | *Campanula ramosissima* 2 | 8531017 |
| Clade cam02 | *Campanula robinsiae* | 11738447 |
| Clade cam12 | *Campanula rotundifolia* | 8842035 |
| Clade cam12 | *Campanula scheuchzeri* | 14583158 |
| Clade cam09 | *Campanula sparsa* | 11650292 |
| Clade cam03 | *Campanula stevenii* subsp. *alberti* | 8719369 |
| Clade Favratia | *Favratia zoysii* | 6301321 |
| Clade cam02 | *Githopsis pulchella* | 5241340 |
| Clade cam02 | *Githopsis specularioides* | 7361539 |
| Clade cam06 | *Hanabusaya latisepala* | 3501012 |
| Clade cam02 | *Heterocodon rariflorum* | 14285928 |
| Clade Homocodon | *Homocodon brevipes* | 12879326 |
| Clade cam04 | *Legousia hybrida* | 8310850 |
| Clade cam04 | *Legousia speculum-veneris* | 7613780 |
| Clade Peracarpa | *Peracarpa carnosa* 1 | 4499048 |
| Clade Peracarpa | *Peracarpa carnosa* 2 | 6498238 |
| Clade cam04 | *Petromarula pinnata* | 7004899 |
| Clade cam04 | *Phyteuma orbiculare* | 11099316 |
| Clade cam04 | *Phyteuma spicatum* | 8969986 |
| Clade cam04 | *Triodanis perfoliata* | 7920941 |
| Clade cam04 | *Triodanis perfoliata* subsp. *biflora* | 8540104 |
| Outgroup | *Platycodon grandiflorus* | 9491206 |
| Outgroup | *Roella decurrens* | 11638166 |
| Outgroup | *Wahlenbergia undulata* | 9628679 |
| Clade cam06 | *Adenophora petiolata* | 6049351 |
| Clade cam06 | *Adenophora remotiflora* | 11163214 |
| Clade cam06 | *Adenophora palustris* | 22697004 |
| Clade cam06 | *Adenophora triphylla* | 25421211 |
| Clade cam06 | *Adenophora capillaris* subsp. *paniculata* | 22322099 |
| Clade cam06 | *Adenophora divaricata* 1 | 27168137 |
| Clade cam06 | *Adenophora potaninii* | 23613271 |
| Clade cam06 | *Adenophora potaninii* subsp. *wawreana* 1 | 23139846 |
| Clade cam06 | *Adenophora stenanthina* subsp. *sylvatica* | 20957838 |
| Clade cam06 | *Adenophora stricta* | 22273508 |
| Clade cam15 | *Campanula cana* | 423041398 |
| Clade cam17 | *Campanula cervicaria* | 376031430 |
| Clade cam15 | *Campanula dichotoma* | 405944796 |
| Clade cam15 | *Campanula incanescens* | 306273288 |
| Clade cam17 | *Campanula longistyla* | 454081018 |
| Clade cam17 | *Campanula macrostachya* | 414231314 |
| Clade cam17 | *Campanula medium* | 277062170 |
| Clade cam15 | *Campanula mekongensis* | 405341946 |
| Clade cam17 | *Campanula orphanidea* | 278468732 |
| Clade cam17 | *Campanula rapunculoides* | 251994178 |
| Clade cam17 | *Campanula takesimana* | 215874374 |
| Clade cam17 | *Campanula thyrsoides* | 295015068 |

| | | |
|---|---|---|
| Clade cam17 | *Campanula trachelium* | 413102014 |
| Clade cam15 | *Campanula yunnanensis* | 345067168 |
| Clade Jasione | *Hesperocodon hederaceus* | 361754336 |
| Clade Jasione | *Jasione heldreichii 1* | 261426928 |
| Clade Jasione | *Jasione laevis* | 128428212 |
| Clade cam04 | *Asyneuma thomsonii* | 396002522 |
| Clade cam05 | *Campanula fastigiata* | 352427784 |
| Clade cam04 | *Campanula floridana* | 352165524 |
| Clade cam04 | *Campanula reverchonii* | 419076940 |
| Clade cam04 | *Campanula uniflora* | 294882588 |
| Clade cam04 | *Triodanis coloradoensis* | 85402602 |
| Clade cam04 | *Triodanis lamprosperma* | 288469206 |
| Outgroup | *Codonopsis canescens* | 372746540 |
| Outgroup | *Codonopsis foetens* | 347498090 |
| Outgroup | *Codonopsis lanceolata* | 369436871 |
| Outgroup | *Cyananthus delavayi* | 460015756 |
| Outgroup | *Cyananthus formosus* | 392466318 |
| Outgroup | *Cyananthus hookeri* | 300705250 |
| Outgroup | *Cyclocodon lancifolius* | 274095220 |
| Outgroup | *Pankycodon purpureus* | 235388248 |
| Outgroup | *Pseudocodon rosulatus* | 358904876 |
| Outgroup | *Wahlenbergia androsacea* | 377767992 |
| Outgroup | *Wahlenbergia capensis* | 396747958 |
| Outgroup | *Wahlenbergia marginata* | 441838312 |
| Outgroup | *Lobelia chinensis* | 236138193 |
| Outgroup | *Lobelia seguinii* | 350867002 |
| Clade cam15 | *Campanula chinensis 2* | 343721893 |
| Clade cam06 | *Adenophora divaricata* 2 | 365129660 |
| Clade cam06 | *Adenophora potaninii subsp. wawreana* 2 | 359745725 |
| Clade cam06 | *Adenophora stenanthina* | 374956604 |
| Clade cam06 | Adenophora trachelioides | 402903678 |
| Clade cam06 | *Adenophora palustris* | 29438236 |
| Clade cam06 | *Adenophora petiolata* | 23493943 |
| Clade cam06 | *Adenophora remotiflora* | 18135047 |
| Clade cam06 | *Adenophora triphylla* | 35348691 |

| Number of bases (bp) | data size (G) | Library_Strategy | Platform | Model |
|---|---|---|---|---|
| 111983049300 | 104.29 | Hi-C | BGISEQ | BGISEQ-500 |
| 1330271400 | 1.24 | Hyb-Seq† | ILLUMINA | Illumina NovaSeq 6000 |
| 1409451600 | 1.31 | Hyb-Seq† | ILLUMINA | Illumina NovaSeq 6000 |
| 1301888400 | 1.21 | Hyb-Seq† | ILLUMINA | Illumina NovaSeq 6000 |
| 990714300 | 0.92 | Hyb-Seq† | ILLUMINA | Illumina NovaSeq 6000 |
| 806315700 | 0.75 | Hyb-Seq† | ILLUMINA | Illumina NovaSeq 6000 |
| 2420258700 | 2.25 | Hyb-Seq† | ILLUMINA | Illumina NovaSeq 6000 |
| 1471995600 | 1.37 | Hyb-Seq† | ILLUMINA | Illumina NovaSeq 6000 |
| 1145217450 | 1.07 | Hyb-Seq† | ILLUMINA | Illumina NovaSeq 6000 |
| 795753300 | 0.74 | Hyb-Seq† | ILLUMINA | Illumina NovaSeq 6000 |
| 979988550 | 0.91 | Hyb-Seq† | ILLUMINA | Illumina NovaSeq 6000 |
| 2680700700 | 2.50 | Hyb-Seq† | ILLUMINA | Illumina NovaSeq 6000 |
| 926598000 | 0.86 | Hyb-Seq† | ILLUMINA | Illumina NovaSeq 6000 |
| 1911107400 | 1.78 | Hyb-Seq† | ILLUMINA | Illumina NovaSeq 6000 |
| 892726200 | 0.83 | Hyb-Seq† | ILLUMINA | Illumina NovaSeq 6000 |
| 1038789750 | 0.97 | Hyb-Seq† | ILLUMINA | Illumina NovaSeq 6000 |
| 917534100 | 0.85 | Hyb-Seq† | ILLUMINA | Illumina NovaSeq 6000 |
| 929525850 | 0.87 | Hyb-Seq† | ILLUMINA | Illumina NovaSeq 6000 |
| 1142081700 | 1.06 | Hyb-Seq† | ILLUMINA | Illumina NovaSeq 6000 |
| 1279873200 | 1.19 | Hyb-Seq† | ILLUMINA | Illumina NovaSeq 6000 |
| 1395646950 | 1.30 | Hyb-Seq† | ILLUMINA | |
| 1658335950 | 1.54 | Hyb-Seq† | ILLUMINA | Illumina NovaSeq 6000 |
| 1347329850 | 1.25 | Hyb-Seq† | ILLUMINA | Illumina NovaSeq 6000 |
| 1642796700 | 1.53 | Hyb-Seq† | ILLUMINA | Illumina NovaSeq 6000 |
| 1664710800 | 1.55 | Hyb-Seq† | ILLUMINA | Illumina NovaSeq 6000 |
| 1325256150 | 1.23 | Hyb-Seq† | ILLUMINA | Illumina NovaSeq 6000 |
| 1208339550 | 1.13 | Hyb-Seq† | ILLUMINA | Illumina NovaSeq 6000 |
| 848164950 | 0.79 | Hyb-Seq† | ILLUMINA | Illumina NovaSeq 6000 |
| 1213837650 | 1.13 | Hyb-Seq† | ILLUMINA | Illumina NovaSeq 6000 |
| 760682400 | 0.71 | Hyb-Seq† | ILLUMINA | Illumina NovaSeq 6000 |
| 1379991600 | 1.29 | Hyb-Seq† | ILLUMINA | Illumina NovaSeq 6000 |
| 1071115050 | 1.00 | Hyb-Seq† | ILLUMINA | Illumina NovaSeq 6000 |
| 4409050350 | 4.11 | Hyb-Seq† | ILLUMINA | Illumina NovaSeq 6000 |
| 731270400 | 0.68 | Hyb-Seq† | ILLUMINA | Illumina NovaSeq 6000 |
| 986764800 | 0.92 | Hyb-Seq† | ILLUMINA | Illumina NovaSeq 6000 |
| 1059525600 | 0.99 | Hyb-Seq† | ILLUMINA | Illumina NovaSeq 6000 |
| 694330800 | 0.65 | Hyb-Seq† | ILLUMINA | Illumina NovaSeq 6000 |
| 932544600 | 0.87 | Hyb-Seq† | ILLUMINA | Illumina NovaSeq 6000 |
| 1795419000 | 1.67 | Hyb-Seq† | ILLUMINA | Illumina NovaSeq 6000 |
| 1216942350 | 1.13 | Hyb-Seq† | ILLUMINA | Illumina NovaSeq 6000 |
| 1336282200 | 1.24 | Hyb-Seq† | ILLUMINA | Illumina NovaSeq 6000 |
| 1430044650 | 1.33 | Hyb-Seq† | ILLUMINA | Illumina NovaSeq 6000 |
| 858710550 | 0.80 | Hyb-Seq† | ILLUMINA | Illumina NovaSeq 6000 |
| 474813000 | 0.44 | Hyb-Seq† | ILLUMINA | Illumina NovaSeq 6000 |
| 1362425550 | 1.27 | Hyb-Seq† | ILLUMINA | Illumina NovaSeq 6000 |
| 1401189300 | 1.30 | Hyb-Seq† | ILLUMINA | Illumina NovaSeq 6000 |
| 1741793400 | 1.62 | Hyb-Seq† | ILLUMINA | Illumina NovaSeq 6000 |
| 601610400 | 0.56 | Hyb-Seq† | ILLUMINA | Illumina NovaSeq 6000 |
| 675810000 | 0.63 | Hyb-Seq† | ILLUMINA | Illumina NovaSeq 6000 |
| 1409609100 | 1.31 | Hyb-Seq† | ILLUMINA | Illumina NovaSeq 6000 |
| 1790005650 | 1.67 | Hyb-Seq† | ILLUMINA | Illumina NovaSeq 6000 |

| | | | | |
|---|---|---|---|---|
| 1358033250 | 1.26 | Hyb-Seq† | ILLUMINA | Illumina NovaSeq 6000 |
| 1358188200 | 1.26 | Hyb-Seq† | ILLUMINA | Illumina NovaSeq 6000 |
| 831382050 | 0.77 | Hyb-Seq† | ILLUMINA | Illumina NovaSeq 6000 |
| 284029050 | 0.26 | Hyb-Seq† | ILLUMINA | Illumina NovaSeq 6000 |
| 1156422000 | 1.08 | Hyb-Seq† | ILLUMINA | Illumina NovaSeq 6000 |
| 810048600 | 0.75 | Hyb-Seq† | ILLUMINA | Illumina NovaSeq 6000 |
| 1279652550 | 1.19 | Hyb-Seq† | ILLUMINA | Illumina NovaSeq 6000 |
| 1760767050 | 1.64 | Hyb-Seq† | ILLUMINA | Illumina NovaSeq 6000 |
| 1326305250 | 1.24 | Hyb-Seq† | ILLUMINA | Illumina NovaSeq 6000 |
| 2187473700 | 2.04 | Hyb-Seq† | ILLUMINA | Illumina NovaSeq 6000 |
| 1747543800 | 1.63 | Hyb-Seq† | ILLUMINA | Illumina NovaSeq 6000 |
| 1307905350 | 1.22 | Hyb-Seq† | ILLUMINA | Illumina NovaSeq 6000 |
| 945198150 | 0.88 | Hyb-Seq† | ILLUMINA | Illumina NovaSeq 6000 |
| 786201000 | 0.73 | Hyb-Seq† | ILLUMINA | Illumina NovaSeq 6000 |
| 1104230850 | 1.03 | Hyb-Seq† | ILLUMINA | Illumina NovaSeq 6000 |
| 525151800 | 0.49 | Hyb-Seq† | ILLUMINA | Illumina NovaSeq 6000 |
| 2142889200 | 2.00 | Hyb-Seq† | ILLUMINA | Illumina NovaSeq 6000 |
| 1931898900 | 1.80 | Hyb-Seq† | ILLUMINA | Illumina NovaSeq 6000 |
| 1246627500 | 1.16 | Hyb-Seq† | ILLUMINA | Illumina NovaSeq 6000 |
| 1142067000 | 1.06 | Hyb-Seq† | ILLUMINA | Illumina NovaSeq 6000 |
| 674857200 | 0.63 | Hyb-Seq† | ILLUMINA | Illumina NovaSeq 6000 |
| 974735700 | 0.91 | Hyb-Seq† | ILLUMINA | Illumina NovaSeq 6000 |
| 1050734850 | 0.98 | Hyb-Seq† | ILLUMINA | Illumina NovaSeq 6000 |
| 1664897400 | 1.55 | Hyb-Seq† | ILLUMINA | Illumina NovaSeq 6000 |
| 1345497900 | 1.25 | Hyb-Seq† | ILLUMINA | Illumina NovaSeq 6000 |
| 1188141150 | 1.11 | Hyb-Seq† | ILLUMINA | Illumina NovaSeq 6000 |
| 1281015600 | 1.19 | Hyb-Seq† | ILLUMINA | Illumina NovaSeq 6000 |
| 1423680900 | 1.33 | Hyb-Seq† | ILLUMINA | Illumina NovaSeq 6000 |
| 1745724900 | 1.63 | Hyb-Seq† | ILLUMINA | Illumina NovaSeq 6000 |
| 1444301850 | 1.35 | Hyb-Seq† | ILLUMINA | Illumina NovaSeq 6000 |
| 907402650 | 0.85 | Hyb-Seq†* | ILLUMINA | Illumina NovaSeq 6000 |
| 1674482100 | 1.56 | Hyb-Seq†* | ILLUMINA | Illumina NovaSeq 6000 |
| 3404550600 | 3.17 | RNA_Seq†* | ILLUMINA | Illumina NovaSeq 6000 |
| 3813181650 | 3.55 | RNA_Seq†* | ILLUMINA | Illumina NovaSeq 6000 |
| 3348314850 | 3.12 | RNA-Seq† | ILLUMINA | Illumina NovaSeq 6000 |
| 4075220550 | 3.80 | RNA-Seq† | ILLUMINA | Illumina NovaSeq 6000 |
| 3541990650 | 3.30 | RNA-Seq† | ILLUMINA | Illumina NovaSeq 6000 |
| 3470976900 | 3.23 | RNA-Seq† | ILLUMINA | Illumina NovaSeq 6000 |
| 3143675700 | 2.93 | RNA-Seq† | ILLUMINA | Illumina NovaSeq 6000 |
| 3341026200 | 3.11 | RNA-Seq† | ILLUMINA | Illumina NovaSeq 6000 |
| 63456209700 | 59.10 | WGS | BGISEQ | DNBSEQ-T7 |
| 56404714500 | 52.53 | WGS | BGISEQ | DNBSEQ-T7 |
| 60891719400 | 56.71 | WGS | BGISEQ | DNBSEQ-T7 |
| 45940993200 | 42.79 | WGS | BGISEQ | DNBSEQ-T7 |
| 68112152700 | 63.43 | WGS | BGISEQ | DNBSEQ-T7 |
| 62134697100 | 57.87 | WGS | BGISEQ | DNBSEQ-T7 |
| 41559325500 | 38.71 | WGS | BGISEQ | DNBSEQ-T7 |
| 60801291900 | 56.63 | WGS | BGISEQ | DNBSEQ-T7 |
| 41770309800 | 38.90 | WGS | BGISEQ | DNBSEQ-T7 |
| 37799126700 | 35.20 | WGS | BGISEQ | DNBSEQ-T7 |
| 32381156100 | 30.16 | WGS | ILLUMINA | Illumina HiSeq 2000 |
| 44252260200 | 41.21 | WGS | BGISEQ | DNBSEQ-T7 |

| | | | | |
|---|---|---|---|---|
| 61965302100 | 57.71 | WGS | BGISEQ | DNBSEQ-T7 |
| 51760075200 | 48.21 | WGS | BGISEQ | DNBSEQ-T7 |
| 54263150400 | 50.54 | WGS | BGISEQ | DNBSEQ-T7 |
| 39214039200 | 36.52 | WGS | BGISEQ | DNBSEQ-T7 |
| 19264231800 | 17.94 | WGS | BGISEQ | DNBSEQ-T7 |
| 59400378300 | 55.32 | WGS | BGISEQ | DNBSEQ-T7 |
| 52864167600 | 49.23 | WGS | BGISEQ | DNBSEQ-T7 |
| 52824828600 | 49.20 | WGS | BGISEQ | DNBSEQ-T7 |
| 62861541000 | 58.54 | WGS | BGISEQ | DNBSEQ-T7 |
| 44232388200 | 41.19 | WGS | BGISEQ | DNBSEQ-T7 |
| 12810390300 | 11.93 | WGS | BGISEQ | DNBSEQ-T7 |
| 43270380900 | 40.30 | WGS | BGISEQ | DNBSEQ-T7 |
| 55911981000 | 52.07 | WGS | BGISEQ | DNBSEQ-T7 |
| 52124713500 | 48.54 | WGS | BGISEQ | DNBSEQ-T7 |
| 55415530650 | 51.61 | WGS | ILLUMINA | HiSeq X Ten |
| 69002363400 | 64.26 | WGS | BGISEQ | DNBSEQ-T7 |
| 58869947700 | 54.83 | WGS | BGISEQ | DNBSEQ-T7 |
| 45105787500 | 42.01 | WGS | BGISEQ | DNBSEQ-T7 |
| 41114283000 | 38.29 | WGS | BGISEQ | DNBSEQ-T7 |
| 35308237200 | 32.88 | WGS | BGISEQ | DNBSEQ-T7 |
| 53835731400 | 50.14 | WGS | BGISEQ | DNBSEQ-T7 |
| 56665198800 | 52.77 | WGS | BGISEQ | DNBSEQ-T7 |
| 59512193700 | 55.43 | WGS | BGISEQ | DNBSEQ-T7 |
| 66275746800 | 61.72 | WGS | BGISEQ | DNBSEQ-T7 |
| 35420728950 | 32.99 | WGS | BGISEQ | DNBSEQ-T7 |
| 52630050300 | 49.02 | WGS | BGISEQ | BGISEQ-500 |
| 34372189300 | 32.01 | WGS† | BGISEQ | BGISEQ-500 |
| 36512966000 | 34.01 | WGS† | BGISEQ | BGISEQ-500 |
| 35974572500 | 33.50 | WGS† | BGISEQ | BGISEQ-500 |
| 37495660400 | 34.92 | WGS† | BGISEQ | BGISEQ-500 |
| 40290367800 | 37.52 | WGS† | BGISEQ | BGISEQ-500 |
| 4415735400 | 4.11 | WGS†* | ILLUMINA | Illumina NovaSeq 6000 |
| 3524091450 | 3.28 | WGS†* | ILLUMINA | Illumina NovaSeq 6000 |
| 2720257050 | 2.53 | WGS†* | ILLUMINA | Illumina NovaSeq 6000 |
| 5302303650 | 4.94 | WGS†* | ILLUMINA | Illumina NovaSeq 6000 |

locality
-
Greece
Xizang, China
Chongqing ,China
Greece
Nordjemen, Germany
Italy
Xizang, China
Greece
Xizang, China
Iran
Yunnan, China
-
Hebei, China
Greece
Bulgaria
Syria
Bulgaria
-
France
Bulgaria
Poland
Russian Federation
-
Cyprus
Beijing, China
Sichuan, China
Province Kyeong Sang Nam do, Korea
Greece
Greece
Nebraska, United States
California, United States
Illinois, United States
Xizang, China
Yunnan, China
California, United States
Slovakia
Yunnan, China
Albania
Yunnan, China
Turkey
Italy
California, United States
Bulgaria
Hungary
Greece
Greece
California, United States
England, United Kingdom
Greece
Poland

Switzerland
-
Oregon, United States
Hungary
Hungary
Hungary
Greece
Florida, United States
Colorado, United States
Switzerland
Bulgaria
Xinjiang, China
Hungary
California, United States
California, United States
Korea
Nevada, United States
Yunnan, China
Netherlands
France
Hubei, China
Yunnan, China
Iraq
Switzerland
Poland
Fujian, China
Missouri, United States
Shandong, China
South Africa
South Africa
Gansu, China
Korea
Jilin, China
Sichuan, China
Hebei, China
Hebei, China
Gansu, China
Hebei, China
Sichuan, China
Zhejiang, China
Yunnan, China
Sweden
-
-
-
Hungary
Shanxi, China
Guangxi, China
Bulgaria
Armenia
-
Sweden

Poland
Yunnan, China
-
Romania
-
Massachusetts Hall, Cambridge
-
Massachusetts Hall, Cambridge
Massachusetts Hall, Cambridge
Massachusetts Hall, Cambridge
Massachusetts Hall, Cambridge
Massachusetts Hall, Cambridge
Qinghai, China
Tibet, China
-
Sichuan, China
Tibet, China
Tibet, China
Tibet, China
Yunnan, China
Sichuan, China
South Africa
Australia
Hubei, China
Zhejiang, China
-
Tibet, China
Yunnan, China
Chongqing ,China
Hebei, China
Yunnan, China
Jilin, China
Gansu, China
Korea
Sichuan, China

| voucher | BioProject No. |
|---|---|
| s.coll. # s.n. | PRJNA516111 |
| D. Tzanoudakis et al. # 14723 | PRJNA895940 |
| Q. Wang # BOP018163 | PRJNA895940 |
| Z. Y. Liu # 990246 (PE-01882919) | PRJNA895940 |
| S. Vlachos # 289 | PRJNA895940 |
| D. Podlech # 36461 (PE-01361911) | PRJNA895940 |
| A. Pistarino # 1689 (PE-01361912) | PRJNA895940 |
| W. H. Jin Tian et al. # 103 (PE-01670809) | PRJNA895940 |
| D. Tzanoudakis # 1520 | PRJNA895940 |
| PE Tibet team # 3977 (PE-02056771) | PRJNA895940 |
| J. F. N. Bornmüller et al. # 7610 (US-02708432) | PRJNA895940 |
| Y. M. Shui # 003171 (PE-01372097) | PRJNA895940 |
| Unknown (US-02708408) | PRJNA895940 |
| D. Y. Hong et al. # H03005 (PE-01670820) | PRJNA895940 |
| A. Vaccari (US-02707935) | PRJNA895940 |
| B. Stefanoff (US-02707999) | PRJNA895940 |
| Gaillardot # 110 (US-02708562) | PRJNA895940 |
| L. Kirova (PE-01264238) | PRJNA895940 |
| A. Grossheim et al. (PE-01241292) | PRJNA895940 |
| J. Parnell (PE-01619951) | PRJNA895940 |
| St. Stoyanov (PE-01619953) | PRJNA895940 |
| M. Kozak (PE-01619956) | PRJNA895940 |
| A. Schreter # 11198 (US-02708381) | PRJNA895940 |
| Hohenacker (US-02707760) | PRJNA895940 |
| R. Hand # 5738 (US-01286828) | PRJNA895940 |
| B. Liu # BOP205309 | PRJNA895940 |
| X. H. Jin et al. # 437 (PE-01621959) | PRJNA895940 |
| H. Kim et al. # 0468 (PE-01619938) | PRJNA895940 |
| D. Tzanoudakis et al. # X765 | PRJNA895940 |
| Grebenchikoff (K-001394565) | PRJNA895940 |
| V. A. Sivicek et al. # 45 (PE-01917023) | PRJNA895940 |
| L. Constance et al. # 3045 (US-00788184) | PRJNA895940 |
| L. R. Phillippe # 39747 (PE-01882091) | PRJNA895940 |
| Eco-lab plateau group # 10481 (PE-01479796) | PRJNA895940 |
| Qinghai-Tibet team # 14924 (PE-01173893) | PRJNA895940 |
| J. B. Davy et al. # 6056 (US-00903502) | PRJNA895940 |
| P.Sillinger (PE-01240732) | PRJNA895940 |
| G. Forrest # 11690 (PE-01240762) | PRJNA895940 |
| D. Lakusic et al. # 47200 | PRJNA895940 |
| Kangzang Plant Expedition # 10-3043 (PE-01876206) | PRJNA895940 |
| B. Balansa # 836 (US-02708267) | PRJNA895940 |
| Gibelli (K-001394568) | PRJNA895940 |
| H. L. Mason # 8310 (US-00903719) | PRJNA895940 |
| N. Vihodcevsky (PE-01241130) | PRJNA895940 |
| Pichler (PE-01241133) | PRJNA895940 |
| J. Damboldt. # IV/1 | PRJNA895940 |
| D. Phitos # 8292 | PRJNA895940 |
| J. T. Howell # 21813 (US-00903727) | PRJNA895940 |
| F. C. Stern (US-02707651) | PRJNA895940 |
| D. Phitos et al. # 25528 | PRJNA895940 |
| W. Bartoszek et al. (PE-01505652) | PRJNA895940 |

| | |
|---|---|
| J. R. Chen # 94035 (PE-01361923) | PRJNA895940 |
| Pichler (PE-02022598) | PRJNA895940 |
| R. K. Beattie # 5380 (US-00904068) | PRJNA895940 |
| Unknown (PE-01241389) | PRJNA895940 |
| Porta. (PE-01241650) | PRJNA895940 |
| Rigo. (PE-01241312) | PRJNA895940 |
| C. G. Baenitz (US-02707986) | PRJNA895940 |
| C. van den Hoek et al. # HR0442 (US-02707344) | PRJNA895940 |
| F. R. Fosberg # 60687 (PE-01619942) | PRJNA895940 |
| J. C. Chen # 94188 (PE-01619943) | PRJNA895940 |
| St. Stoyanov (PE-01619944) | PRJNA895940 |
| Z. T. Wang et al. # 345 (PE-02057005) | PRJNA895940 |
| Unknown (PE-01241769) | PRJNA895940 |
| I. L.Wiggins # 14334 (PE-01372344) | PRJNA895940 |
| J. H. Thomas # 1813 (US-02709951) | PRJNA895940 |
| Unknown (PE-01264249) | PRJNA895940 |
| E. C. Twisselmann # 16124 (PE-01372345) | PRJNA895940 |
| Y. M. Shui et al. # 40256 (PE-02057000) | PRJNA895940 |
| s. coll. # 154 (PE-01372432) | PRJNA895940 |
| s. coll. # s.n. (PE-01372426) | PRJNA895940 |
| R. Long et al. # 090135 (PE-02020744) | PRJNA895940 |
| G. P. Yang # 425 (PE-02056588) | PRJNA895940 |
| K. Larsen et al. # 38164 (PE-01361631) | PRJNA895940 |
| J. R. Chen # 94180 (PE-01619969) | PRJNA895940 |
| D. Florczyk (PE-01505495) | PRJNA895940 |
| G. S. He # 15045 (PE-01621244) | PRJNA895940 |
| D. Castaner # 11242 (PE-00868457) | PRJNA895940 |
| C. Y. Guo # 20063-403-3 (PE-01885042) | PRJNA895940 |
| D. Y. Hong # H99002 (PE-01885023) | PRJNA895940 |
| G. Germishuizen # 7081 (PE-01372452) | PRJNA895940 |
| D. Y. Hong et al. # BOP213060 | PRJNA895940 |
| J. Lee (PE-01372393) | PRJNA895940 |
| C. Xu et al. # BOP221163 | PRJNA895940 |
| D. Y. Hong et al. # BOP212789 | PRJNA895940 |
| B. B. Liu et al. # POC588623 | PRJNA895940 |
| B. B. Liu et al. # POC588633 | PRJNA895940 |
| D. Y. Hong et al. # BOP212940 | PRJNA895940 |
| B. B. Liu et al. # POC588595 | PRJNA895940 |
| D. Y. Hong et al. # BOP212794 | PRJNA895940 |
| H. W. Zhang et al. # BOP213399 | PRJNA895940 |
| Jia-Rui Chen # 92532 (PE02056865) | PRJNA895940 |
| Gosta Kjellmert # (PE01917019) | PRJNA895940 |
| Joseph Matera & Francois Malaisse # (PE01361909) | PRJNA895940 |
| L.Smo-ljianinova et Tzvetkova # (PE01241274) | PRJNA895940 |
| A.Kolakovsky et V.Jabrova. # (PE01241310) | PRJNA895940 |
| S.Kupcsok # (PE01241313) | PRJNA895940 |
| T.H.Wang # 260 (PE01241324) | PRJNA895940 |
| Zhi-Song Zhang & Yong-Tian Zhang # 028 (PE01241325) | PRJNA895940 |
| C.Navarro et al. # 4853 (PE01504236) | PRJNA895940 |
| M.Oganesian et al. # 08-0402 (PE01912022) | PRJNA895940 |
| s.coll. # s.n. | PRJNA630544 |
| De-Yuan Hong & Yong-Ming Yuan # H96144 (PE01361954) | PRJNA895940 |

| | |
|---|---|
| H.Trzcinska Tacik # 69 (PE01917020) | PRJNA895940 |
| E-Shen Liu # 22471 (PE01241767) | PRJNA895940 |
| Dr G.C.Van Haesendonck. # 155 (PE01372451) | PRJNA895940 |
| Al.Borza E.Ghisa et P.Pteancu # 3278 (PE01672696) | PRJNA895940 |
| PE01264295 | PRJNA895940 |
| stewart, Nasir # 25594 | PRJNA895940 |
| Reverchon # (PE01241132) | PRJNA895940 |
| A.H.Certis # 5863 | PRJNA895940 |
| R.R.Innes # 898 | PRJNA895940 |
| J.M.Powell # 531 | PRJNA895940 |
| Lundell & Lundell # 13559 | PRJNA895940 |
| S.M.Mcore # 57-184 | PRJNA895940 |
| T.N.Ho B.Bartholomew M.Watson et al # 2276 (PE01670823) | PRJNA895940 |
| FLPH Tibet Expedition # 12-1425 (PE01966840) | PRJNA895940 |
| s.coll. # s.n. | PRJNA627046 |
| Qinghai-Tibet team # 13238 (PE01253542) | PRJNA895940 |
| Qinghai-Tibet team # 12689 (PE01253539) | PRJNA895940 |
| Qinghai-Tibet team # 7402 (PE01253696) | PRJNA895940 |
| De-Yuan Hong et al. # H12010 (PE02056954) | PRJNA895940 |
| Su-Gui Xu # 5285 (PE01253208) | PRJNA895940 |
| Xin-Fen Gao et al. # 3669 (PE01621499) | PRJNA895940 |
| Smook,L. # 10457 (PE01523918) | PRJNA895940 |
| Liang-Qian Li et al. # 280 (PE01882092) | PRJNA895940 |
| Jia-Xin Yang et al. # POC595415 | PRJNA895940 |
| Shi-Liang Zhou # POC524593 | PRJNA895940 |
| s.coll. # s.n. | PRJNA438407 |
| PE- Tibet team # 3977 | PRJNA895940 |
| D. Y. Hong et al. # H10030 (PE-02056914) | PRJNA895940 |
| Z. Z. Liu # 181338 (PE-02056709) | PRJNA895940 |
| B. Liu # BOP205314 | PRJNA895940 |
| X. W. Tian et al. # 903 (PE-01442792) | PRJNA895940 |
| C. Xu et al. # BOP221163 | PRJNA895940 |
| D. Y. Hong et al. # BOP213060 | PRJNA895940 |
| J. Lee (PE-01372393) | PRJNA895940 |
| D. Y. Hong et al. # BOP212789 | PRJNA895940 |

| NCBI accession |
| --- |
| **SRR10026547†** |
| **SRR22197989†** |
| **SRR22197984†** |
| **SRR22197980†** |
| **SRR22197979†** |
| **SRR22197975†** |
| **SRR22197972†** |
| **SRR22197966†** |
| **SRR22197961†** |
| **SRR22197957†** |
| **SRR22197955†** |
| **SRR22197954†** |
| **SRR22197947†** |
| **SRR22197945†** |
| **SRR22197942†** |
| **SRR22197937†** |
| **SRR22197934†** |
| **SRR22197933†** |
| **SRR22197960†** |
| **SRR22197924†** |
| **SRR22197923†** |
| **SRR22197922†** |
| **SRR22197959†** |
| **SRR22197958†** |
| **SRR22197952†** |
| **SRR22197918†** |
| **SRR22197999†** |
| **SRR22197997†** |
| **SRR22197996†** |
| **SRR22197995†** |
| **SRR22197993†** |
| **SRR22197992†** |
| **SRR22197991†** |
| **SRR22197990†** |
| **SRR22197988†** |
| **SRR22197986†** |
| **SRR22197985†** |
| **SRR22197983†** |
| **SRR22197927†** |
| **SRR22197982†** |
| **SRR22197981†** |
| **SRR22197974†** |
| **SRR22197971†** |
| **SRR22197970†** |
| **SRR22197969†** |
| **SRR22197967†** |
| **SRR22197968†** |
| **SRR22197965†** |
| **SRR22197964†** |
| **SRR22197963†** |
| **SRR22197953†** |

**SRR22197950†**
**SRR22197949†**
**SRR22197948†**
**SRR22197946†**
**SRR22197944†**
**SRR22197956†**
**SRR22197943†**
**SRR22197941†**
**SRR22197939†**
**SRR22197938†**
**SRR22197935†**
**SRR22197994†**
**SRR22197932†**
**SRR22197931†**
**SRR22197930†**
**SRR22197928†**
**SRR22197926†**
**SRR22197925†**
**SRR22197921†**
**SRR22197920†**
**SRR22197919†**
**SRR22197917†**
**SRR22197916†**
**SRR22197915†**
**SRR22197914†**
**SRR22197977†**
**SRR22197978†**
**SRR22197913†**
**SRR22197912†**
**SRR22197976†**
**SRR22197940†**
**SRR22198006†**
**SRR22197962†**
**SRR22198001†**
**SRR22198010†**
**SRR22197987†**
**SRR22198008†**
**SRR22198007†**
**SRR22198003†**
**SRR22198002†**
**SRR24743432†**
**SRR24743401†**
**SRR24743403†**
**SRR24743421†**
**SRR24743431†**
**SRR24743400†**
**SRR24743402†**
**SRR24743410†**
**SRR24743428†**
**SRR24743430†**
**SRR11805300†**
**SRR24743399†**

**SRR24743429†**
**SRR24743404†**
**SRR24743409†**
**SRR24743427†**
**SRR24743426†**
**SRR24743424†**
**SRR24743433†**
**SRR24743423†**
**SRR24743420†**
**SRR24743422†**
**SRR24743419†**
**SRR24743425†**
**SRR24743418†**
**SRR24743417†**
**SRR11585868†**
**SRR24743415†**
**SRR24743414†**
**SRR24743413†**
**SRR24743412†**
**SRR24743416†**
**SRR24743411†**
**SRR24743407†**
**SRR24743408†**
**SRR24743406†**
SRR24743405†
SRR7121795
**SRR22197936†**
**SRR22197998†**
**SRR22197973†**
**SRR22198004†**
**SRR22198009†**
**SRR22197951†**
**SRR22197929†**
**SRR22198005†**
**SRR22198000†**
