## Supplementary material for "Dense Sampling of Taxa and Genomes Untangles the Phylogenetic Backbone of a Non-model Plant Lineage Rife with Deep Hybridization and Allopolyploidy": Table S2

| species_recognized in this study | Number of genes assembled by HybPiper |
| --- | --- |
| *Adenophora stenanthina* subsp. *sylvatica* | 519 |
| *Adenophora potaninii* subsp. *wawreana* | 545 |
| *Adenophora divaricata* | 550 |
| *Adenophora delavayi* | 556 |
| *Campanula aparinoides* | 561 |
| *Adenophora capillaris* subsp. *paniculata* | 583 |
| *Adenophora stricta* | 585 |
| *Adenophora potaninii* | 611 |
| *Campanula garganica* subsp. *cephallenica* | 612 |
| *Campanula pallida* | 618 |
| *Adenophora palustris* | 620 |
| *Adenophora triphylla* | 621 |
| *Campanula erinus* | 626 |
| *Campanula scheuchzeri* | 627 |
| *Campanula patula* | 631 |
| *Heterocodon rariflorus* | 631 |
| *Campanula sparsa* | 632 |
| *Campanula comosiformis* | 636 |
| *Campanula punctata* | 636 |
| *Homocodon brevipes* | 636 |
| *Jasione montana* | 636 |
| *Platycodon grandiflorus* | 636 |
| *Triodanis perfoliata* | 636 |
| *Cyananthus delavayi* | 640 |
| *Roella decurrens* | 640 |
| *Campanula glomerata* | 641 |
| *Jasione heldreichii* 2 | 641 |
| *Phyteuma orbiculare* | 641 |
| *Campanula hawkinsiana* 2 | 642 |
| *Campanula rotundifolia* | 642 |
| *Musschia peregrina* | 642 |
| *Cyananthus hookeri* | 643 |
| *Lobelia chinensis* | 643 |
| *Campanula aizoides* | 644 |
| *Campanula garganica* | 644 |
| *Campanula ramosissima* 2 | 644 |
| *Campanula rhodensis* | 644 |
| *Campanula stellaris* | 644 |
| *Hesperocodon hederaceus* | 644 |
| *Legousia speculum-veneris* | 644 |
| *Adenophora polyantha* | 645 |
| *Campanula americana* | 645 |
| *Campanula chinensis* | 645 |
| *Campanula portenschlagiana* | 645 |
| *Codonopsis lanceolata* | 645 |
| *Cyananthus formosus* | 645 |
| *Jasione crispa* | 645 |
| *Adenophora remotiflora* | 646 |
| *Asyneuma chinense* | 646 |
| *Campanula asperuloides* | 646 |
| *Campanula californica* | 646 |

*Campanula dimorphantha* 646
*Campanula elatines* 646
*Campanula fenestrellata* 646
*Codonopsis pilosula* 646
*Jasione laevis* 646
*Adenophora petiolata* 647
*Campanula cymbalaria* 647
*Campanula robinsiae* 647
*Codonopsis canescens* 647
*Musschia lactiflora* 1 647
*Theodorovia karakuschensis* 647
*Wahlenbergia undulata* 647
*Campanula edulis* 648
*Campanula persicifolia* 648
*Campanula propinqua* 648
*Campanula uniflora* 648
*Githopsis specularioides* 648
*Lobelia seguinii* 648
*Adenophora longipedicellata* 649
*Adenophora stenanthina* 649
*Campanula angustiflora* 649
*Campanula crenulata* 649
*Campanula scutellata* 649
*Campanula takesimana* 649
*Campanula trachelium* 649
*Codonopsis foetens* 649
*Musschia lactiflora* 2 649
*Turcocodon albertii* 649
*Adenophora coelestis* 650
*Campanula calcicola* 650
*Campanula cervicaria* 650
*Campanula macrostachya* 650
*Campanula medium* 650
*Campanula rainerii* 650
*Githopsis pulchella* 650
*Jasione heldreichii* 1 650
*Legousia hybrida* 650
*Peracarpa carnosa* 2 650
*Phyteuma spicatum* 650
*Triodanis coloradoensis* 650
*Triodanis lamprosperma* 650
*Asyneuma japonicum* 651
*Asyneuma thomsonii* 651
*Campanula carpatica* 651
*Campanula dichotoma* 651
*Campanula floridana* 651
*Campanula latifolia* 651
*Campanula mekongensis* 651
*Pankycodon purpureus* 651
*Petromarula pinnata* 651
*Pseudocodon rosulatus* 651
*Triodanis perfoliata* subsp. *biflora* 651

*Wahlenbergia capensis* 651
*Campanula aizoon* 652
*Campanula aristata* 652
*Campanula cana* 652
*Campanula chrysosplenifolia* 652
*Campanula exigua* 652
*Campanula griffinii* 652
*Campanula incanescens* 652
*Campanula lourica* 652
*Campanula orphanidea* 652
*Campanula reverchonii* 652
*Campanula yunnanensis* 652
*Edraianthus serbicus* 652
*Wahlenbergia marginata* 652
*Campanula expansa* 653
*Campanula jacquinii* 653
*Campanula longistyla* 653
*Campanula prenanthoides* 653
*Campanula pulla* 653
*Campanula rapunculoides* 653
*Campanula thyrsoides* 653
*Cyclocodon lancifolius* 653
*Favratia zoysii* 653
*Peracarpa carnosa* 1 653
*Wahlenbergia androsacea* 653
*Campanula* 654
*Campanula drabifolia* 654
*Campanula fastigiata* 654
*Campanula hawkinsiana* 1 654
*Campanula ramosissima* 1 654
*Hanabusaya latisepala* 654

| Number of genes with paralog warnings | Number of 1to1 orthologs | Number of MO orthologs |
|---|---|---|
| 567 | 374 | 499 |
| 577 | 392 | 519 |
| 589 | 398 | 521 |
| 564 | 378 | 525 |
| 563 | 380 | 543 |
| 632 | 417 | 557 |
| 636 | 420 | 559 |
| 672 | 428 | 592 |
| 646 | 442 | 627 |
| 631 | 418 | 592 |
| 676 | 430 | 600 |
| 664 | 432 | 596 |
| 634 | 430 | 596 |
| 636 | 424 | 589 |
| 646 | 426 | 599 |
| 642 | 429 | 607 |
| 647 | 428 | 606 |
| 656 | 441 | 628 |
| 655 | 434 | 598 |
| 688 | 424 | 583 |
| 669 | 434 | 600 |
| 679 | 437 | 625 |
| 666 | 435 | 611 |
| 674 | 438 | 632 |
| 653 | 436 | 612 |
| 659 | 441 | 614 |
| 716 | 434 | 603 |
| 653 | 439 | 623 |
| 649 | 440 | 627 |
| 650 | 440 | 628 |
| 670 | 441 | 623 |
| 674 | 441 | 635 |
| 649 | 413 | 602 |
| 664 | 441 | 627 |
| 655 | 445 | 632 |
| 650 | 441 | 627 |
| 644 | 441 | 616 |
| 644 | 438 | 617 |
| 653 | 430 | 595 |
| 645 | 439 | 610 |
| 650 | 446 | 623 |
| 699 | 438 | 615 |
| 646 | 440 | 623 |
| 655 | 429 | 608 |
| 670 | 442 | 635 |
| 674 | 442 | 637 |
| 660 | 438 | 609 |
| 650 | 446 | 622 |
| 655 | 447 | 635 |
| 663 | 440 | 623 |
| 652 | 443 | 624 |

| | | |
|---|---|---|
| 649 | 441 | 621 |
| 654 | 445 | 633 |
| 626 | 416 | 585 |
| 674 | 438 | 632 |
| 650 | 446 | 611 |
| 653 | 445 | 625 |
| 657 | 443 | 630 |
| 654 | 443 | 632 |
| 670 | 443 | 638 |
| 675 | 444 | 627 |
| 649 | 440 | 614 |
| 644 | 442 | 613 |
| 656 | 444 | 625 |
| 660 | 444 | 632 |
| 648 | 442 | 617 |
| 672 | 446 | 628 |
| 650 | 444 | 623 |
| 651 | 438 | 632 |
| 664 | 446 | 623 |
| 668 | 446 | 624 |
| 650 | 444 | 624 |
| 648 | 444 | 621 |
| 655 | 443 | 625 |
| 647 | 443 | 624 |
| 669 | 443 | 620 |
| 673 | 445 | 641 |
| 658 | 445 | 621 |
| 660 | 444 | 634 |
| 670 | 447 | 627 |
| 653 | 446 | 626 |
| 670 | 445 | 626 |
| 647 | 445 | 622 |
| 645 | 445 | 624 |
| 659 | 447 | 638 |
| 653 | 446 | 627 |
| 688 | 444 | 614 |
| 650 | 446 | 619 |
| 644 | 447 | 634 |
| 653 | 446 | 637 |
| 646 | 448 | 601 |
| 655 | 445 | 615 |
| 657 | 447 | 633 |
| 652 | 445 | 629 |
| 652 | 446 | 638 |
| 647 | 445 | 629 |
| 657 | 445 | 629 |
| 658 | 445 | 624 |
| 644 | 445 | 622 |
| 671 | 447 | 644 |
| 656 | 447 | 633 |
| 647 | 449 | 644 |
| 668 | 446 | 621 |

| | | |
|---|---|---|
| 656 | 448 | 623 |
| 655 | 448 | 636 |
| 650 | 447 | 625 |
| 655 | 447 | 628 |
| 685 | 448 | 627 |
| 654 | 446 | 631 |
| 654 | 445 | 630 |
| 654 | 447 | 627 |
| 658 | 443 | 622 |
| 648 | 447 | 629 |
| 663 | 445 | 625 |
| 665 | 447 | 629 |
| 655 | 446 | 621 |
| 644 | 447 | 623 |
| 658 | 448 | 633 |
| 658 | 448 | 626 |
| 663 | 447 | 628 |
| 654 | 448 | 626 |
| 653 | 447 | 641 |
| 662 | 447 | 629 |
| 654 | 447 | 627 |
| 690 | 449 | 646 |
| 650 | 448 | 638 |
| 652 | 448 | 638 |
| 656 | 447 | 625 |
| 650 | 448 | 621 |
| 656 | 448 | 628 |
| 636 | 444 | 627 |
| 654 | 448 | 640 |
| 655 | 448 | 637 |
| 646 | 449 | 628 |

| Number of RT orthologs |
|---|
| 504 |
| 520 |
| 526 |
| 527 |
| 542 |
| 566 |
| 567 |
| 599 |
| 627 |
| 600 |
| 605 |
| 600 |
| 603 |
| 592 |
| 600 |
| 612 |
| 609 |
| 629 |
| 606 |
| 587 |
| 603 |
| 620 |
| 608 |
| 626 |
| 610 |
| 621 |
| 606 |
| 623 |
| 626 |
| 627 |
| 629 |
| 629 |
| 616 |
| 630 |
| 635 |
| 627 |
| 618 |
| 619 |
| 594 |
| 609 |
| 623 |
| 612 |
| 626 |
| 613 |
| 628 |
| 631 |
| 613 |
| 622 |
| 633 |
| 629 |
| 625 |

625
634
590
624
613
626
631
633
632
633
617
612
632
637
620
635
623
650
627
629
623
623
632
633
629
634
623
636
632
622
638
633
634
639
628
616
617
638
633
602
613
631
632
634
638
633
629
629
637
631
637
617

622
634
623
638
624
631
630
637
628
637
635
639
629
622
630
630
638
626
637
640
638
639
636
640
624
624
631
625
636
634
626
